# CRISPR-Cas9-mediated editing of *myb28* impairs glucoraphanin accumulation of *Brassica oleracea* in the field

**DOI:** 10.1101/2020.07.16.206813

**Authors:** Mikhaela Neequaye, Sophia Stavnstrup, Tom Lawrenson, Penny Hundleby, Perla Troncoso-Rey, Shikha Saha, Wendy Harwood, Maria H Traka, Richard Mithen, Lars Østergaard

## Abstract

We sought to quantify the role of *MYB28* in the regulation of aliphatic glucosinolate biosynthesis and associated sulphur metabolism in field-grown *B. oleracea* with the use of CRISPR-Cas9-mediated gene editing technology. We describe the first characterised *myb28* knockout mutant in *B. oleracea*, and the first UK field trial of CRISPR-Cas9-mediated gene edited plants under the European Court of Justice interpretation of the 2001/18 EU GMO directive. We report that knocking-out *myb28* results in downregulation of aliphatic glucosinolate biosynthesis genes and reduction in accumulation of the methionine-derived glucosinolate, glucoraphanin, in leaves and florets of field-grown *myb28* mutant broccoli plants. There were no significant changes to the accumulation of sulphate, S-methyl cysteine sulfoxide and indole glucosinolate in leaf and floret tissues.

## Introduction

Diets rich in cruciferous vegetables are associated with beneficial effects upon human health. This may be due to the activity of the degradation products of thioglucosides known as glucosinolates that accumulate in these vegetables. These compounds comprise a common glycine moiety and a variable side chain derived from a small number of amino acids (Halkier and Gershenzon, 2006). Upon consumption, glucosinolates with an aliphatic side chain derived from methionine are largely converted to isothiocyanates, whereas glucosinolates with an indole side chain derived from tryptophan and hydrolysed to a complex series of indole compounds (Mithen, 2001). Aliphatic and indole glucosinolate derivatives may both contribute to the putative health-promoting effects of cruciferous vegetables (Juge et al., 2007). Of these diverse compounds, the isothiocyanate sulforaphane derived from 4-methylsulphinylbutyl glucosinolate glucoraphanin, which accumulates in broccoli, *Brassica oleracea* var. *italica*, has received by far the greatest attention. Within the sulphur metabolome of cruciferous vegetables, glucosinolates accumulate alongside S-methyl cysteine sulfoxide (SMCSO) which also degrades to bioactive metabolites (Edmands et al., 2011; Traka et al., 2013). Following the uptake of inorganic sulphate from the roots, vacuolar sulphate forms the vital precursor to these bioactives (Kopriva, 2006). Many environmental factors influence the glucosinolate profile and sulphur metabolome of *Brassica* crops, including climate, soil content, exposure to pathogens and insect pests. Nonetheless, the most important factor in determining the glucosinolate profile and sulphur metabolome of a crop is genotype, though contributions of regulatory genes is yet to be fully understood.

The R2R3 MYB transcription factor MYB28 has been characterised as a key regulator of aliphatic glucosinolate biosynthesis. This includes direct regulation of expression of glucosinolate biosynthesis genes involved in every step of glucosinolate production (Gigolashvili et al., 2007; Hirai et al., 2007; Sonderby et al., 2007) in addition to regulating primary sulphur assimilation genes and partitioning of methionine, vital precursors for aliphatic glucosinolates (Frerigmann and Gigolashvili, 2014; Sonderby et al., 2007; Yatusevich et al., 2010). This regulatory role is conserved within the *Brassica* genus, with homologues characterised in *Brassica napus* (Li et al., 2014), *B. juncea* (Augustine et al., 2013) and *B. rapa* (Kim et al., 2013; Seo et al., 2016).This includes studies to a lesser extent in *B. oleracea* varieties kohlrabi (Yi et al., 2015) and Chinese kale (Yin et al., 2017), which generated RNAi lines to study the function of this transcription factor. To date no stable *MYB28* knockout has been published in this species. Single-gene editing in *Brassica* species can be complicated by the historical whole genome triplication, resulting in multiple copies of numerous genes, including *MYB28* (Cheng et al., 2014). Within the *B. oleracea* genome, *MYB28* exists in three copies; one on chromosome 2 (Bo2g161590), chromosome 9 (Bo9g014610) and chromosome C7 (Bo7g098590). This is further complicated by two additional copies of the close homologue, *MYB29*, on chromosome 3 (Bo3g004500) and chromosome 9 (Bo9g175680), which is known to work alongside MYB28 in regulating aliphatic glucosinolate biosynthesis (Gigolashvili et al., 2007; Hirai et al., 2007; Sonderby et al., 2007). The expression of the C2 copy is thought to be of most importance in regulating aliphatic glucosinolate biosynthesis in aerial organs (Kittipol et al., 2019).

This work aims to further characterise the role *MYB28* plays in the regulation of aliphatic glucosinolate biosynthesis and sulphur metabolism in field-grown *B. oleracea* using CRISPR-Cas9 mediated gene editing technology. This work describes the first characterised *myb28* knockout mutant in a *B. oleracea* background and the subsequent effect on the transcriptome of the crop through RNAseq analysis. We describe the effect of knocking out *myb28* function in regulating aliphatic glucosinolate biosynthesis within the context of the wider sulphur metabolome, with targeted metabolomic analysis of primary sulphur metabolites, sulphate and s-methyl cysteine sulfoxide in addition to indolic glucosinolates. In 2018 the European Court of Justice (ECJ) ruled that gene-edited crops such as those developed in the current study would be considered as genetically modified organisms (GMOs) and would not be exempt from the 2001/18 EU GMO Directive, in the same way as crops derived from older forms of mutagenesis (Hundleby and Harwood, 2019). Prior to this ruling the first UK CRISPR field trial (of edited Camelina) had been initiated and approved by the Department of Environment, Food and Rural Affairs (DEFRA) free from the GMO restrictions set out by this ECJ ruling in early 2018 (Faure and Napier, 2018). The research described in this publication involved the first DEFRA-regulated field experiment with CRISPR-Cas9 edited plants under the ECJ interpretation of the 2001/18 EU GMO directive (Callaway, 2018).

## Results

### CRISPR-Cas9 mediated editing of *myb28* in *B. oleracea*

In order to characterise the role of *MYB28* in *B. oleracea* a series of *myb28* mutant lines were generated using CRISPR-Cas9-mediated gene-editing technology. Guides designed to target the three copies of *MYB28* found in the *B. oleracea* genome were introduced to 4-day old cotyledonary explants of the *Brassica oleracea* line DH1012 by *Agrobacterium-mediated* transformation as described in (Lawrenson et al., 2019; Lawrenson et al., 2015). This encompassed two constructs each containing two guides, with four guides in total (see Figure S1 for guide sequences and design). A series of T0 plants were screened by amplifying and sequencing the three copies of *MYB28* to look for characteristic Cas9-editing activity. No homozygous mutations were found in the T0 generation. Of the 192 subsequent T1 seedlings screened, 9 showed editing in *MYB28* C2 with an additional 12 showing editing at *MYB28* C9. These plants were derived from those transformed using the construct referred to as ‘1986’ which contained the sequences for guide 59 and guide 60. The T2 generation gave rise to plants with homozygous mutations, in which two types of homozygous mutation in the C2 copy of *MYB28* were found to be segregating within the population. One of which is a single base pair insertion, a thymine, leading to a frameshift mutation in which a premature stop codon is introduced. Lines with this mutation were designated as *bolC2.myb28-1^ge^* (See Figure 1a-b and Figures S2-S3). The second contained a mutation appearing at the same point in the gene sequence producing a three base pair deletion, leading to the deletion a single leucine residue in the resulting protein. Lines with this mutation were designated as *bolC2.myb28-2^ge^* (See Figure 1c-d and Figures S4-S5). Both of these alleles have a homozygous 562 base pair deletion in the C9 copy of *MYB28* likely to render the resulting protein non-functional (Figure S6-S7). All lines carry the wild-type C7 copy of *MYB28* as well as all copies of the most closely-related MYB transcription factor, *MYB29*.

**Figure 1.**
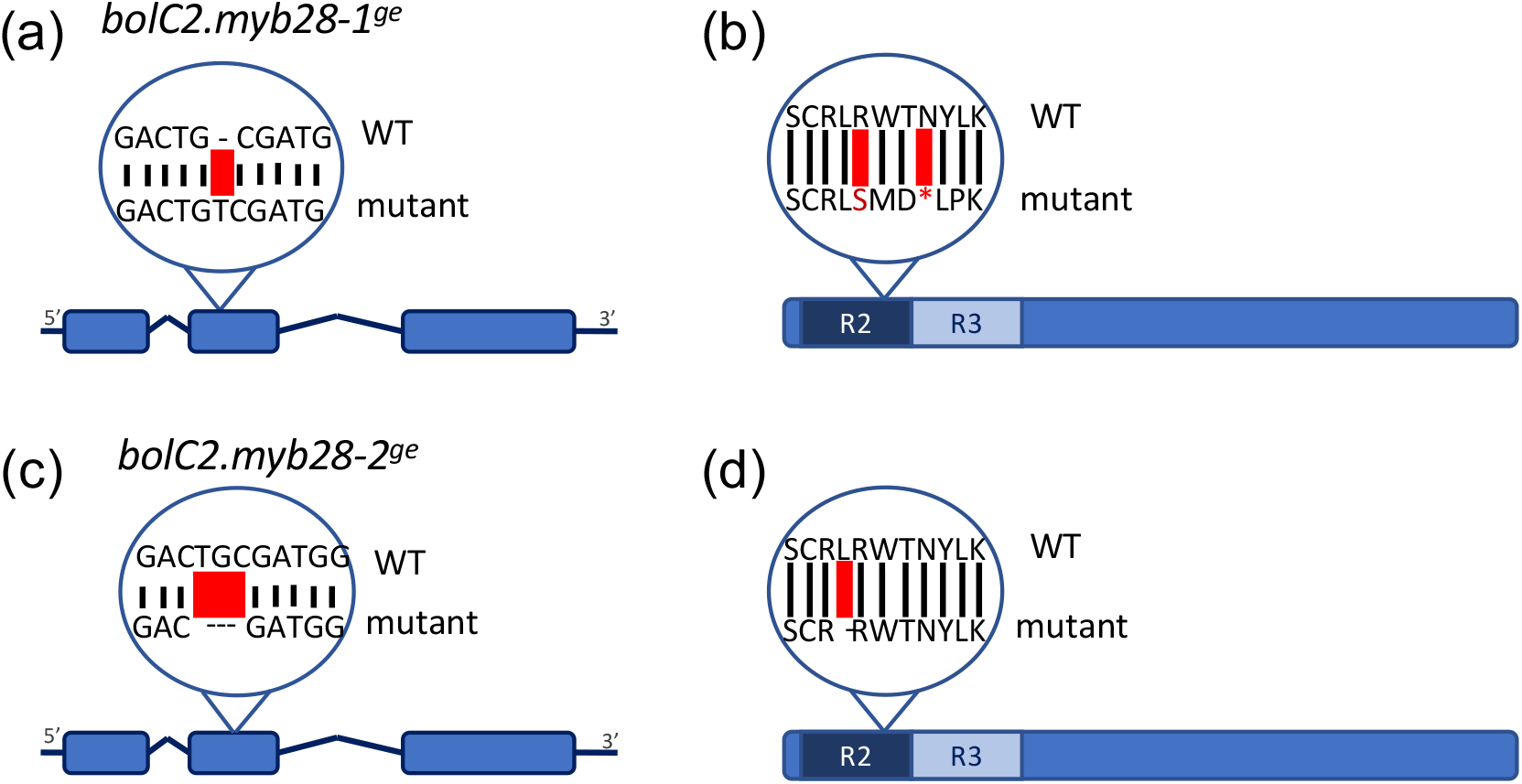
Description of *B. oleracea myb28* mutant alleles generated by CRISPR-Cas9. (a) *MYB28* C2 gene sequence with thymine (T) insertion in mutant allele, *bolC2.myb28^ge-1^*. (b) Resulting protein sequence of MYB28C2 with premature stop codon in the R2 domain as a result of thymine insertion in *bolC2.myb28^ge-1^*. (c) *MYB28C2* gene sequence with a 3-base pair deletion in mutant allele *bolC2.myb28^ge-2^*. (d) Resulting protein sequence of MYB28C2 with leucine deletion in R2 domain in *bolC2.myb28^ge-2^*. Wild type (WT) and mutant sequences are indicated.

### Transgene screening and field-trial layout

In order to confirm the absence of the Cas9-encoding transgene, components of the four cassettes, including the two antibiotic selection markers of NPTII and SPEC as well as the *Cas9* gene and U626 promoter, were analysed for potential presence in individual from the T2 generation of these lines (Lawrenson et al., 2019; Lawrenson et al., 2015). This analysis comprised of PCR screening for the presence of SPEC and U626 (Figure S8 and Figure S9) and copy-number screening by qPCR of the Cas9 and NPTII (Table S1 and Table S2). None of these components were found in the wild-type or homozygous *bolC2.myb28* mutant plants that were transferred to the field.

Four-week old plantlets were transferred to the field in a randomised design (Figure 2) with a border of *B. napus* plantlets. The trial included 10 *bolC2.myb28-1^ge^* plants, 10 *bolC2.myb28-2^ge^* plants, 10 *B. oleracea* 1012 wild type plants and 6 T2 plants that scored as wild-type when genotyping for the *MYB28* mutations (Figure 2a). Due to damage, we were unable to carry out the analyses on all plants but obtained sufficient material for biological triplicates for the analyses presented.

**Figure 2.**
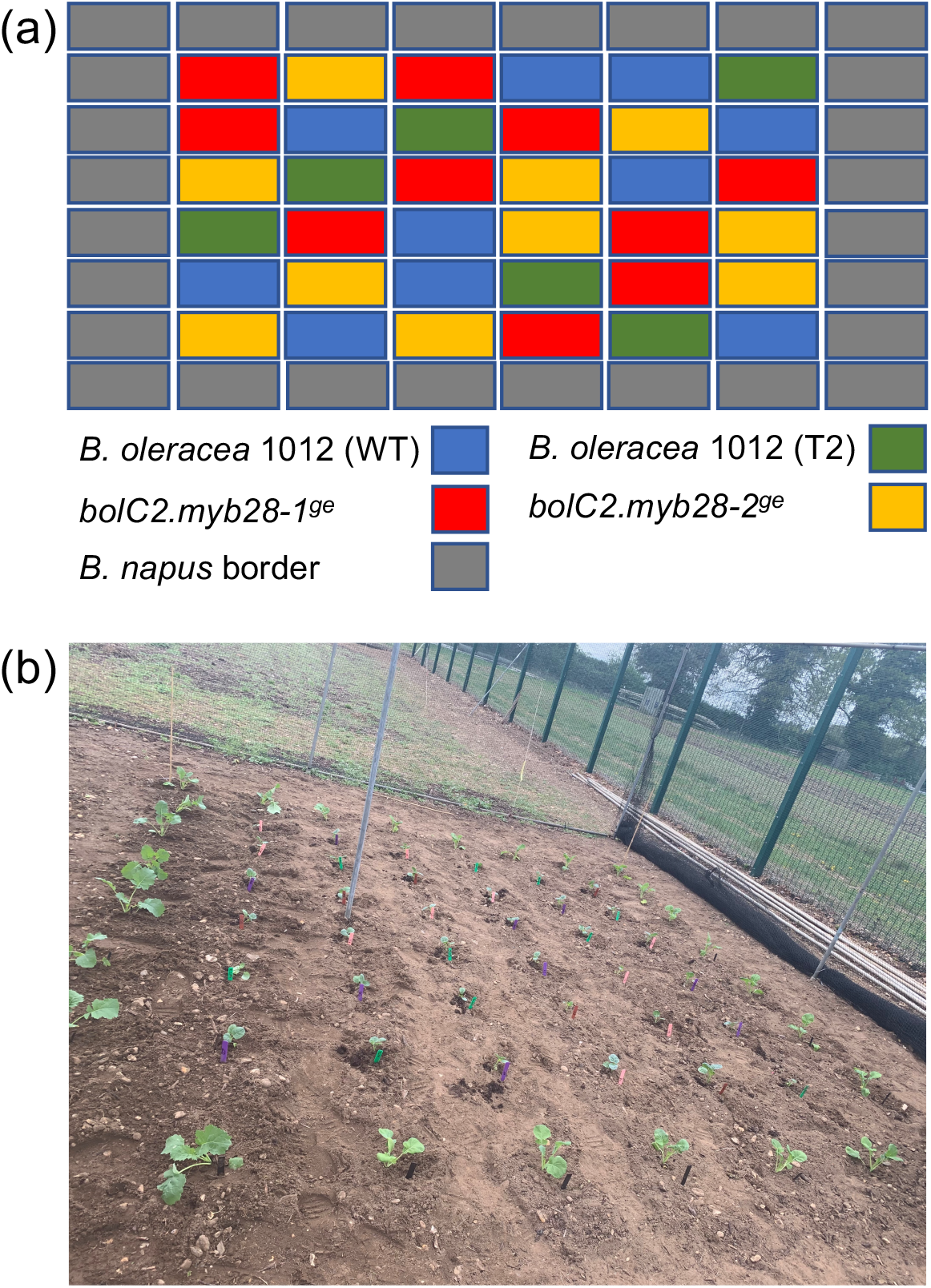
Field-trial layout. (a) Schematic layout with colour-coding representing each genotype according to scheme below. (b) Photo of field plot taken on the day of seedling transplantation. The bigger seedlings are the *B. napus* border plants.

### Gene expression changes in *bolC2.myb28* mutants grown in the field

In order to characterise the changes in global gene expression that occur as a result of knocking out this transcription factor, an RNA-Seq analysis was performed on vegetative leaves, generative leaves and florets of the *bolC2.myb28* mutants grown under field conditions (see Table S3 for details of sequenced material). From this, the vegetative leaves of both mutant lines were found to have 121 differentially expressed genes compared to the wild-type T2, 17 of which were uniquely downregulated in this tissue (Table S4), while generative leaves had 196, in which 55 of were unique to this tissue (Table S5). Florets of the mutant lines had 133 significantly differentially expressed genes, 28 of which were unique to this tissue (Table S6). Of the 27 significantly down-regulated genes (p<0.05) in all tissues were those involved in sulphur metabolism such as *ADENOSINE-5’-PHOSPHOSULFATE (APS) KINASE 2 (APK2), HOMOCYSTEINE S-METHYLTRANSFERASE 3 (HMT3)* and *MRSA2*, which is involved in L-methionine thioredoxin-disulfide S-oxidoreductase activity. Downregulated genes in all tissues of the *myb28* mutants also include glucosinolate biosynthesis genes *BCAT4, BAT5, MAM3, IPMI, IMDH3* of the first step of glucosinolate biosynthesis, methionine elongation. Of the second step of this pathway, core structure formation, *CYP79F1, CYP83A1, SUR1, UGT74C1, GSTF11, SOT17* and *SOT18* are also found to be downregulated in all analysed tissues of the field-grown *myb28* mutant lines (Figure 3). Interestingly glucosinolate and sulphur metabolism genes such as the flavin monooxygenase *FMO-GSOX2*, methylthioalkylmalate isomerase *IIL1* and methionine synthase, *MS2* are found only significantly downregulated in the vegetative leaf tissues. The glutathione transferase *GSTU23* and UDP glycosyltranferase *UGT73B5* are found to be significantly downregulated in both leaf tissue types but not the florets. The degree of reduction in gene expression is comparable in both mutant lines. Moreover, most significantly differentially expressed genes appear to be involved in metabolism and defence responses with little variation in other pathways including no detected differences in expression of MYB or MYC transcription factors, which are often implicated in regulation of glucosinolate biosynthesis. This is further supported in a Gene Ontology (GO) enrichment analysis of these differentially expressed genes as terms associated with glucosinolate metabolic processes, amino acid and sulphur metabolism are amongst those most significantly enriched in all tissues (Table S7-S9). Additional enriched pathways also include hormone responses such as auxin metabolic processes in addition to biotic and abiotic stress responses.

**Figure 3.**
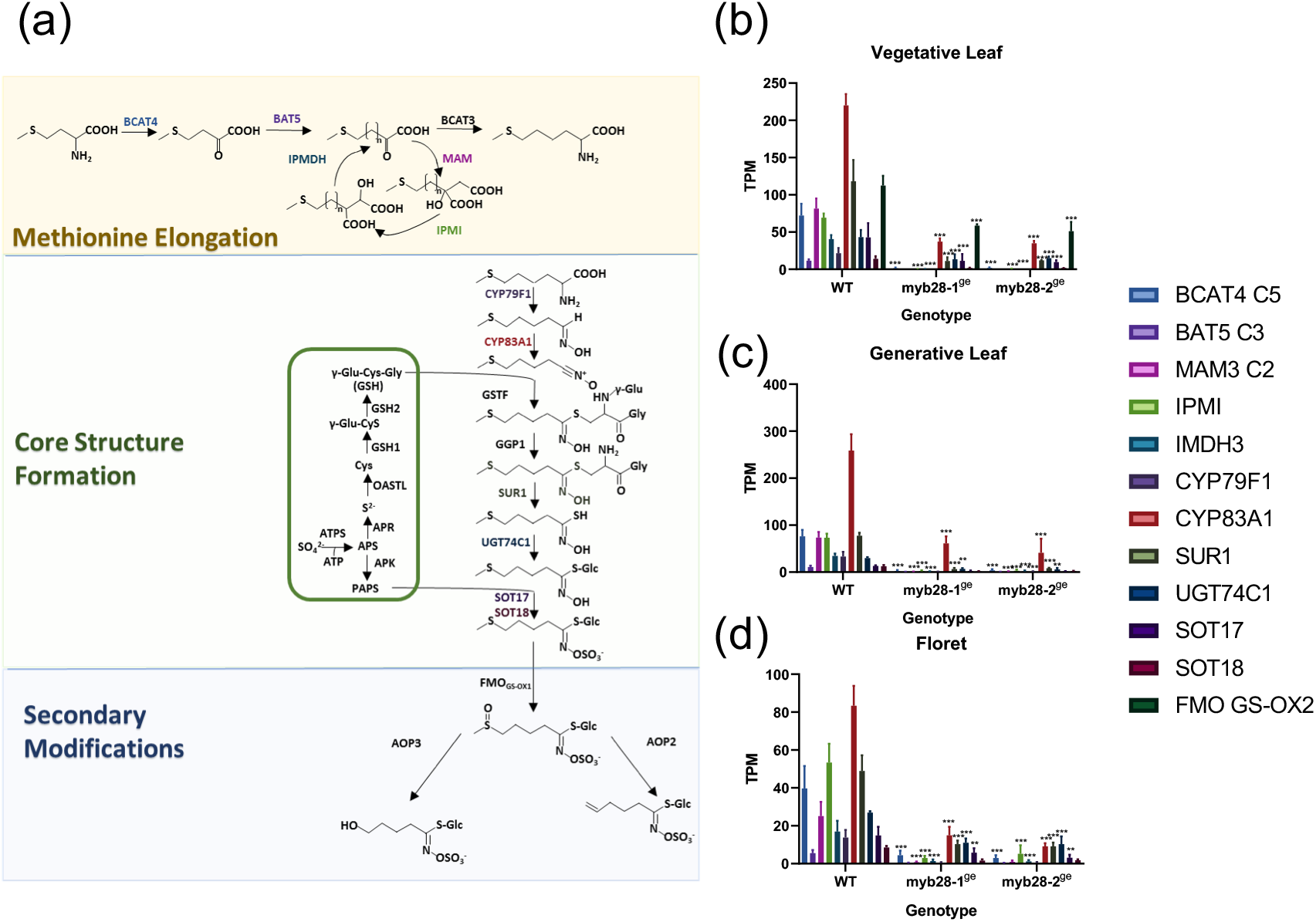
The effect of *myb28* loss-of-function on sulphur metabolism in field-grown *B. oleracea*. (a) Glucoraphanin biosynthesis pathway. Genes responsible for each stage in the biosynthesis pathway alongside each arrow, as colour coded. The first stage, methionine elongation, is highlighted in the yellow box. The second stage, core structure formation, is highlighted in the green box. Within the second step, the smaller box includes the sulphate assimilatory pathway, which provides the additional substrates for biosynthesis, GSH and PAPS. The final stage, secondary modification, is highlighted in the blue box. Abbreviations: APR, APS reductase; APS, adenosine-5’-phosphosulfate; OASTL, O-acetylserine(thiol)lyase; PAPS, 3’-phosphoadenosine-5’-phosphosulfate. Adapted from Edited from Biosynthesis of glucosinolates – gene discovery and beyond (Sonderby et al., 2010). (b) Downregulated genes in glucoraphanin biosynthesis in *myb28* mutants in Vegetative Leaves. (c) Downregulated genes in glucoraphanin biosynthesis in *myb28* mutants in Generative Leaves. (d) Downregulated genes in glucoraphanin biosynthesis in *myb28* mutants in Florets. WT = T2 progeny of transgenic CRISPR line with no presence of transgene and wild-type *MYB28* sequences; *bolC2.myb28-1^g^; bolC2.myb28-2^ge^*. Both lines have a large deletion in C9 *MYB28* leading to a non-functional protein. Mean values plotted for 3 biological replicates each of WT, *bolC2.myb28-1^g^, bolC2.myb28-2^ge^*, respectively. Standard deviation is shown as error bars. Significance calculated using Tukey’s multiple comparison of mutant lines against WT T2, * = p<0.05, ** = p<0.01, *** = p<0.001.

### Sulphur metabolite content of *myb28* mutants in the field

Next, we investigated whether the effect on gene expression led to changes in metabolite profiles. To this end, HPLC analysis was used to quantify metabolite levels in field-grown *B. oleracea myb28* mutants. The methionine-derived aliphatic glucosinolates glucoiberin (3-MSP) and glucoraphanin (4-MSB) were found to be altered in these mutants (Figure 4). In the vegetative stage leaves a significant reduction was found only in glucoraphanin with glucoiberin levels not significantly changed (Figure 4a). This trend became more pronounced in tissues of a later developmental stage in which the florets and generative leaves show a significant reduction in glucoraphanin as well as a significant, but less pronounced, reduction in glucoiberin (Figure 4b-c).

**Figure 4.**
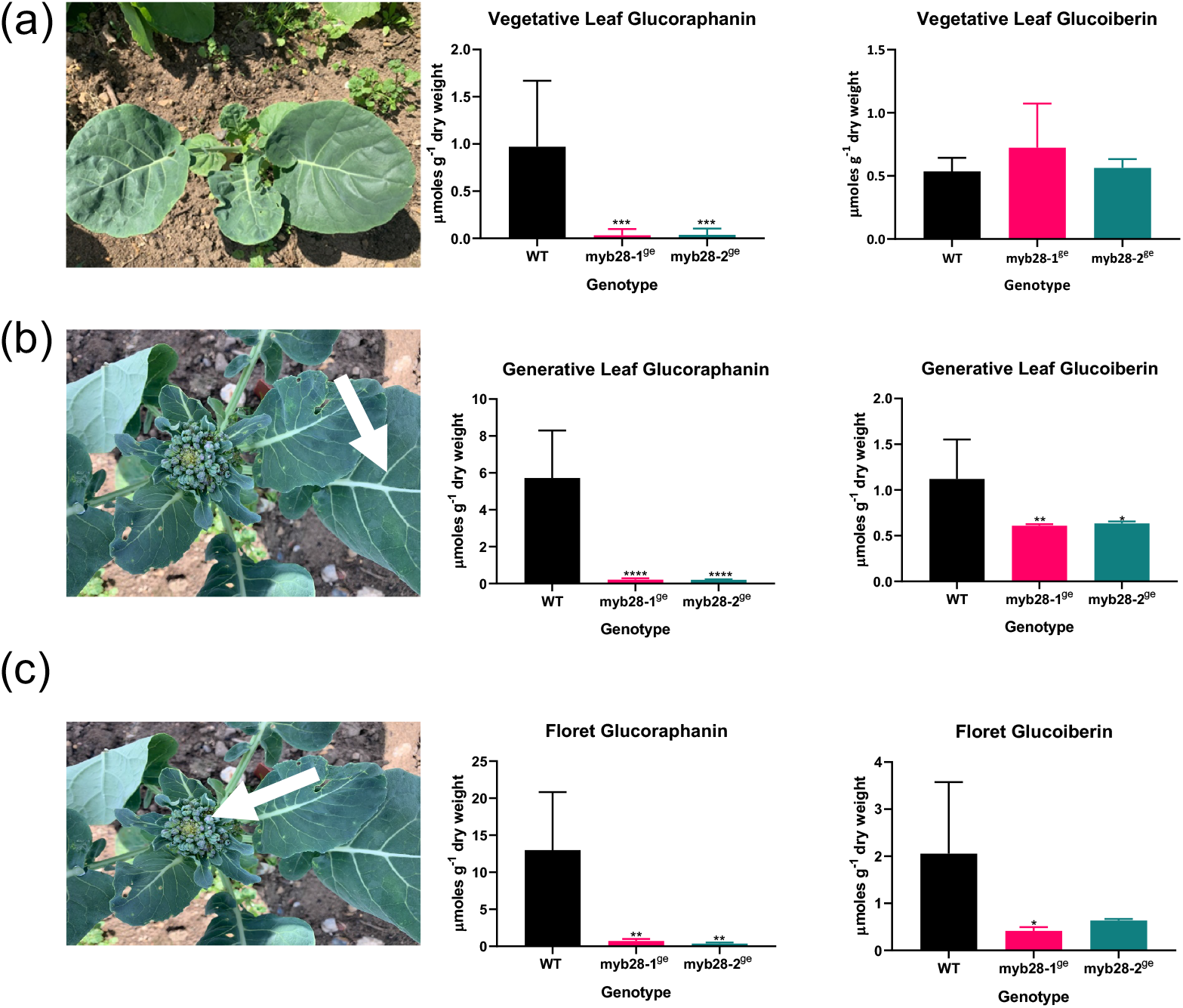
Aliphatic glucosinolate levels of glucoraphanin and glucoiberin in (a) vegetative leaves, (b) generative leaves and (c) florets of field-grown *bolC2.myb28* mutant lines. WT = T2 progeny of transgenic CRISPR line with no presence of transgene and wild-type *MYB28* sequences. *bolC2.myb28-1^ge^; bolC2.myb28-2^ge^*. Mean values plotted for 4 biological replicates each of WT, *bolC2.myb28-1^g^, bolC2.myb28-2^ge^*, respectively. Arrows in (b) and (c) indicate the harvested tissue. Standard deviation is shown as error bars. Significance calculated using Tukey’s multiple comparison of mutant lines against WT T2, * = p<0.05, ** = p<0.01, *** = p<0.001. (See Tables S10-S12 for a summary of statistical analyses)

In order to identify whether the effect on metabolism was exclusive to methionine-derived glucosinolates, additional major sulphur pools were also analysed for disruption in the *myb28* mutants. This included analysis of the indolic glucosinolate biosynthesis pathway, derived from tryptophan, in which no significant variation was detected between the wild type and two *myb28* mutant lines analysed, with the exception of indolylmethyl in generative leaves (Table 1, Table S11).

**Table 1.** Sulphur metabolite levels of total methionine-derived aliphatic glucosinolates (A-GSLs), total tryptophan-derived indolic glucosinolates (I-GSLs) sulphate and S-methyl cysteine sulfoxide (SMCSO) in vegetative leaves, generative leaves and florets of field-grown *myb28* mutant lines.

|  | Vegetative Leaves |  |  |  | Generative Leaves |  |  |  | Florets |  |  |  |
| --- | --- | --- | --- | --- | --- | --- | --- | --- | --- | --- | --- | --- |
|  | Total A-GSLs | Total I-GSLs | Sulphate | SMCSO | Total A-GSLs | Total I-GSLs | Sulphate | SMCSO | Total A-GSLs | Total I-GSLs | Sulphate | SMCSO |
| WT | 1.51 ± 0.76<sup>a</sup> | 1.29 ± 0.77<sup>a</sup> | 503.56 ± 122.51<sup>a</sup> | 11.58 ± 3.21<sup>a</sup> | 6.84 ± 2.96<sup>a</sup> | 7.08 ± 2.03<sup>a</sup> | 438.65 ± 189.31<sup>a</sup> | 18.03 ± 9.91<sup>a</sup> | 15.04 ± 9.36<sup>a</sup> | 7.66 ± 5.36<sup>a</sup> | 253.12 ± 98.25<sup>a</sup> | 88.08 ± 24.50<sup>a</sup> |
| *myb28-1<sup>ge</sup>* | 0.76 ± 0.35<sup>a</sup> | 0.88 ± 0.70<sup>a</sup> | 394.86 ± 108.93<sup>a</sup> | 8.40 ± 2.95<sup>a</sup> | 0.82 ± 0.06<sup>b</sup> | 4.45 ± 0.85<sup>a</sup> | 344.83 ± 178.80<sup>a</sup> | 13.24 ± 4.30<sup>a</sup> | 1.14 ± 0.21<sup>b</sup> | 6.40 ± 2.50<sup>a</sup> | 151.95 ± 48.11<sup>a</sup> | 57.54 ± 15.8<sup>a</sup> |
| *myb28-2<sup>ge</sup>* | 0.60 ± 0.10<sup>a</sup> | 0.91 ± 0.76<sup>a</sup> | 421.66 ± 98.13<sup>a</sup> | 10.54 ± 3.45<sup>a</sup> | 0.83 ± 0.02<sup>b</sup> | 3.73 ± 0.98<sup>b</sup> | 282.06 ± 184.77<sup>a</sup> | 14.52 ± 8.06<sup>a</sup> | 1.01 ± 0.13<sup>b</sup> | 4.24 ± 3.3<sup>a</sup> | 138.71 ± 100.5<sup>a</sup> | 63.55 ± 33.36<sup>a</sup> |

WT = T2 progeny of transgenic CRISPR line with no presence of transgene and wild-type *MYB28* sequences. *bolC2.myb28<sup>ge-1</sup>*. *bolC2.myb28<sup>ge-2</sup>*. Both mutant lines have a large deletion in C9 *MYB28* leading to a non-functional protein. Data are expressed in $\mu\text{mol g}^{-1}$ dry weight as mean ± SD (n=4 plants). Within columns, data followed by the same letter are not significantly different ( $P < 0.05$ ). (See Table S7-S9 for a summary of statistical analyses)

|  | Vegetative Leaves |  |  |  | Generative Leaves |  |  |  | Florets |  |  |  |
| --- | --- | --- | --- | --- | --- | --- | --- | --- | --- | --- | --- | --- |
|  | Total A-GSLs | Total I-GSLs | Sulphate | SMCSO | Total A-GSLs | Total I-GSLs | Sulphate | SMCSO | Total A-GSLs | Total I-GSLs | Sulphate | SMCSO |
| WT | 1.51 ± 0.76<sup>a</sup> | 1.29 ± 0.77<sup>a</sup> | 503.56 ± 122.51<sup>a</sup> | 11.58 ± 3.21<sup>a</sup> | 6.84 ± 2.96<sup>a</sup> | 7.08 ± 2.03<sup>a</sup> | 438.65 ± 189.31<sup>a</sup> | 18.03 ± 9.91<sup>a</sup> | 15.04 ± 9.36<sup>a</sup> | 7.66 ± 5.36<sup>a</sup> | 253.12 ± 98.25<sup>a</sup> | 88.08 ± 24.50<sup>a</sup> |
| *myb28*<br>-1<sup>ge</sup> | 0.76 ± 0.35<sup>a</sup> | 0.88 ± 0.70<sup>a</sup> | 394.86 ± 108.93<sup>a</sup> | 8.40 ± 2.95<sup>a</sup> | 0.82 ± 0.06<sup>b</sup> | 4.45 ± 0.85<sup>a</sup> | 344.83 ± 178.80<sup>a</sup> | 13.24 ± 4.30<sup>a</sup> | 1.14 ± 0.21<sup>b</sup> | 6.40 ± 2.50<sup>a</sup> | 151.95 ± 48.11<sup>a</sup> | 57.54 ± 15.8<sup>a</sup> |
| *myb28*<br>-2<sup>ge</sup> | 0.60 ± 0.10<sup>a</sup> | 0.91 ± 0.76<sup>a</sup> | 421.66 ± 98.13<sup>a</sup> | 10.54 ± 3.45<sup>a</sup> | 0.83 ± 0.02<sup>b</sup> | 3.73 ± 0.98<sup>b</sup> | 282.06 ± 184.77<sup>a</sup> | 14.52 ± 8.06<sup>a</sup> | 1.01 ± 0.13<sup>b</sup> | 4.24 ± 3.3<sup>a</sup> | 138.71 ± 100.5<sup>a</sup> | 63.55 ± 33.36<sup>a</sup> |

WT = T2 progeny of transgenic CRISPR line with no presence of transgene and wild-type *MYB28* sequences. *bolC2.myb28-1<sup>ge</sup>*. *bolC2.myb28-2<sup>ge</sup>*. Data are expressed in $\mu\text{mol g}^{-1}$ dry weight as mean ± SD (n=4 plants). Within columns, data followed by the same letter are not significantly different ( $P < 0.05$ ). (See Table S7-S9 for a summary of statistical analyses)

In primary sulphur metabolism inorganic sulphate is taken up by the roots, with vacuolar sulphate stored as a key precursor for all sulphur containing metabolites, including a bioactive known as s-methyl cysteine sulfoxide or SMCSO. Although a reduction in both sulphate and SMCSO is observed in these field-grown plants when comparing the two mutant lines to wild type, this effect was not significant due to the high standard deviation between samples in this experiment (Table 1).

## Discussion

4-Day old cotyledonary petioles were successfully transformed to deliver the Cas9 enzyme and dual sgRNAs. Evidence of gene editing was found in primary transgenics and in a subsequent T1 generation. The self-fertilised T1 generation produced stable transgene-free T2 homozygous *myb28* mutant line in which the C2 (Bo2g161590) and C9 (Bo9g014610) copies of *MYB28* were rendered non-functional. The C7 (Bo7g098590) copy showed no signs of editing in these lines. Through the use of a single construct, the dual sgRNAs were able to produce small INDELs, a single thymine insertion and a three-base pair deletion, to the C2 copy of MYB28 through the targeting of a single guide (guide 59). From this same construct, a large-scale deletion of 562 base pairs was introduced to the C9 copy of *MYB28* through the targeting of two guides (guide 59 and guide 60) at this locus. Field-grown plants with mutated C2 and C9 *MYB28* copies accumulated lower levels of the aliphatic glucosinolate glucoraphanin in both leaves and florets, and a reduction in the aliphatic glucosinolate glucoiberin. As *MYB28* encodes an R2R3 MYB transcription factor, this metabolomic change is mediated through reduced activation of genes involved in biosynthesis, as demonstrated in the reduced expression of aliphatic glucosinolate biosynthesis genes involved in this pathway, including *BCAT4, BAT5, MAM3, IPMI, IMDH3, CYP79F1, CYP83A1, SUR1, UGT74C1, SOT17* and *SOT18*. The transcriptome analysis suggests that knocking out *myb28* specifically affects the aliphatic glucosinolate biosynthesis pathway. This is supported at the metabolite level as accumulation of sulphur metabolites including indole glucosinolates, sulphate and S-methyl cysteine sulfoxide was not significantly affected. Further investigation would include analysis of the additional primary sulphur metabolites such as the amino acid precursor to aliphatic glucosinolates, methionine, which may have elevated levels of accumulation in the tissues analysed here if it is not being diverted into glucosinolate biosynthesis. In addition, quantifying levels of total sulphur in these tissues and particularly in roots would give insight into a potential role for *MYB28* in encouraging increased sulphur accumulation.

### Mutations in MYB28 DNA Binding Domain

R2R3-MYBs are a subfamily of MYB DNA binding domain transcription factors classified due to the presence of two adjacent repeats in the MYB domain. MYB protein domains are made up of a sequence of roughly 53 amino acids repeat (R) domains which in turn form three alpha helices, characterised by three equally spaced tryptophan residues (Stracke et al., 2001). These tryptophan residues form a hydrophobic core of a helix-turn-helix structure. The “recognition” helix which makes direct contact with the DNA is typically the third helix of each repeat (Jia et al., 2004). Due to the activity of a single guide RNA, mutations characterised here in the C2 and C9 copies of *MYB28* disrupt the single amino acid residue before the third conserved tryptophan in the R2 domain of the R2R3 DNA binding domain. Both mutants used in this analysis had a large-scale deletion in the C9 that would render the protein unable to bind target DNA. This proposed inability to bind DNA is shared at the *MYB28* C2 locus in Mutant 1, in which a thymine insertion introduces a premature stop codon in the R2 domain. The *MYB28* C2 locus in Mutant 2 has a 3-base pair deletion at the same point in the sequence, leading to the removal of a leucine residue. Whilst a single amino acid deletion may be considered a weak allele, *bolC2.myb28-2^ge^* mutants are as severe as *bolC2.myb28-1^ge^* mutants in terms of impact on gene expression and metabolite levels. It is therefore possible that deleting this leucine residue, which is positioned in close proximity to the third conserved tryptophan, has drastic structural consequences to the MYB28 proteins making it unable to bind DNA. An alternative explanation is that the C2 copy is less dominant compared to the other *MYB28* copies in this *B. oleracea* background and that the effect on the phenotype is predominantly down to the large deletion in the C9 copy. However, the prominent role of the C2 copy of *MYB28* is supported by a recent publication in which the A9 and C2 copies of *MYB28* in *Brassica napus* are found to be responsible for glucosinolate accumulation, especially glucoraphanin, in leaves (Kittipol et al., 2019). Further analysis may include protein binding assays and analysis of progeny of back-crossed individuals with single mutations.

### CRISPR-Cas9 in Complex Genomes & Functional Redundancy

The guide RNAs were effective in specifically targeting the characteristic R2R3 MYB DNA binding domain of MYB28, one of many MYB transcription factors found in plants (Dubos et al., 2010). The *Brassica* genus has undergone a whole-genome triplication event within its evolutionary history and has often been considered the foundation on which the multiple morphotypes and domestication events within this lineage were able to occur (Cheng et al., 2014). In particular this whole-genome triplication event has been attributed to the diversity of glucosinolate profiles found throughout the genus, due to multiple copies of glucosinolate biosynthesis genes within the genomes of these agronomically important plants (Hofberger et al., 2013; Wu et al., 2017). From the analysis presented here, the loss of two of the three copies of *MYB28* is sufficient to almost completely abolish accumulation of the aliphatic glucosinolate glucoraphanin. This shows that the C7 copy of *MYB28* cannot compensate for the loss of the C2 and C9 copies, suggesting that the C7 may have sub-functionalised or become a pseudogene following the triplication event. MYB29 is also well characterised as a key regulator in aliphatic glucosinolate production in Arabidopsis, with the *myb28myb29* double mutant phenotype rendering aliphatic glucosinolate accumulation almost undetectable, producing lower aliphatic glucosinolate levels than those found in each respective single mutant phenotype(Gigolashvili et al., 2007; Hirai et al., 2007; Sonderby et al., 2007). Our analyses also indicate that two copies of *MYB29* present in *B. oleracea* cannot compensate for the loss of C2 and C9 copies of *MYB28*. The lack of off-target editing in *MYB29* suggests that RNA guide design has avoided unintended mutations in other *MYB* genes. This is further supported by the highly targeted transcriptional changes of sulphur metabolism genes in the transcriptomes of these mutant plants. Studies have used CRISPR-Cas9 mediated genome editing to target multiple gene copies in the allotetraploid *B. napus* and *B. carinata* genome (Braatz et al., 2017; Okuzaki et al., 2018; Sun et al., 2018; Yang et al., 2017; Yang et al., 2018) (Kirchner et al., 2017) with few reported off-target effects. To date, the only use of RNA-mediated CRISPR-Cas9 genome editing in *Brassica oleracea* is that of the technique used here developed at the John Innes Centre and described in (Lawrenson et al., 2019; Lawrenson et al., 2015).

### MYB28, Glucosinolates and the Environment

Glucosinolates are secondary metabolites produced predominantly by plants of this group to defend the plants from herbivore and pathogen attack (Wink, 1988) with biosynthesis highly susceptible to changes in abiotic factors like temperature change (Mølmann et al., 2015), in addition to abiotic factors such as soil rhizobia (Bressan et al., 2009). It is possible that growing the *myb28* CRISPR mutants in the field in subsequent years, with varied conditions, would affect the accumulation of glucosinolates and regulation of the genes involved in this pathway. Studies in *Arabidopsis thaliana* have found that *myb28* mutant plants showed increased levels of herbivory of lepidopteran insects *Mamestra brassicae* (Beekwilder et al., 2008) and *Spodoptera exigua* (Gigolashvili et al., 2007; Muller et al., 2010). However some insect species, including butterflies (Pieridae) have co-evolved with members of the Brassicaceae to counter-adapt to the effects of these defence compounds (Edger et al., 2015; Winde and Wittstock, 2011). It is possible that the mutant plants in this analysis were more susceptible to increased herbivory levels of aerial organs. An interesting further analysis from this would be to grow these *myb28* mutant plants in a controlled environment with exposure to both generalist and specialist *Brassica* herbivores.

### Concluding remarks

In summary, these analyses comprise the first stable *myb28* knock-out line in a *Brassica oleracea* background and provides a resource to further understand the function of MYB28 transcription factor in regulating sulphur metabolism in this important crop species. The study also describes the first officially-regulated CRISPR field-trial in the UK to require compliance with the 2001/18 EU GMO directive. Altogether this work highlights the immense potential of using gene-editing technology in characterising transcription factor regulators of biosynthetic processes to develop nutritionally fortified foods.

## Experimental Procedures

### Plant Material

The material used in this analysis is *Brassica oleracea* AG DH1012, a double haploid genotype from the *Brassica oleracea* ssp *alboglabra* (A12DHd) and *Brassica oleracea* ssp *italica* (Green Duke GDDH33) mapping population as described in (Bohuon et al., 1996; Bohuon, 1995).

### Guide Design

In total, 4 guides were designed to target the *MYB28* sequences of the C2 (Bo2g161590), C7 (Bo7g098590) and C9 (Bo9g014610) copies. Guides ‘57’ and ‘58’ were designed to target only the C2 copy of *MYB28* and were transformed in the construct named ‘1985’, while guides ‘59’ and ‘60’ were transformed in the construct ‘1986’ and designed to target all three copies of *MYB28* (See Figure S1 for guide sequences and position of these guides on the *MYB28* C2 gene sequence). Guides ‘59’ and ‘60’ were designed to specifically target the three copies of *MYB28* found in the *Brassica oleracea* genome without disrupting other related MYB transcription factors. Guides were designed by first aligning all three copies of *MYB28*, its most related gene *MYB29* (of which there are two confirmed copies in the *B. oleracea* genome) and the 8 genes most closely related to these transcription factors as annotated in Ensembl Plants (Bolser et al., 2016). Alignment was performed using Clustal Omega Multiple Sequence Aligner (Sievers et al., 2011). After alignment, regions of coding sequence which were shared uniquely between the target *MYB28* copies with proximity to an essential ‘NGG’ PAM motif were selected as guides.

### Transformation

Each dual guide vector containing either guides ‘57 and ‘58’ (1985) or guides ‘59’and ‘60’ (1986) was introduced separately to individual 4-day old cotyledonary explants by *Agrobacterium-mediated* transformation, performed as updated from the method published in (Hundleby and Irwin, 2015). A total of 184 positively identified primary transformants were screened, 92 for each construct. Details of the protocol used for *B. oleracea* gene editing, including details of the vector, cloning and transformation process can be found in (Lawrenson et al., 2019; Lawrenson et al., 2015).

### Propagation

Cotyledonary explants underwent were maintained on shoot regeneration media under kanamycin selection, repeated following three weeks of propagation, with successful transformants initially identified by the emergence of green shoots. Transformants were transferred to 100 mL jars (75 x 50mm) containing 25mL of Gamborgs B5 medium, 25mg/L kanamycin and 160mg/L timentin and maintained on this media with increasing kanamycin concentration at 23°C under 16-hour day length until root elongation. QPCR for the NPTII gene was used to confirm and determine copy number of the T-DNA. Plantlets were transferred to soil prior to transferral to the glasshouse where they were grown in controlled environment rooms at 22°C with a 10-hour day length. All plants were bagged in pollen proof bags prior to pollen emergence.

### DNA Extraction

Single leaf tissue from individual *B. oleracea* plantlets was taken prior to planting for DNA extraction using a protocol modified from (Edwards et al., 1991) for genotyping.

### PCR Amplification and Sequencing

PCR reactions were carried out on a GSTORM Thermocycler and prepared to a final volume of 20μl with 150ng of genomic DNA per reaction, using the New England Biolabs Taq Polymerase with Standard Taq Buffer. PCR products were resolved using (1-2%) agarose gel electrophoresis. The subsequent PCR products were sent to Eurofins Genomics® for sequencing using the Ready2Load sequencing service.

### Copy Number Assay for Presence of the Transgene

A copy number assay was performed on leaf material from plantlets prior to planting in the field to confirm the absence of the transgenes. Two components of the dual guide vector were screened, the Cas9 DNA endonuclease and the NPTII component of the vector which encodes a neomycin phosphotransferase II enzyme, allowing transformed plants to metabolize neomycin and kanamycin. This was performed by Peter Isaac at iDNA Genetics using a proprietary method. Only plants with a result of “0” were transplanted into the field (See Table S1 for Cas9 results and Table S2 for NPTII).

### Field Conditions

The *B. oleracea* plants in the field were surrounded by a border of *B. napus* plants with a zone of 20 meters left between the plants in this plot and other test plots of non-GM *Brassica*. The *Brassica* plants were grown in a cage, used in previous field trials by the John Innes Field Experimentation team. Upon completion of the trial all the *Brassica* plants were destroyed, to the satisfaction of the regulator, and the land continuously monitored for the unlikely chance of residual *Brassica* plants emerging.

Planting of four-week-old plantlets took place on 25^th^ April 2019. Sampling of ‘vegetative’ leaf material took place on the 28th May, in which 3 mature leaves were pooled from individual plants which were roughly 8 weeks old. On 24^th^ June 2019 entire plants were uprooted on site prior to flowering, with separation of 3 pooled ‘generative’ leaves and floral organs for each individual plant which were roughly 12 weeks old. At all sampling stages material from individual plants was kept separate and double-contained in sealed containers before being transported to contained freezers at the John Innes Centre. Material was snap frozen in liquid nitrogen and stored at −80 °C for RNA isolation with further freeze-drying and grinding of some material for high performance liquid chromatography (HPLC) analysis. The *B. napus* plants were uprooted and destroyed to the satisfaction of the regulator.

The details of the application and consent conditions can be found at https://www.gov.uk/government/publications/genetically-modified-organisms-john-innes-centre-19r5201.

### RNA-Seq Analysis

*Brassica oleracea* RNA extraction was performed using E.Z.N.A.® Plant RNA Kit provided by Omega Bio-tek Inc (Kit Catalogue Number: R6827-01, DNAse Treatment Catalogue Number: E1091). A total of 27 RNA samples from individual plants were sent to Novogene® for stranded RNA library preparation and sequencing using the Illumina NovaSeq PE150 platform. This included three individual plant replicates of each genotype (wild type, *bolC2.myb28-1^ge^* and *bolC2.myb28-2^ge^)* for each of the three tissue types (pooled ‘vegetative leaves’, pooled ‘generative leaves’ and inflorescences each from a single individual). See Table S3 for the detailed IDs of individual plants, tissues and genotypes sequenced. Raw sequence data is available on the National Centre for Biotechnology Information (NCBI) under BioProject ID PRJNA633531. The package Kallisto v0.44.0 (Bray et al., 2016) was used to produce abundance files for each sample, using the publicly available genome for *Brassica oleracea* TO1000 for mapping of reads (Assembly BOL, INSDC Assembly GCA_000695525.1 version 97.1 (Parkin et al., 2014). Differential expression analysis was performed using edgeR with an FDR cut off of 0.05 and minimum read count of 20 (Robinson et al., 2010). The results of these analyses can be found in Table S4 (Vegetative Leaves) Table S5 (Generative Leaves) and Table S6 (Florets). This data was used to perform Gene Ontology (GO) enrichment analysis using the package TopGo (Alexa and Rahnenführer, 2009) by aligning the *Brassica oleracea* gene models using the *Arabidopsis thaliana* org.At.tair.db library (version 3.2.3) (Carlson, 2016). The results of these analyses can be found in Table S7 (Vegetative Leaves) Table S8 (Generative Leaves) and Table S9 (Florets).

### Metabolite Analysis

Quantification of the following glucosinolates was performed according to (Saha et al., 2012) with the use of glucotropaeolin as the internal standard: Glucoiberin (3-MSP), Glucoraphanin (4-MSB), Hyrdoxyindolylmethyl (OHI3M), Indolylmethyl (I3M), 1 – methoxyindolylmethyl (1MOI3M) and 4 – methoxyindolylmethyl (4MOI3M). Sulphate analysis was performed as modified from (Koprivova et al., 2008). SMCSO analysis was performed as modified from (Kubec and Dadáková, 2009).

### Statistics

Analyses were performed by one-way ANOVA to assess the effect of genotype on metabolite levels with Tukey’s multiple comparisons t-tests. Statistical analyses in the figures portrays Tukey’s multiple comparisons t-test results compared with the corresponding wild type T2 plants. Statistical analyses were performed in GraphPad Prism v.8.2.0, the results of which can be found in Table S10 (Vegetative Leaves) Table S11 (Generative Leaves) and Table S12 (Florets).

## Acknowledgements

We would like to thank Burkhard Steuernagel for providing bioinformatic support, Judith Irwin for access to the *B. oleracea* DH1012 genome sequence and Peter Isaac for performing the copy number screening. We are also grateful to Emilie Knight, Cathy Mumford and the Field Experimentation Team for providing support throughout the field experiments, to Felicity Perry at the John Innes Centre for assistance with public relations and to Monika Chhetry for assistance with *B. oleracea* transformation. This work was support by the UKRI Biotechnology and Biological Sciences Research Council (BBSRC) through institute strategic programmes BB/P013511/1 to the John Innes Centre and BB/R012512/1 and its constituent project(s) BBS/E/F/000PR10343 to the Quadram Institute. MN was supported by the BBSRC Norwich Research Park Biosciences Doctoral Training Partnership grant number BB/M011216/1. We also acknowledge the support of BBSRC grant BB/N019466/1 that allowed production of the CRISPR-Cas9-edited lines.

## Author Contributions

MN, MT, RM and LØ designed the experiments, analysed the data and wrote the paper. MN conducted the experiments. SS^1^ assisted in screening and field experiments. WH oversaw the guide design and transformation process. TL assisted in the guide design process and generated constructs for transformation. PH performed *Brassica oleracea* transformations. PR and SS^2^ assisted in bioinformatic and metabolite analyses, respectively.

## Supporting Information

**Supplementary Figure 1** - Position of guide RNA sequences in relation to the C2 copy

*MYB28* in *Brassica oleracea* 1012

**Supplementary Figure 2** - Alignment of *‘bolC2.myb28-1^ge^’* or ‘M1’ MYB28 C2 gene sequence to wild type (WT)

**Supplementary Figure 3** - Alignment of *‘bolC2.myb28-1^ge^’* ‘M1’ MYB28 C2 protein sequence to wild type (WT)

**Supplementary Figure 4** - Alignment of *‘bolC2.myb28-2^ge^’* or ‘M2’ MYB28 C2 gene sequence to wild type (WT)

**Supplementary Figure 5** - Alignment of *‘bolC2.myb28-2^ge^’* or ‘M2’ MYB28 C2 protein sequence

**Supplementary Figure 6** - Gel Image of MYB28 C9 deletion in *bolC2.myb28-1^ge^’* and *‘bolC2.myb28-2^ge^’* lines compared to two wild type controls

**Supplementary Figure 7** - Alignment of MYB28 C9 protein sequence in *‘bolC2.myb28-1^ge^’* and *‘bolC2.myb28-2^ge^’*

**Supplementary Figure 8** – SPEC PCR Screening

**Supplementary Figure 9** – U626 PCR Screening

**Supplementary Table 1** – Cas9 copy number Screening

**Supplementary Table 2** – NPTII copy number Screening

**Supplementary Table 3** – Summary of samples used in transcriptome analysis

**Supplementary Table 4** - Differentially expressed genes (p<0.05) in the vegetative leaf tissue

**Supplementary Table 5** - Differentially expressed genes (p<0.05) in the generative leaf tissue

**Supplementary Table 6** - Differentially expressed genes (p<0.05) in the floret tissue

**Supplementary Table 7** - GO enrichment analysis of vegetative leaf tissue

**Supplementary Table 8** - GO enrichment analysis of generative leaf tissue

**Supplementary Table 9** – GO enrichment analysis of floret tissue

**Supplementary Table 10** - Statistical analyses of vegetative leaf tissue metabolites

**Supplementary Table 11** - Statistical analyses of generative leaf tissue metabolites

**Supplementary Table 12** - Statistical analyses of floret tissue metabolites

