## Supplemental Information for "CRISPR-Cas9-mediated editing of *myb28* impairs glucoraphanin accumulation of *Brassica oleracea* in the field" (from another version of this preprint: version 1, 2020-07-17)

Neequaye et al. – Supplementary Material

**Supplementary Figure 1**
Position of guide RNA sequences in relation to the C2 copy *MYB28* in *Brassica oleracea* 1012. Guide RNAs are numbered and colour coded, exons are highlighted in yellow. Guides 58 & 59 were in construct 1985, these did not produce lines used in these analyses. Guides 59 and 60 were most effective as part of construct 1986, in which INDELs were introduced to the C2 copy of MYB28 by Guide 59 and a large deletion was introduced to the C9 copy of MYB28 through the combined activities of guides 59 and 60. The T2 progeny of transgenic lines transformed with construct 1986 (Guides 59 and 60) are those used in the analyses described in the main text.

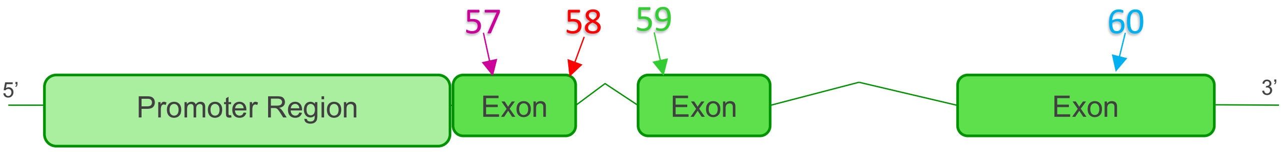

**Guide 57**
**GTCGGAGAAGGGCTGAAGAAAGG**
**Guide 58**
**CGCGACATTCCCCAAAAAGCTGg**
**Guide 59**
**GAAAGAGTTGTAGACTGCGATGG**
**Guide 60**
**GTTCCGACAAGCAATACTCCCGG**

>DH1012_scaffold_29853 MYB28 C2
aacatacattaatactgcacgtcattagtgctaatccttcttttgtgctagaaatgcttcaaaataaatatgtatagcaaaatgtagtaatatataca
gcatcaaattgcaaatatttttcatgttagagaataattgcgagctagtttaaacaaattgatctatttaaataattaataattcgcgtaccaaaatat
ctatcataatcaacattgtgagcatgtacgtgtgtatatatagaagagaatagctcgtaagctcaccaccatcacacaattcattgctcttcttcac
aagtttctgtagtctgatcttatagtgttataaaaatacatataatatttcaaggatatatagaaggcaaaaactcaaagtctacttgaaaaacatg
aaaacacctagcagctctgtgggtaagacccaagagcgcttctcgattagtctcatattcagatgtatcagagttctcattaacagatctatttcttt
cttaccttttagaaaatttcctttctgattttagtttcctcaagtatatttttctcaatatagtatttcctttggtattgtttgaaccttttacacattagtgttcatct
atatatatcggagaaa**ATGTCAAGAAAGCGATGTTGTGTCGGAGAAGGGCTGAAGAAAGGTGCATGG**
**ACCATTGAGGAAGACAAGAAGCTCATCTCTTATATTCACGAACATGGAGAAGGTGGCTGGCGC**
**GACATTCCCCAAAAAGCTG**gttagtatatagtatacacatgtatatttagtatttatatattgtataatattttgatgagaatatggtgttta
attag**GGTTGAAAAGGTGTGGAAAGAGTTGTAGACTGCGATGGACTAACTACCTAAAACCTGAGA**
**TCAAAAGAGGCGAGTTTAGTTCAGAGGAGGAACAGATTATCATCATGCTCCATGCTGCTCGTG**
**GCAACAA**gtacgtttattttagaccaaaaaaaaacaagtacgtttatttttaacagaaaggacgattatatatttttatgtgtgtatggatcctcc
agtgatcatcattctagtttctcttttttttttataccgcaaacaaatttcattagtaaaaaaattaaaattccaaagtgaatattcaaaaacacagtgtt
atataatcctatatatgtcatatattaaaaaagtacaacatgagaaatga
atttaagtatgcttctaaagcgaagttttacttcccgaaaaattattctttattttttcatgtatttgacaattctctgatgcaaaatatgtgtttgattagca
atatgtgactaaaaattgcaatagcacacatcattttagtctctattccataaaaaagcttcaaaataaatttgattaactttggtcttccatcttatctct
ttcactattcttgtctttag**GTGGTCAGTCATAGCTAGACATTTACCTAGAAGAACCGACAATGAGATCAAG**
**AATTACTGGAACACGCATCTCAAAAAACGTTTGATCGAAGAGGGTATTGACCCCGTGACTCAC**
**AAGCCACTAGCTTCTAATACTAACCCTACTGTCACCACCACGCAGCCAGAGAACTTGCATTCC**
**CTAGATGCATCTAGTTCCGACAAGCGATACTCCCGGTCAAGCTCAATGCCTCCCATGTCTCTTC**
**CTCTTTCCTCGGGTTTCAACCCGGTTTCCGAGATCATCACACCAGTTCAGAGAGGTTCCTTGAG**
**TTGCAAGAAAAGTATTAAGAAATCGAACTCTACATCAAGGCTTTTAAACAAAGTTGCTGCTAAG**
**GCCACTACCATCAAAGATATCTTGAGTGCTACTACAATATCATATGAAAGCTTTTCCAATGGCT**
**TCCCTGAGCAGACTCGCAATGAAGAAGATAGTTCTAATGCTCTCGCTGACTTCGATCCATTTCC**
**CCAATCATTGTTGTACACTGAACATGAGGTCCATGCTACTTCTGATCTTGATATGGATCAGGGT**
**TACGATTTCTCATATTTTCTCGAGACACTCGGGAGAGATGAGCACAATACGAATGTCGAGTATG**
**GTCATGATATTCTTCTTATGTCCGATGTGTCACAAGAAGTCTCATCA**

**ATTAGCGTTGATGATCAAGAAAATATGATAGGAAACTTCGAGGGCTGGTCAAATTATCTTCTTG ACCATGATGGTTTGATAAATGACACGGACTATGATTTCCTCGAGAAGCAAATCATATGA**atcttcat atttaaacaaaacggtttcaaactagttgaaacatgcctgaataagaagttatatatgtatttgttatttggattgtttagtagttagtatatg

**Supplementary Figure 2**

Alignment of *bolC2.myb28-1*$^{ge}$ (MYB28C2M1) coding sequence to wild type (MYB28C2WT). The thymine insertion is highlighted in yellow.

```
CLUSTAL O(1.2.4) multiple sequence alignment

MYB28C2WT      ATGTCAAGAAAGCGATGTTGTGTCGGAGAAGGGCTGAAGAAAGGTGCATGGACCATTGAG 60
MYB28C2M1      ATGTCAAGAAAGCGATGTTGTGTCGGAGAAGGGCTGAAGAAAGGTGCATGGACCATTGAG 60
               ************************************************************

MYB28C2WT      GAAGACAAGAAGCTCATCTCTTATATTCACGAACATGGAGAAGGTGGCTGGCGCGACATT 120
MYB28C2M1      GAAGACAAGAAGCTCATCTCTTATATTCACGAACATGGAGAAGGTGGCTGGCGCGACATT 120
               ************************************************************

MYB28C2WT      CCCCAAAAAGCTGGGTTGAAAAGGTGTGGAAAGAGTTGTAGACTG-CGATGGACTAACTA 179
MYB28C2M1      CCCCAAAAAGCTGGGTTGAAAAGGTGTGGAAAGAGTTGTAGACTGTCGATGGACTAACTA 180
               ********************************************* **************

MYB28C2WT      CCTAAAACCTGAGATCAAAAGAGGCGAGTTTAGTTCAGAGGAGGAACAGATTATCATCAT 239
MYB28C2M1      CCTAAAACCTGAGATCAAAAGAGGCGAGTTTAGTTCAGAGGAGGAACAGATTATCATCAT 240
               ************************************************************

MYB28C2WT      GCTCCATGCTGCTCGTGGCAACAAGTGGTCAGTCATAGCTAGACATTTACCTAGAAGAAC 299
MYB28C2M1      GCTCCATGCTGCTCGTGGCAACAAGTGGTCAGTCATAGCTAGACATTTACCTAGAAGAAC 300
               ************************************************************

MYB28C2WT      CGACAATGAGATCAAGAATTACTGGAACACGCATCTCAAAAAACGTTTGATCGAAGAGGG 359
MYB28C2M1      CGACAATGAGATCAAGAATTACTGGAACACGCATCTCAAAAAACGTTTGATCGAAGAGGG 360
               ************************************************************

MYB28C2WT      TATTGACCCCGTGACTCACAAGCCACTAGCTTCTAATACTAACCCTACTGTCACCACCAC 419
MYB28C2M1      TATTGACCCCGTGACTCACAAGCCACTAGCTTCTAATACTAACCCTACTGTCACCACCAC 420
               ************************************************************

MYB28C2WT      GCAGCCAGAGAACTTGCATTCCCTAGATGCATCTAGTTCCGACAAGCGATACTCCCGGTC 479
MYB28C2M1      GCAGCCAGAGAACTTGCATTCCCTAGATGCATCTAGTTCCGACAAGCGATACTCCCGGTC 480
               ************************************************************

MYB28C2WT      AAGCTCAATGCCTCCCATGTCTCTTCCTCTTTCCTCGGGTTTCAACCCGGTTTCCGAGAT 539
MYB28C2M1      AAGCTCAATGCCTCCCATGTCTCTTCCTCTTTCCTCGGGTTTCAACCCGGTTTCCGAGAT 540
               ************************************************************

MYB28C2WT      CATCACACCAGTTCAGAGAGGTTCCTTGAGTTGCAAGAAAAGTATTAAGAAATCGAACTC 599
MYB28C2M1      CATCACACCAGTTCAGAGAGGTTCCTTGAGTTGCAAGAAAAGTATTAAGAAATCGAACTC 600
               ************************************************************

MYB28C2WT      TACATCAAGGCTTTTAAACAAAGTTGCTGCTAAGGCCACTACCATCAAAGATATCTTGAG 659
MYB28C2M1      TACATCAAGGCTTTTAAACAAAGTTGCTGCTAAGGCCACTACCATCAAAGATATCTTGAG 660
               ************************************************************

MYB28C2WT      TGCTACTACAATATCATATGAAAGCTTTTCCAATGGCTTCCCTGAGCAGACTCGCAATGA 719
MYB28C2M1      TGCTACTACAATATCATATGAAAGCTTTTCCAATGGCTTCCCTGAGCAGACTCGCAATGA 720
               ************************************************************

MYB28C2WT      AGAAGATAGTTCTAATGCTCTCGCTGACTTCGATCCATTTCCCCAATCATTGTTGTACAC 779
MYB28C2M1      AGAAGATAGTTCTAATGCTCTCGCTGACTTCGATCCATTTCCCCAATCATTGTTGTACAC 780
               ************************************************************

MYB28C2WT      TGAACATGAGGTCCATGCTACTTCTGATCTTGATATGGATCAGGGTTACGATTTCTCATA 839
MYB28C2M1      TGAACATGAGGTCCATGCTACTTCTGATCTTGATATGGATCAGGGTTACGATTTCTCATA 840
               ************************************************************

MYB28C2WT      TTTTCTCGAGACACTCGGGAGAGATGAGCACAATACGAATGTCGAGTATGGTCATGATAT 899
MYB28C2M1      TTTTCTCGAGACACTCGGGAGAGATGAGCACAATACGAATGTCGAGTATGGTCATGATAT 900
               ************************************************************
```

```
MYB28C2WT      TCTTCTTATGTCCGATGTGTCACAAGAAGTCTCATCAATTAGCGTTGATGATCAAGAAAA 959
MYB28C2M1      TCTTCTTATGTCCGATGTGTCACAAGAAGTCTCATCAATTAGCGTTGATGATCAAGAAAA 960
               ************************************************************

MYB28C2WT      TATGATAGGAAACTTCGAGGGCTGGTCAAATTATCTTCTTGACCATGATGGTTTGATAAA 1019
MYB28C2M1      TATGATAGGAAACTTCGAGGGCTGGTCAAATTATCTTCTTGACCATGATGGTTTGATAAA 1020
               ************************************************************

MYB28C2WT      TGACACGGACTATGATTTCCTCGAGAAGCAAATCATATGA    1059
MYB28C2M1      TGACACGGACTATGATTTCCTCGAGAAGCAAATCATATGA    1060
               ****************************************
```

**Supplementary Figure 3**

Alignment of *bolC2.myb28-1[ge]* (MYB28C2M1) protein sequence to wild type (MYB28C2WT).

```
MYB28C2WT       MSRKRCCVGEGLKKGAWTIEEDKKLISYIHEHGEGGWRDIPQKAGLKRCGKSCRLRWTNY 60
MYB28C2M1       MSRKRCCVGEGLKKGAWTIEEDKKLISYIHEHGEGGWRDIPQKAGLKRCGKSCRLSMD*L 59
                *******************************************************

MYB28C2WT       LKPEIKRGEFSSEEEQIIIMLHAARGNKWSVIARHLPRRTDNEI---KNYWNTHLKKRLI 117
MYB28C2M1       PKT*DQKRRV*FRG---GTDYHHAPCCSWQQVVSHS*TFT*KNRQ*DQELLEHASQKTFD 111
                 *   :: ..  .        * *   .*. :. *    * ::    ::  :   :* :

MYB28C2WT       EEGIDPVTHKPLASNTNPTVTTTQPENLHSLDASSSDKRYSRSSSMPPMSLPLSSGFNPV 177
MYB28C2M1       RRGY*PRDSQATSF*Y*---------------------PYCHHHAARELAFPRCI*FR-- 144
                ..*  *   :  :                          *.:  :   :::* .  *.

MYB28C2WT       SEIITPVQRGSLSCKKSIKKSNSTSRLLNKVAAKATTIKDILSATTISYESFSNGFPEQT 237
MYB28C2M1       -QAILPVKLN-------------------------------ASHVSSSFLGFQPGFRDHH 172
                 : * **: .                                * .: *: .*. ** ::

MYB28C2WT       RNEEDSSNALADFDPFPQSLLYTEHEVHATSDLDMDQ----------------------- 274
MYB28C2M1       TSSERFLEL---QEKY*EIELYIKAFKQSCC*GHYHQRYLECYYNII*KLFQWLP*ADSQ 225
                 ..*   :     : : :  ** :   :: .  . .*

MYB28C2WT       ----------------------GYDFSYFLETLGRDEHNTNVEYGHDILLMSDVSQEVSS 312
MYB28C2M1       *RR*F*CSR*LRSISPIIVVH*T*GPCYF*S*YGS---------GLRFLIFSRDTRER-- 266
                                        . .** .  *          *  :*::*  ::*

MYB28C2WT       ISVDDQENMIGNFEGWSNYLLDH----DGLIND----TDYDFLEKQII------------ 352
MYB28C2M1       -------*AQYECRVWS*YSSYVRCVTRSLIN*R**SRKYDRKLRGLVKLSS*P*WFDK* 311
                           : . ** *         .***      .**   : ::

MYB28C2WT       ------------       352
MYB28C2M1       HGL*FPREANHM       322
```

**Supplementary Figure 4**

Alignment of *bolC2.myb28-2^ge^* (MYB28C2M2) coding sequence to wild type (MYB28C2WT). The three base pair deletion is highlighted in yellow.

```
CLUSTAL O(1.2.4) multiple sequence alignment

MYB28C2WT      ATGTCAAGAAAGCGATGTTGTGTCGGAGAAGGGCTGAAGAAAGGTGCATGGACCATTGAG 60
MYB28C2M2      ATGTCAAGAAAGCGATGTTGTGTCGGAGAAGGGCTGAAGAAAGGTGCATGGACCATTGAG 60
               ************************************************************

MYB28C2WT      GAAGACAAGAAGCTCATCTCTTATATTCACGAACATGGAGAAGGTGGCTGGCGCGACATT 120
MYB28C2M2      GAAGACAAGAAGCTCATCTCTTATATTCACGAACATGGAGAAGGTGGCTGGCGCGACATT 120
               ************************************************************

MYB28C2WT      CCCCAAAAAGCTGGGTTGAAAAGGTGTGGAAAGAGTTGTAGACTGCGATGGACTAACTAC 180
MYB28C2M2      CCCCAAAAAGCTGGGTTGAAAAGGTGTGGAAAGAGTTGTAGA---CGATGGACTAACTAC 177
               ******************************************   ***************

MYB28C2WT      CTAAAACCTGAGATCAAAAGAGGCGAGTTTAGTTCAGAGGAGGAACAGATTATCATCATG 240
MYB28C2M2      CTAAAACCTGAGATCAAAAGAGGCGAGTTTAGTTCAGAGGAGGAACAGATTATCATCATG 237
               ************************************************************

MYB28C2WT      CTCCATGCTGCTCGTGGCAACAAGTGGTCAGTCATAGCTAGACATTTACCTAGAAGAACC 300
MYB28C2M2      CTCCATGCTGCTCGTGGCAACAAGTGGTCAGTCATAGCTAGACATTTACCTAGAAGAACC 297
               ************************************************************

MYB28C2WT      GACAATGAGATCAAGAATTACTGGAACACGCATCTCAAAAAACGTTTGATCGAAGAGGGT 360
MYB28C2M2      GACAATGAGATCAAGAATTACTGGAACACGCATCTCAAAAAACGTTTGATCGAAGAGGGT 357
               ************************************************************

MYB28C2WT      ATTGACCCCGTGACTCACAAGCCACTAGCTTCTAATACTAACCCTACTGTCACCACCACG 420
MYB28C2M2      ATTGACCCCGTGACTCACAAGCCACTAGCTTCTAATACTAACCCTACTGTCACCACCACG 417
               ************************************************************

MYB28C2WT      CAGCCAGAGAACTTGCATTCCCTAGATGCATCTAGTTCCGACAAGCGATACTCCCGGTCA 480
MYB28C2M2      CAGCCAGAGAACTTGCATTCCCTAGATGCATCTAGTTCCGACAAGCGATACTCCCGGTCA 477
               ************************************************************

MYB28C2WT      AGCTCAATGCCTCCCATGTCTCTTCCTCTTTCCTCGGGTTTCAACCCGGTTTCCGAGATC 540
MYB28C2M2      AGCTCAATGCCTCCCATGTCTCTTCCTCTTTCCTCGGGTTTCAACCCGGTTTCCGAGATC 537
               ************************************************************

MYB28C2WT      ATCACACCAGTTCAGAGAGGTTCCTTGAGTTGCAAGAAAAGTATTAAGAAATCGAACTCT 600
MYB28C2M2      ATCACACCAGTTCAGAGAGGTTCCTTGAGTTGCAAGAAAAGTATTAAGAAATCGAACTCT 597
               ************************************************************

MYB28C2WT      ACATCAAGGCTTTTAAACAAAGTTGCTGCTAAGGCCACTACCATCAAAGATATCTTGAGT 660
MYB28C2M2      ACATCAAGGCTTTTAAACAAAGTTGCTGCTAAGGCCACTACCATCAAAGATATCTTGAGT 657
               ************************************************************

MYB28C2WT      GCTACTACAATATCATATGAAAGCTTTTCCAATGGCTTCCCTGAGCAGACTCGCAATGAA 720
MYB28C2M2      GCTACTACAATATCATATGAAAGCTTTTCCAATGGCTTCCCTGAGCAGACTCGCAATGAA 717
               ************************************************************

MYB28C2WT      GAAGATAGTTCTAATGCTCTCGCTGACTTCGATCCATTTCCCCAATCATTGTTGTACACT 780
MYB28C2M2      GAAGATAGTTCTAATGCTCTCGCTGACTTCGATCCATTTCCCCAATCATTGTTGTACACT 777
               ************************************************************

MYB28C2WT      GAACATGAGGTCCATGCTACTTCTGATCTTGATATGGATCAGGGTTACGATTTCTCATAT 840
MYB28C2M2      GAACATGAGGTCCATGCTACTTCTGATCTTGATATGGATCAGGGTTACGATTTCTCATAT 837
               ************************************************************

MYB28C2WT      TTTCTCGAGACACTCGGGAGAGATGAGCACAATACGAATGTCGAGTATGGTCATGATATT 900
MYB28C2M2      TTTCTCGAGACACTCGGGAGAGATGAGCACAATACGAATGTCGAGTATGGTCATGATATT 897
               ************************************************************
```

```
MYB28C2WT      CTTCTTATGTCCGATGTGTCACAAGAAGTCTCATCAATTAGCGTTGATGATCAAGAAAAT 960
MYB28C2M2      CTTCTTATGTCCGATGTGTCACAAGAAGTCTCATCAATTAGCGTTGATGATCAAGAAAAT 957
               ************************************************************

MYB28C2WT      ATGATAGGAAACTTCGAGGGCTGGTCAAATTATCTTCTTGACCATGATGGTTTGATAAAT 1020
MYB28C2M2      ATGATAGGAAACTTCGAGGGCTGGTCAAATTATCTTCTTGACCATGATGGTTTGATAAAT 1017
               ************************************************************

MYB28C2WT      GACACGGACTATGATTTCCTCGAGAAGCAAATCATATGA     1059
MYB28C2M2      GACACGGACTATGATTTCCTCGAGAAGCAAATCATATGA     1056
               ***************************************
```

**Supplementary Figure 5**

Alignment of *bolC2.myb28-2^ge^* (MYB28C2M2) protein sequence to wild type (MYB28C2WT). The resulting leucine deletion is highlighted in yellow.

```
MYB28C2WT      MSRKRCCVGEGLKKGAWTIEEDKKLISYIHEHGEGGWRDIPQKAGLKRCGKSCRLRWTNY 60
MYB28C2M2      MSRKRCCVGEGLKKGAWTIEEDKKLISYIHEHGEGGWRDIPQKAGLKRCGKSCR-RWTNY 59
               ****************************************************** *****

MYB28C2WT      LKPEIKRGEFSSEEEQIIIMLHAARGNKWSVIARHLPRRTDNEIKNYWNTHLKKRLIEEG 120
MYB28C2M2      LKPEIKRGEFSSEEEQIIIMLHAARGNKWSVIARHLPRRTDNEIKNYWNTHLKKRLIEEG 119
               ************************************************************

MYB28C2WT      IDPVTHKPLASNTNPTVTTTQPENLHSLDASSSDKRYSRSSSMPPMSLPLSSGFNPVSEI 180
MYB28C2M2      IDPVTHKPLASNTNPTVTTTQPENLHSLDASSSDKRYSRSSSMPPMSLPLSSGFNPVSEI 179
               ************************************************************

MYB28C2WT      ITPVQRGSLSCKKSIKKSNSTSRLLNKVAAKATTIKDILSATTISYESFSNGFPEQTRNE 240
MYB28C2M2      ITPVQRGSLSCKKSIKKSNSTSRLLNKVAAKATTIKDILSATTISYESFSNGFPEQTRNE 239
               ************************************************************

MYB28C2WT      EDSSNALADFDPFPQSLLYTEHEVHATSDLDMDQGYDFSYFLETLGRDEHNTNVEYGHDI 300
MYB28C2M2      EDSSNALADFDPFPQSLLYTEHEVHATSDLDMDQGYDFSYFLETLGRDEHNTNVEYGHDI 299
               ************************************************************

MYB28C2WT      LLMSDVSQEVSSISVDDQENMIGNFEGWSNYLLDHDGLINDTDYDFLEKQII-        352
MYB28C2M2      LLMSDVSQEVSSISVDDQENMIGNFEGWSNYLLDHDGLINDTDYDFLEKQII*        351
               ****************************************************
```

**Supplementary Figure 6**

Gel Image of MYB28 C9 deletion in both *bolC2.myb28-1$^{ge}$* (M1) and *bolC2.myb28-2$^{ge}$* (M2) lines compared to two wild type controls in which WT refers to true wild type *Brassica oleracea* DH1012 and WT WT2 refers to the lines used in the field analysis which underwent transformation and propagation alongside the mutant lines but are wild-type for all *MYB28* sequences and transgene free.

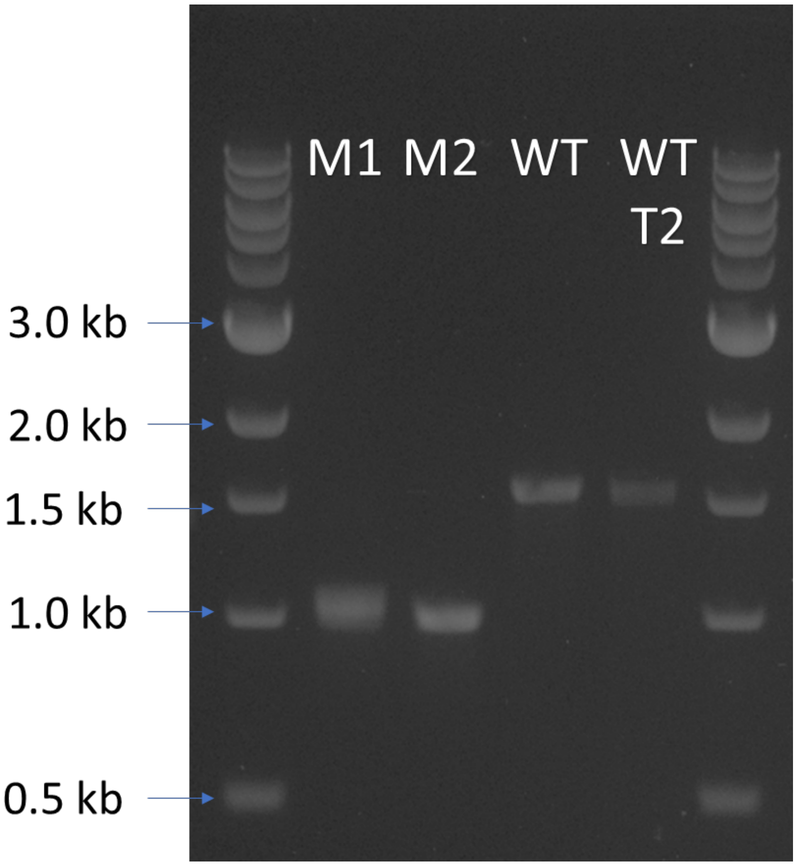

**Supplementary Figure 7**

Alignment of MYB28 C9 protein sequence in both *bolC2.myb28-1$^{ge}$* and *bolC2.myb28-2$^{ge}$*.

```
CLUSTAL O(1.2.4) multiple sequence alignment

WTC9         MSRKPCCVGEGLKKGAWTTEEDKKLISYIHEHGEGGWRDIPQKAGLKRCGKSCRLRWTNY     60
MutantC9     MSRKPCCVGEGLKKGAWTTEEDKKLISYIHEHGEGGWRDIPQKAGY*-------------     46
             *********************************************

WTC9         LKPEIKRGEFSSEEEQIIIMLHASRGNKWSVIARHLPRRTDNEIKNYWNTHLKKRLIEQG     120
MutantC9     ------------------------------------------------------------     46

WTC9         THPLTHKPLASNTNPTVPENLHSLDASSNSDKQYSRSSSMPSMSCTPSSGFNTVFENTSK     180
MutantC9     -------------------------------------------DCTPSSGFNTVFENTSK     63
                                                        .****************

WTC9         DGTPVREDDSLSRKKRFKKSSSTSRLLNKVAAKATSMKEALSASMEGSLNANISFSNGYS     240
MutantC9     DGTPVREDDSLSRKKRLKKSSSTSRLLNKVAAKATSMKKALSASMEGSLNANISFSNGYS     123
             ****************:*********************:*********************

WTC9         EQILNEDDSSNASLINTLAEFDPFLQTTFYPENEMNTTSDLGIDQDYFSHFLENFGNHNE     300
MutantC9     EQILNEDDSSNASLINTLAEFDPFLQTTFYPENEMNTTSDLGIDQDYFSHFLENFGNHNE     183
             ************************************************************

WTC9         EHYMNHNYGHGLLMSYVSQEVSSTSVDDQDNTNEGWSNYLLDHADFIHDMDSDSLGKHLI     360
MutantC9     EHYMNHNYGHGLLMSYVSQEVSSTSVDDQDNTNEGWSNYLLDHADFIHDMDSDSLGKHLI     243
             ************************************************************
```

**Supplementary Figure 8**

The ‘SPEC’ sequence, encoding an antibiotic resistance gene for selection on media was screened, producing a 1673bp fragment which covers the entire length of the SPEC CDS (1008bp) as well as flanking sequences found at the 3’ and 5’ ends of the sequence (31bp upstream and 634bp downstream). This PCR returned positive results for 5 of the positive controls, with all negative controls confirmed. None of the plants screened were positive for this aspect of the vector.

Positive controls identified by + symbol
Negative controls identified by – symbol
Blank controls identified by ‘B’

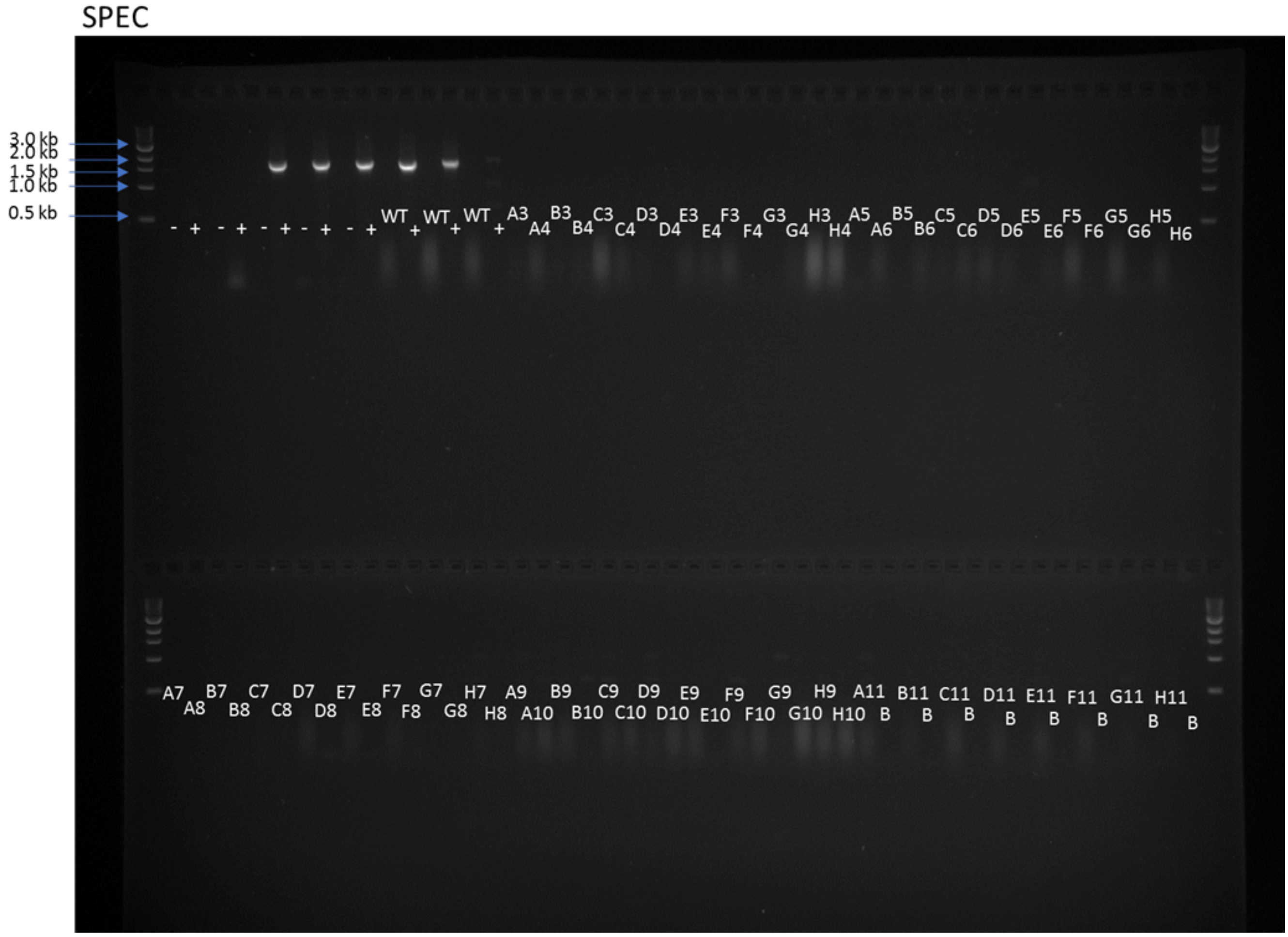

**Supplementary Figure 9**
Using PCR amplification the U6 26 promoter, used to drive the expression of sgRNA was conducted, this was using PCR primers designed to amplify a 685bp fragment. All positive controls amplified the fragment as well as the 11 plants being screened which correspond to the 11 plants identified to be Cas9 and NPTII positive in the copy number assay performed by Peter Isaac at iDNA Genetics.

Positive controls identified by + symbol
Negative controls identified by – symbol
Blank controls identified by 'B'

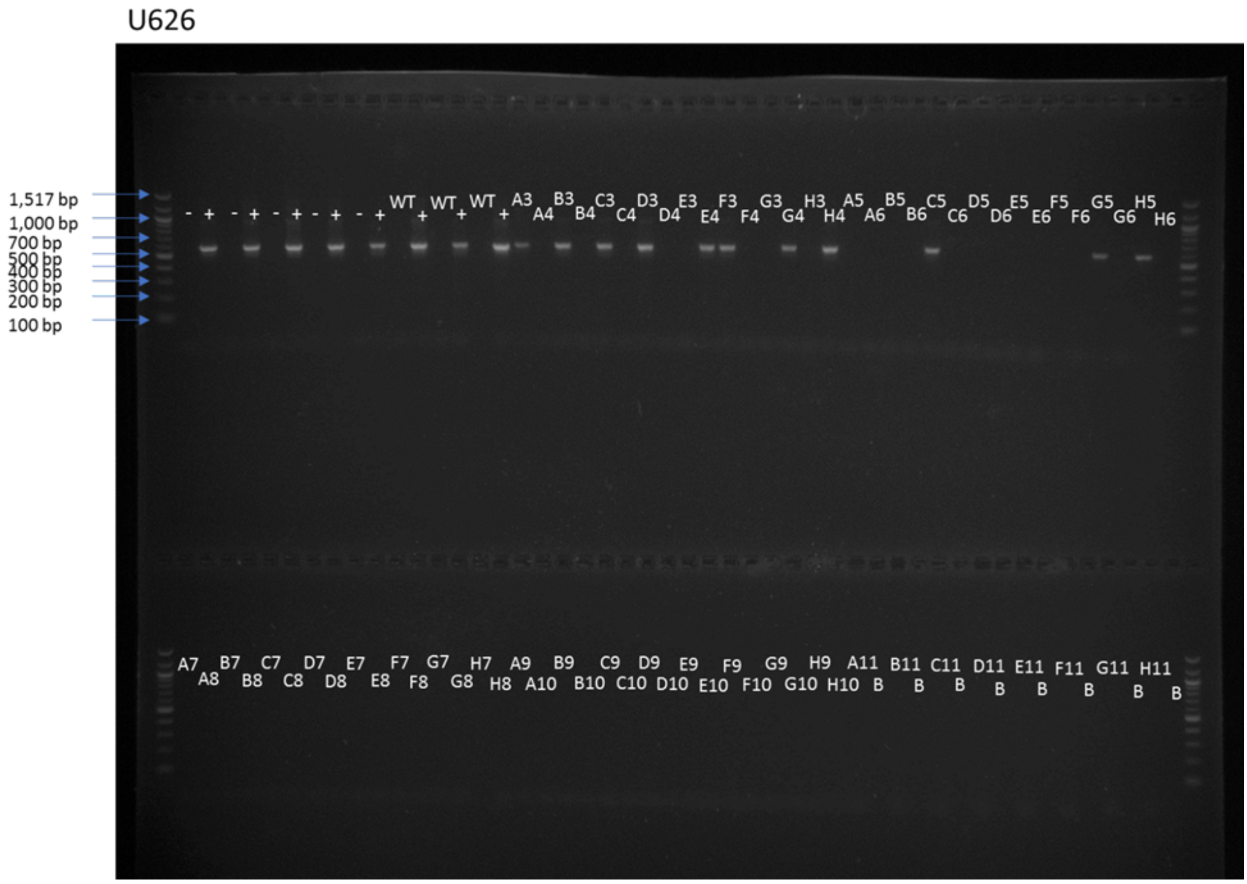

**Supplementary Table 1**
To screen for the presence of the Cas9 DNA endonuclease, which is responsible for the 'cutting' or gene-editing step of the genetic modification, we used an external service provided by iDna Genetics Ltd. in which copy number of the gene in question is assayed. This screen was performed by Dr Peter Isaac who has given his consent to be contacted with any queries regarding the nature and details of this diagnostic service. From this screen 11 of the 75 plants tested were found to have a single (Hemizygous) or double (Homozygous) copy of the Cas9 (see below). Plants with a positive result for the Cas9 or 'no amplification' were not transplanted into the field, only those with a 'Null' result were taken forward.

| Sample | Copies_CRISPR | Assumed_Zygosity |
|---|---|---|
| A10 F7 | 0 | Null |
| A11 F7 | 0 | Null |
| A3 C10 | 1 | Hemi |
| A4 C10 | 0 | Null |
| A5 C10 | No_amplification | |
| A6 F7 | 0 | Null |
| A7 F7 | 0 | Null |
| A8 F7 | 0 | Null |
| A9 F7 | 0 | Null |
| B10 F7 | 0 | Null |
| B11 F7 | 0 | Null |
| B3 C10 | 1 | Hemi |
| B4 C10 | 0 | Null |
| B5 C10 | No_amplification | |
| B6 F7 | 0 | Null |
| B7 F7 | 0 | Null |
| B8 F7 | 0 | Null |
| B9 F7 | 0 | Null |
| Blank | No_amplification | |
| Blank | No_amplification | |
| Blank | No_amplification | |
| Blank | No_amplification | |
| Blank | No_amplification | |

| | | |
|---|---|---|
| Blank | No_amplification | |
| Blank | No_amplification | |
| Blank | No_amplification | |
| Blank | No_amplification | |
| Blank | No_amplification | |
| Blank | No_amplification | |
| Blank | No_amplification | |
| Blank | No_amplification | |
| C10 F7 | 0 | Null |
| C11 F7 | 0 | Null |
| C3 C10 | 2 | Hom |
| C4 C10 | 0 | Null |
| C5 C10 | 1 | Hemi |
| C6 F7 | 0 | Null |
| C7 F7 | 0 | Null |
| C8 F7 | 0 | Null |
| C9 F7 | 0 | Null |
| D10 F7 | 0 | Null |
| D11 F7 | 0 | Null |
| D3 C10 | 1 | Hemi |
| D4 C10 | 0 | Null |
| D5 C10 | 0 | Null |
| D6 F7 | 0 | Null |
| D7 F7 | 0 | Null |
| D8 F7 | 0 | Null |
| D9 F7 | 0 | Null |
| E10 F7 | 0 | Null |
| E11 F7 | 0 | Null |
| E3 C10 | 0 | Null |
| E4 C10 | 2 | Hom |

| | | |
|---|---|---|
| E5 C10 | 0 | Null |
| E6 F7 | 0 | Null |
| E7 F7 | 0 | Null |
| E8 F7 | 0 | Null |
| E9 F7 | 0 | Null |
| F1 WT | 0 | Null |
| F10 F7 | 0 | Null |
| F11 F7 | 0 | Null |
| F3 C10 | 1 | Hemi |
| F4 C10 | 0 | Null |
| F5 C10 | 0 | Null |
| F6 F7 | 0 | Null |
| F7 F7 | 0 | Null |
| F8 F7 | 0 | Null |
| F9 F7 | 0 | Null |
| G1 WT | 0 | Null |
| G10 F7 | 0 | Null |
| G11 F7 | 0 | Null |
| G3 C10 | No_amplification | |
| G4 C10 | 1 | Hemi |
| G5 C10 | 1 | Hemi |
| G6 F7 | 0 | Null |
| G7 F7 | 0 | Null |
| G8 F7 | 0 | Null |
| G9 F7 | 0 | Null |
| H1 WT | 0 | Null |
| H10 F7 | 0 | Null |
| H11 F7 | 0 | Null |
| H3 C10 | 0 | Null |
| H4 C10 | 2 | Hom |

| | | |
|---|---|---|
| H5 C0 | 1 | Hemi |
| H6 F7 | 0 | Null |
| H7 F7 | 0 | Null |
| H8 F7 | 0 | Null |
| H9 F7 | 0 | Null |

**Supplementary Table 2**

In addition to the Cas9, iDna Genetics Ltd also performed copy number analysis of the NPTII component of the vector which encodes a neomycin phosphotransferase II enzyme which allows transformed plants to metabolize neomycin and kanamycin. This screen yielded the same results as the Cas9 screen with the same 11 of the 75 plants tested found to have a single (Hemizygous) or double (Homozygous) copy of NPTII, this was also confirmed by PCR amplification of the U6 26 promoter (see below). Plants with a positive result for NPTII or 'no amplification' were not transplanted into the field, only those with a 'Null' result were taken forward.

| Sample | Copies_NPT2 | Assumed Zygosity |
|---|---|---|
| A10 F7 | 0 | Null |
| A11 F7 | 0 | Null |
| A3 C10 | 1 | Hemi |
| A4 C10 | 0 | Null |
| A5 C10 | No_Amplification | No_Amplification |
| A6 F7 | 0 | Null |
| A7 F7 | 0 | Null |
| A8 F7 | 0 | Null |
| A9 F7 | 0 | Null |
| B10 F7 | 0 | Null |
| B11 F7 | 0 | Null |
| B3 C10 | 1 | Hemi |
| B4 C10 | 0 | Null |
| B5 C10 | No_Amplification | No_Amplification |
| B6 F7 | 0 | Null |
| B7 F7 | 0 | Null |
| B8 F7 | 0 | Null |
| B9 F7 | 0 | Null |
| Blank | weak amplification, contamination? | weak amplification, contamination? |
| Blank | weak amplification, contamination? | weak amplification, contamination? |
| Blank | No_Amplification | No_Amplification |
| Blank | No_Amplification | No_Amplification |

| | | |
|---|---|---|
| Blank | No_Amplification | No_Amplification |
| Blank | No_Amplification | No_Amplification |
| Blank | No_Amplification | No_Amplification |
| Blank | No_Amplification | No_Amplification |
| Blank | No_Amplification | No_Amplification |
| Blank | No_Amplification | No_Amplification |
| Blank | No_Amplification | No_Amplification |
| Blank | No_Amplification | No_Amplification |
| Blank | No_Amplification | No_Amplification |
| C10 F7 | 0 | Null |
| C11 F7 | 0 | Null |
| C3 C10 | 2 | Hom |
| C4 C10 | 0 | Null |
| C5 C10 | 1 | Hemi |
| C6 F7 | 0 | Null |
| C7 F7 | 0 | Null |
| C8 F7 | 0 | Null |
| C9 F7 | 0 | Null |
| D10 F7 | 0 | Null |
| D11 F7 | 0 | Null |
| D3 C10 | 1 | Hemi |
| D4 C10 | 0 | Null |
| D5 C10 | 0 | Null |
| D6 F7 | 0 | Null |
| D7 F7 | 0 | Null |
| D8 F7 | 0 | Null |
| D9 F7 | 0 | Null |
| E10 F7 | 0 | Null |
| E11 F7 | 0 | Null |
| E3 C10 | 0 | Null |

| | | |
|---|---|---|
| E4 C10 | 2 | Hom |
| E5 C10 | 0 | Null |
| E6 F7 | 0 | Null |
| E7 F7 | 0 | Null |
| E8 F7 | 0 | Null |
| E9 F7 | 0 | Null |
| F1 WT | 0 | Null |
| F10 F7 | 0 | Null |
| F11 F7 | 0 | Null |
| F3 C10 | 1 | Hemi |
| F4 C10 | 0 | Null |
| F5 C10 | 0 | Null |
| F6 F7 | 0 | Null |
| F7 F7 | 0 | Null |
| F8 F7 | 0 | Null |
| F9 F7 | 0 | Null |
| G1 WT | 0 | Null |
| G10 F7 | 0 | Null |
| G11 F7 | 0 | Null |
| G3 C10 | weak amplification, contamination? | weak amplification, contamination? |
| G4 C10 | 1 | Hemi |
| G5 C10 | 1 | Hemi |
| G6 F7 | 0 | Null |
| G7 F7 | 0 | Null |
| G8 F7 | 0 | Null |
| G9 F7 | 0 | Null |
| H1 WT | 0 | Null |
| H10 F7 | 0 | Null |
| H11 F7 | 0 | Null |
| H3 C10 | 0 | Null |

| | | |
|---|---|---|
| H4 C10 | 2 | Hom |
| H5 C0 | 1 | Hemi |
| H6 F7 | 0 | Null |
| H7 F7 | 0 | Null |
| H8 F7 | 0 | Null |
| H9 F7 | 0 | Null |

**Supplementary Table 3**
Table of individual plants and tissues used for the transcriptome analysis. Sample ID corresponds to individual plants IDs used throughout screening and analysis.

| **Sample ID** | **Species** | **Genotype** | **Tissue source** |
|---|---|---|---|
| A4_W_F | *Brassica oleracea* | Wild-Type | Florets |
| F5_W_F | *Brassica oleracea* | Wild-Type | Florets |
| H3_W_F | *Brassica oleracea* | Wild-Type | Florets |
| B11_1_F | *Brassica oleracea* | *myb28-1$^{ge}$* | Florets |
| F9_1_F | *Brassica oleracea* | *myb28-1$^{ge}$* | Florets |
| H9_1_F | *Brassica oleracea* | *myb28-1$^{ge}$* | Florets |
| B10_2_F | *Brassica oleracea* | *myb28-2$^{ge}$* | Florets |
| D9_2_F | *Brassica oleracea* | *myb28-2$^{ge}$* | Florets |
| E10_2_F | *Brassica oleracea* | *myb28-2$^{ge}$* | Florets |
| E3_W_JL | *Brassica oleracea* | Wild-Type | Generative Leaves |
| F5_W_JL | *Brassica oleracea* | Wild-Type | Generative Leaves |
| H3_W_JL | *Brassica oleracea* | Wild-Type | Generative Leaves |
| B11_1_JL | *Brassica oleracea* | *myb28-1$^{ge}$* | Generative Leaves |
| F9_1_JL | *Brassica oleracea* | *myb28-1$^{ge}$* | Generative Leaves |
| G10_1_JL | *Brassica oleracea* | *myb28-1$^{ge}$* | Generative Leaves |
| A10_2_JL | *Brassica oleracea* | *myb28-2$^{ge}$* | Generative Leaves |
| B10_2_JL | *Brassica oleracea* | *myb28-2$^{ge}$* | Generative Leaves |
| D9_2_JL | *Brassica oleracea* | *myb28-2$^{ge}$* | Generative Leaves |
| E3_W_ML | *Brassica oleracea* | Wild-Type | Vegetative Leaves |
| F5_W_ML | *Brassica oleracea* | Wild-Type | Vegetative Leaves |
| H3_W_ML | *Brassica oleracea* | Wild-Type | Vegetative Leaves |

| | | | |
|---|---|---|---|
| B11_1_ML | *Brassica oleracea* | *myb28-1$^{ge}$* | Vegetative Leaves |
| F9_1_ML | *Brassica oleracea* | *myb28-1$^{ge}$* | Vegetative Leaves |
| G10_1_ML | *Brassica oleracea* | *myb28-1$^{ge}$* | Vegetative Leaves |
| A10_2_ML | *Brassica oleracea* | *myb28-2$^{ge}$* | Vegetative Leaves |
| B10_2_ML | *Brassica oleracea* | *myb28-2$^{ge}$* | Vegetative Leaves |
| D9_2_ML | *Brassica oleracea* | *myb28-2$^{ge}$* | Vegetative Leaves |

**Supplementary Table 4**

All significantly differentially expressed genes (p<0.05) in the vegetative leaf tissue of both mutant lines, as normalised to wild type, including False Discovery rate correction (FDR), average expression and p values. Note: not all differentially expressed genes (denoted by gene IDs) were able to be associated with a gene name.

| Gene Name | Gene | Wild-Type | *myb28-1$^{ge}$* ($Log^2$ fold change) | *myb28-2$^{ge}$* ($Log^2$ fold change) | FDR | AveExpr | P value |
|---|---|---|---|---|---|---|---|
| thioredoxin-dependent peroxidase 1 | Bo6g099630.1 | 0 | *myb28-2$^{ge}$* | -12.9037 | 2.70E-246 | 7.181706 | 9.13E-251 |
| MAM3 | Bo2g161100.1 | 0 | *myb28-2$^{ge}$* | -12.9141 | 2.48E-261 | 6.299095 | 4.20E-266 |
| Glycosyltransferase | Bo4g049480.1 | 0 | -5.70637 | -5.7818 | 8.42E-35 | 5.997514 | 3.27E-38 |
| BCAT4 | Bo5g113720.1 | 0 | -5.67018 | -5.51459 | 1.04E-39 | 5.586913 | 2.81E-43 |
| MAM3 | Bo7g098000.1 | 0 | -14.1275 | -11.0513 | 1.09E-144 | 4.93904 | 1.10E-148 |
| IMDH3 | Bo2g011730.1 | 0 | -8.58062 | -8.84726 | 2.34E-216 | 4.922201 | 1.18E-220 |
| IPMI2 | Bo4g018590.1 | 0 | -8.25419 | -6.27196 | 6.98E-197 | 4.843612 | 4.71E-201 |
| SUR1 | Bo7g003330.1 | 0 | -3.4525 | -3.33769 | 4.68E-32 | 6.903317 | 1.98E-35 |
| HMT3 | Bo5g094570.1 | 0 | -8.01037 | -7.8182 | 1.63E-143 | 4.659669 | 1.93E-147 |
| CYP79F1 | Bo5g021810.1 | 0 | -10.9654 | -11.8883 | 2.41E-177 | 4.501945 | 2.04E-181 |
| GSTF11 | Bo5g150180.1 | 0 | -3.59728 | -3.54889 | 3.80E-27 | 5.463329 | 1.93E-30 |
| CYP83A1 | Bo4g130780.1 | 0 | -2.60688 | -2.70385 | 2.61E-39 | 8.112961 | 7.49E-43 |
| GSL-OH | Bo4g173530.1 | 0 | -3.44389 | -3.32622 | 2.39E-52 | 5.081223 | 5.25E-56 |
| CYTB5-C | Bo4g007080.1 | 0 | -2.45367 | -2.3517 | 2.22E-14 | 5.109775 | 1.65E-17 |
| SOT17 | Bo5g025610.1 | 0 | -1.9445 | -2.17928 | 0.00021 | 5.202776 | 2.97E-07 |
| CYTB5-C | Bo4g197240.1 | 0 | -1.92843 | -1.98866 | 4.13E-08 | 6.165794 | 3.97E-11 |

| | | | | | | | |
|---|---|---|---|---|---|---|---|
| BAT5 | Bo3g045530.1 | 0 | -10.3621 | -12.2884 | 6.83E-136 | 3.107902 | 9.23E-140 |
| SUR1 | Bo11842s010.1 | 0 | -3.62495 | -3.12312 | 2.72E-41 | 3.900679 | 6.88E-45 |
| | Bo2g127110.1 | 0 | 1.912137 | 1.913905 | 5.29E-10 | 6.793743 | 4.56E-13 |
| SBP2 | Bo8g036770.1 | 0 | -1.95929 | -1.91696 | 2.98E-21 | 6.2186 | 1.76E-24 |
| AK3 | Bo2g041340.1 | 0 | -12.073 | -12.073 | 9.75E-100 | 2.895043 | 1.65E-103 |
| DJ1F | Bo6g068170.1 | 0 | -1.78524 | -1.91861 | 2.72E-16 | 6.640597 | 1.83E-19 |
| UGT74B1 | Bo5g041080.1 | 0 | -1.70813 | -1.55852 | 3.05E-06 | 5.956101 | 3.61E-09 |
| | Bo4g191120.1 | 0 | -2.55034 | -3.40323 | 3.84E-06 | 3.410617 | 4.60E-09 |
| RID1 | Bo2g028590.1 | 0 | -5.75434 | -6.26855 | 7.13E-14 | 2.451008 | 5.42E-17 |
| BAT5 | Bo9g094080.1 | 0 | -11.8041 | -11.8041 | 2.09E-103 | 2.627886 | 3.18E-107 |
| | Bo8g102270.1 | 0 | -1.71803 | -1.63782 | 5.25E-11 | 5.619925 | 4.26E-14 |
| SOT18 | Bo6g118360.1 | 0 | -3.08215 | -3.07456 | 3.91E-37 | 3.385955 | 1.32E-40 |
| APK2 | Bo7g118850.1 | 0 | -1.54535 | -1.50566 | 0.021903 | 4.809315 | 4.14E-05 |
| BCAT4 | Bo3g073430.1 | 0 | -11.6725 | -11.6725 | 1.35E-72 | 2.499999 | 2.73E-76 |
| GSL-OH | Bo4g173560.1 | 0 | -3.42285 | -3.35756 | 7.75E-36 | 3.086089 | 2.75E-39 |
| LTP2 | Bo4g187660.1 | 0 | -1.25327 | -1.39687 | 0.032346 | 5.557113 | 6.45E-05 |
| UGT73B5 | Bo7g021600.1 | 0 | -2.01895 | -1.85708 | 0.006232 | 3.782309 | 1.08E-05 |
| AOP3 | Bo2g102190.1 | 0 | -11.3872 | -9.57761 | 1.70E-75 | 2.217285 | 3.15E-79 |
| | Bo9g155960.1 | 0 | 1.661092 | 1.673869 | 2E-06 | 4.891681 | 2.30E-09 |
| APK2 | Bo1g002960.1 | 0 | -1.87004 | -2.01722 | 4.64E-08 | 3.812394 | 4.55E-11 |
| IIL1 | Bo00805s020.1 | 0 | -1.30893 | -1.20219 | 2.85E-07 | 8.087462 | 3.04E-10 |
| VDAC1 | Bo5g156300.1 | 0 | -0.42787 | -1.78281 | 0.000415 | 4.421757 | 6.17E-07 |
| | Bo5g127540.1 | 0 | -1.22851 | -1.68058 | 0.009613 | 4.123825 | 1.69E-05 |
| FLS2 | Bo3g103270.1 | 0 | -1.76068 | -1.64145 | 6.09E-09 | 4.012655 | 5.56E-12 |

| | | | | | | | |
|---|---|---|---|---|---|---|---|
| FMO GS-OX2 | Bo9g037180.1 | 0 | -0.97958 | -1.19457 | 0.000187 | 7.557033 | 2.59E-07 |
| Phosphoenolpyruvate carboxylase family protein | Bo5g034350.1 | 0 | -1.50785 | -1.30366 | 2.59E-11 | 4.844296 | 2.05E-14 |
| NIFS3 | Bo7g095040.1 | 0 | -1.29589 | -1.19154 | 0.003586 | 4.606345 | 5.93E-06 |
| SULTR4;1 | Bo3g009240.1 | 0 | -1.23243 | -1.54727 | 1.32E-10 | 4.732707 | 1.12E-13 |
| OASA1 | Bo8g039380.1 | 0 | -1.00937 | -1.08007 | 0.000918 | 7.458539 | 1.41E-06 |
| glutathione S-transferase TAU 20 | Bo6g081630.1 | 0 | -1.00154 | -0.99707 | 1.51E-05 | 9.70326 | 1.86E-08 |
| MS2 | Bo3g057170.1 | 0 | -0.94784 | -1.1654 | 0.000199 | 7.306805 | 2.78E-07 |
| ORP2A | Bo3g164360.1 | 0 | -0.12813 | -1.89597 | 0.02427 | 3.556875 | 4.63E-05 |
| GASA7 | Bo9g023730.1 | 0 | -1.84088 | -2.12565 | 0.005088 | 3.024644 | 8.59E-06 |
| | Bo04741s010.1 | 0 | -6.5725 | -8.14032 | 7.01E-39 | 1.597361 | 2.25E-42 |
| | Bo1g103490.1 | 0 | -3.93449 | -3.86267 | 2.37E-26 | 1.853863 | 1.24E-29 |
| | Bo4g136450.1 | 0 | -1.77881 | -2.67575 | 6.26E-07 | 2.619042 | 6.87E-10 |
| | Bo4g197170.1 | 0 | -1.32817 | -5.10265 | 2.37E-31 | 2.339381 | 1.04E-34 |
| SPL7 | Bo9g154530.1 | 0 | -0.74585 | -1.02962 | 0.01653 | 5.496811 | 2.93E-05 |
| CBR1 | Bo3g012760.1 | 0 | -1.21165 | -1.03455 | 2.83E-07 | 4.607999 | 2.96E-10 |
| PER37 | Bo9g083210.1 | 0 | -1.5914 | -1.16113 | 0.025219 | 3.240626 | 4.85E-05 |
| TS2 | Bo2g079040.1 | 0 | -1.65036 | -1.28184 | 4.87E-09 | 3.391229 | 4.36E-12 |
| | Bo4g025390.1 | 0 | 1.348356 | -7.12754 | 1.48E-06 | 1.570954 | 1.68E-09 |
| | Bo9g174920.1 | 0 | 7.452056 | 6.732199 | 3.01E-39 | 2.169582 | 9.15E-43 |
| | Bo6g081870.1 | 0 | 1.479672 | 1.522083 | 1.86E-07 | 3.617712 | 1.91E-10 |
| MSRB4 | Bo3g048350.1 | 0 | -0.90734 | -1.56099 | 0.017702 | 3.219253 | 3.2E-05 |
| | Bo5g118950.1 | 0 | 5.088027 | 3.940808 | 0.003166 | 1.605828 | 5.19E-06 |
| | Bo07665s010.1 | 0 | 2.9403 | -0.31736 | 5.56E-05 | 1.324788 | 6.95E-08 |
| | Bo04963s010.1 | 0 | -8.30347 | -10.2334 | 4.34E-35 | 1.088703 | 1.61E-38 |
| GSTU23 | Bo6g081660.1 | 0 | -7.02708 | -7.69438 | 1.36E-41 | 1.139449 | 3.20E-45 |

| | | | | | | | |
|---|---|---|---|---|---|---|---|
| | Bo7g009490.1 | 0 | -0.35724 | 2.173892 | 0.001977 | 1.729196 | 3.11E-06 |
| AHL | Bo2g041820.1 | 0 | -1.03959 | -1.00084 | 0.000132 | 4.29212 | 1.76E-07 |
| AOP3 | Bo3g052110.1 | 0 | -5.27615 | -5.05767 | 8.41E-29 | 1.081741 | 4.12E-32 |
| HSK | Bo9g024410.1 | 0 | -0.89916 | -0.71561 | 0.032346 | 4.620226 | 6.44E-05 |
| AOP3 | Bo9g006220.1 | 0 | -9.81919 | -9.81919 | 6.55E-29 | 0.687646 | 3.10E-32 |
| | Bo7g093650.1 | 0 | -1.2589 | 5.250326 | 1.28E-17 | 0.687446 | 8.23E-21 |
| | Bo7g095340.1 | 0 | -0.04039 | 8.882702 | 8.5E-05 | -0.06036 | 1.09E-07 |
| GSTU23 | Bo6g081670.1 | 0 | -7.84096 | -9.77068 | 3.61E-33 | 0.648104 | 1.46E-36 |
| MRSA2 | Bo3g004360.1 | 0 | -6.98043 | -6.24429 | 6.57E-24 | 0.624327 | 3.66E-27 |
| | Bo7g064020.1 | 0 | -1.26464 | -1.20282 | 0.002086 | 2.588125 | 3.31E-06 |
| ADAP | Bo5g021520.1 | 0 | -2.76335 | -2.58408 | 7.93E-10 | 1.093813 | 6.96E-13 |
| | Bo9g011230.1 | 0 | 4.881204 | 0.749145 | 4.90E-18 | 0.531155 | 3.06E-21 |
| | Bo3g053230.1 | 0 | -3.25446 | -4.31609 | 1.10E-08 | 0.537995 | 1.02E-11 |
| | Bo5g081320.1 | 0 | -9.39241 | -9.39241 | 8.34E-30 | 0.288829 | 3.80E-33 |
| | Bo6g026520.1 | 0 | -2.69105 | 0.071009 | 0.00104 | 1.494281 | 1.62E-06 |
| KIPK1 | Bo4g122970.1 | 0 | 8.301671 | 9.288866 | 6.97E-08 | 0.756185 | 7.06E-11 |
| | Bo3g170900.1 | 0 | -4.79768 | -6.61747 | 0.000108 | -0.60238 | 1.40E-07 |
| | Bo1g011100.1 | 0 | -2.51738 | -1.9455 | 0.002147 | 0.787668 | 3.44E-06 |
| | Bo02961s010.1 | 0 | -1.29442 | -8.67924 | 0.00604 | 0.061177 | 1.04E-05 |
| | Bo4g187140.1 | 0 | 3.538704 | 2.561941 | 0.000136 | 0.588945 | 1.84E-07 |
| | Bo1g071050.1 | 0 | -8.50617 | -8.50617 | 2.18E-14 | -0.54346 | 1.59E-17 |
| | Bo2g147800.1 | 0 | -6.98653 | -5.17546 | 3.09E-24 | -0.1109 | 1.67E-27 |
| | Bo00751s090.1 | 0 | -6.77368 | -6.9038 | 7.50E-21 | -0.33379 | 4.56E-24 |
| | Bo1g094560.1 | 0 | -3.90185 | -4.22051 | 1.12E-12 | -0.13852 | 8.72E-16 |
| | Bo2g023740.1 | 0 | -5.13335 | -8.74435 | 2.15E-23 | -0.27815 | 1.23E-26 |

| | | | | | | | |
|---|---|---|---|---|---|---|---|
| MPK8 | Bo5g025110.1 | 0 | 7.788461 | 2.683379 | 0.017702 | -1.13945 | 3.19E-05 |
| APK2 | Bo3g152770.1 | 0 | -4.31819 | -3.98 | 9.03E-15 | -0.22879 | 6.40E-18 |
| AOP3 | Bo9g006240.1 | 0 | -8.45877 | -8.45877 | 1.43E-16 | -0.57689 | 9.40E-20 |
| UPS1 | Bo9g074290.1 | 0 | -2.615 | -3.10748 | 0.000213 | 0.042774 | 3.06E-07 |
| | Bo3g005520.1 | 0 | 3.188599 | 3.595512 | 2.85E-06 | 0.319586 | 3.32E-09 |
| | Bo8g099930.1 | 0 | -8.09618 | -8.09618 | 1.83E-15 | -0.89188 | 1.27E-18 |
| | Bo3g062120.1 | 0 | 2.882803 | 2.893276 | 0.04656 | 0.232128 | 9.43E-05 |
| | Bo4g187150.1 | 0 | 3.13897 | 2.056684 | 0.019705 | -0.27317 | 3.63E-05 |
| | Bo5g122560.1 | 0 | -7.36739 | -7.36739 | 0.000305 | -1.51511 | 4.43E-07 |
| | Bo8g021720.1 | 0 | 3.193258 | 2.872992 | 0.000433 | -0.22226 | 6.51E-07 |
| | Bo9g028940.1 | 0 | 3.191328 | 2.052732 | 0.020035 | -0.66458 | 3.72E-05 |
| | Bo3g041970.1 | 0 | -0.89498 | -4.775 | 0.028557 | -0.93792 | 5.59E-05 |
| | Bo1g080260.1 | 0 | -3.05962 | -3.87069 | 4.77E-08 | -1.08896 | 4.76E-11 |
| | Bo9g174670.1 | 0 | 5.725352 | 4.03617 | 1.15E-08 | -0.94653 | 1.09E-11 |
| BCAT4 | Bo3g074470.1 | 0 | -7.3728 | -7.3728 | 6.78E-11 | -1.46786 | 5.61E-14 |
| NHL25 | Bo6g057620.1 | 0 | -5.4926 | -2.93384 | 8.23E-05 | -1.27985 | 1.04E-07 |
| | Bo3g021600.1 | 0 | 7.221399 | 6.954608 | 1.13E-06 | -0.87707 | 1.26E-09 |
| | Bo2g131800.1 | 0 | -4.67171 | -6.60718 | 0.036629 | -2.03057 | 7.36E-05 |
| MAM3 | Bo2g161080.1 | 0 | -6.66403 | -6.66403 | 0.000168 | -1.97291 | 2.30E-07 |
| | Bo9g082100.1 | 0 | -3.79909 | -7.00926 | 4.33E-06 | -1.68297 | 5.27E-09 |
| MAM3 | Bo2g161090.1 | 0 | -6.4971 | -6.4971 | 0.003629 | -2.08146 | 6.07E-06 |
| | Bo7g032910.1 | 0 | -6.8176 | -6.8176 | 6.12E-07 | -1.8869 | 6.61E-10 |
| | Bo3g068230.1 | 0 | 1.865561 | -3.18649 | 0.018036 | -1.60264 | 3.29E-05 |
| | Bo03247s010.1 | 0 | 0.597253 | -5.94504 | 0.027465 | -1.58317 | 5.33E-05 |
| | Bo6g089280.1 | 0 | -6.55457 | -6.55457 | 0.000117 | -2.05079 | 1.55E-07 |

| | | | | | | | |
|---|---|---|---|---|---|---|---|
| | Bo1g103480.1 | 0 | -4.67613 | -2.90669 | 0.020142 | -1.8748 | 3.78E-05 |
| | Bo3g155610.1 | 0 | -6.21394 | -6.21394 | 0.005628 | -2.25896 | 9.6E-06 |
| | Bo9g088590.1 | 0 | -6.58054 | -3.07609 | 0.000729 | -1.93905 | 1.11E-06 |
| | Bo5g126320.1 | 0 | -6.23242 | -6.23242 | 0.000358 | -2.278 | 5.26E-07 |
| | Bo4g101510.1 | 0 | -5.90179 | -5.90179 | 0.002804 | -2.45565 | 4.54E-06 |

**Supplementary Table 5**

All significantly differentially expressed genes (p<0.05) in the generative leaf tissue of both mutant lines, as normalised to wild type, including False Discovery rate correction (FDR), average expression and p values. Note: not all differentially expressed genes (denoted by gene IDs) were able to be associated with a gene name.

| Gene Names | Gene | Wilf-Type | *myb28-1[ge]* (Log2 fold change) | *myb28-2[ge]* (Log2 fold change) | FDR | AveExpr | P value |
|---|---|---|---|---|---|---|---|
| RAB18 | Bo9g022050.1 | 0 | 0.89577676 | 6.019235 | 0.021088 | 7.980402 | 5.66E-05 |
| thioredoxin-dependent peroxidase 1 | Bo6g099630.1 | 0 | -6.6454628 | -6.45636 | 4.60E-15 | 6.926247 | 2.02E-18 |
| Non-specific lipid-transfer protein | Bo2g026820.1 | 0 | 0.35082042 | 5.326064 | 0.000194 | 7.768215 | 2.55E-07 |
| MAM3 | Bo2g161100.1 | 0 | -7.0932514 | -6.57714 | 6.29E-19 | 6.19477 | 1.59E-22 |
| Bidirectional sugar transporter SWEET | Bo2g011110.1 | 0 | 2.82648359 | 10.19711 | 6.42E-05 | 5.984863 | 7.44E-08 |
| Glycosyltransferase | Bo4g049480.1 | 0 | -5.6786521 | -5.86969 | 1.29E-45 | 5.997019 | 2.18E-50 |
| MAM3 | Bo7g098000.1 | 0 | -7.3082881 | -7.03608 | 3.84E-25 | 5.599144 | 2.59E-29 |
| Bidirectional sugar transporter SWEET | Bo9g168970.1 | 0 | 1.35494189 | 8.419225 | 6.42E-05 | 5.226401 | 7.48E-08 |
| BCAT4 | Bo5g113720.1 | 0 | -5.2088117 | -4.46708 | 1.10E-19 | 5.722773 | 2.41E-23 |
| CYP81F1 | Bo1g003710.1 | 0 | 0.13042905 | 8.165615 | 6.8E-05 | 4.823771 | 8.03E-08 |
| CYP79F1 | Bo5g021810.1 | 0 | -5.5418323 | -5.56844 | 7.17E-10 | 5.226679 | 5.20E-13 |
|  | Bo9g078540.1 | 0 | 0.99626826 | 3.842131 | 0.01497 | 6.297929 | 3.69E-05 |
| GSTF11 | Bo5g150180.1 | 0 | -5.7441758 | -5.42485 | 7.13E-36 | 5.073774 | 2.41E-40 |
| EXLB1 | Bo1g053030.1 | 0 | 4.695319 | 13.74309 | 9.51E-05 | 4.576306 | 1.19E-07 |
| CYP83A1 | Bo4g130780.1 | 0 | -2.2439921 | -2.81121 | 0.000403 | 8.422643 | 5.79E-07 |

| | | | | | | | |
|---|---|---|---|---|---|---|---|
| SUR1 | Bo7g003330.1 | 0 | -3.7550978 | -3.42014 | 6.74E-27 | 6.293079 | 3.41E-31 |
| IPMI2 | Bo4g018590.1 | 0 | -4.9518095 | -4.84714 | 7.91E-13 | 5.015098 | 4.27E-16 |
| MED37C | Bo3g065020.1 | 0 | 0.62985762 | 3.469315 | 0.015919 | 5.605088 | 4.03E-05 |
| IMDH3 | Bo2g011730.1 | 0 | -4.62587 | -4.60775 | 1.21E-06 | 4.820617 | 1.02E-09 |
| GSL-OH | Bo4g173530.1 | 0 | -4.8489669 | -5.26875 | 7.55E-24 | 4.670581 | 1.02E-27 |
| NSP5 | Bo2g149730.1 | 0 | 1.6506806 | 5.426776 | 0.006632 | 4.255287 | 1.39E-05 |
| ATHB-7 | Bo4g197260.1 | 0 | 0.86417433 | 4.871076 | 0.024106 | 4.148892 | 6.76E-05 |
| HMT3 | Bo5g094570.1 | 0 | -4.2615342 | -4.72853 | 4.72E-18 | 4.685425 | 1.52E-21 |
| LTI65 | Bo2g047970.1 | 0 | 2.14671303 | 8.954684 | 9.02E-05 | 3.7555 | 1.11E-07 |
| | Bo3g170670.1 | 0 | 0.57565463 | 4.97428 | 0.000761 | 3.940833 | 1.2E-06 |
| AK3 | Bo2g041340.1 | 0 | -5.7964653 | -5.70397 | 1.18E-14 | 4.155922 | 5.59E-18 |
| FTSH6 | Bo9g164320.1 | 0 | 0.55711805 | 4.793569 | 0.005192 | 3.771564 | 1.04E-05 |
| P5CSA | Bo4g188600.1 | 0 | -0.3056296 | 3.551761 | 0.008132 | 4.050158 | 1.74E-05 |
| CYTB5-C | Bo4g197240.1 | 0 | -3.070141 | -2.71953 | 7.1E-06 | 5.145131 | 6.96E-09 |
| MED37C | Bo1g139130.1 | 0 | 0.31440384 | 4.489335 | 0.043179 | 3.595029 | 0.00014 |
| EGY3 | Bo00831s080.1 | 0 | -0.342561 | 3.379073 | 0.005511 | 3.96059 | 1.13E-05 |
| | Bo2g127110.1 | 0 | 2.30203194 | 2.109571 | 1.11E-05 | 7.232329 | 1.16E-08 |
| | Bo9g175850.1 | 0 | 1.05589578 | 7.632693 | 0.029846 | 3.188024 | 8.77E-05 |
| | Bo4g173400.1 | 0 | 3.48497146 | -0.3913 | 0.008028 | 3.862906 | 1.71E-05 |
| | Bo6g087900.1 | 0 | -0.6795353 | 3.961936 | 0.021355 | 3.430478 | 5.88E-05 |
| SWEET15 | Bo3g009030.1 | 0 | 4.21445139 | 8.636767 | 0.037536 | 3.206157 | 0.000117 |

| | | | | | | | |
|---|---|---|---|---|---|---|---|
| SSL9 | Bo00927s010.1 | 0 | -0.0834466 | 3.63372 | 0.005665 | 3.667179 | 1.17E-05 |
| ABCF2 | Bo9g173670.1 | 0 | 1.55527371 | 5.033788 | 0.031688 | 3.359538 | 9.52E-05 |
| AAC3 | Bo8g054310.1 | 0 | 0.6284917 | 5.980301 | 0.001773 | 3.098644 | 3.05E-06 |
| TSPO | Bo01463s030.1 | 0 | 1.74094337 | 6.260866 | 0.026863 | 3.067716 | 7.76E-05 |
| WSCP | Bo02822s010.1 | 0 | -2.5330301 | -9.68798 | 0.023716 | 3.384694 | 6.61E-05 |
| FTSH6 | Bo9g164300.1 | 0 | 1.83022533 | 5.894795 | 0.003607 | 2.935115 | 6.88E-06 |
| | Bo5g122890.1 | 0 | 0.69780124 | 9.828433 | 0.000342 | 2.65698 | 4.86E-07 |
| | Bo3g171010.1 | 0 | 1.4956457 | 6.502288 | 0.021355 | 2.690767 | 5.87E-05 |
| GSTF9 | Bo4g173610.1 | 0 | -1.7162767 | -1.82752 | 0.005054 | 7.110865 | 9.98E-06 |
| UGT74B1 | Bo5g041080.1 | 0 | -2.274577 | -2.52225 | 3.37E-09 | 5.196983 | 2.50E-12 |
| BCAT4 | Bo3g073430.1 | 0 | -7.060444 | -6.35473 | 2.06E-21 | 3.11146 | 3.82E-25 |
| SUR1 | Bo11842s010.1 | 0 | -4.4985163 | -4.00847 | 1.48E-24 | 3.367794 | 1.49E-28 |
| FTSH6 | Bo04609s010.1 | 0 | 2.23114144 | 6.747303 | 0.03081 | 2.452789 | 9.16E-05 |
| MT4B | Bo3g042480.1 | 0 | 0.70604926 | 9.471535 | 0.000101 | 2.293808 | 1.27E-07 |
| CYTB5-C | Bo4g007080.1 | 0 | -2.9196085 | -3.29874 | 1.33E-18 | 3.952128 | 3.82E-22 |
| MIOX1 | Bo5g019730.1 | 0 | 2.57954583 | 4.643021 | 0.000437 | 3.084708 | 6.65E-07 |
| BAT5 | Bo3g045530.1 | 0 | -4.5897941 | -5.0115 | 3.60E-11 | 3.125927 | 2.37E-14 |
| | Bo9g177270.1 | 0 | 4.1942885 | 0.389683 | 0.046791 | 2.810934 | 0.000153 |
| UGT73B5 | Bo7g021600.1 | 0 | -2.594324 | -2.20761 | 1.20E-08 | 4.450583 | 9.11E-12 |
| | Bo3g055630.1 | 0 | -0.155465 | 7.57505 | 0.000903 | 2.170428 | 1.45E-06 |

| | | | | | | | |
|---|---|---|---|---|---|---|---|
| SBP2 | Bo8g036770.1 | 0 | -1.8967991 | -1.84848 | 9.15E-07 | 6.412817 | 7.57E-10 |
| RID1 | Bo2g028590.1 | 0 | -7.7546061 | -7.48568 | 1.10E-16 | 2.784029 | 4.08E-20 |
| DTX43 | Bo1g145050.1 | 0 | 1.52872384 | 4.955496 | 0.010203 | 2.297545 | 2.31E-05 |
| GAD1 | Bo9g161390.1 | 0 | -0.6066514 | -5.10489 | 0.035416 | 2.964923 | 0.000109 |
| BCAT2 | Bo8g110790.1 | 0 | 0.63477297 | 6.516702 | 0.01294 | 2.028681 | 3.06E-05 |
| | Bo7g114510.1 | 0 | 0.81164597 | 4.074088 | 0.010669 | 2.42284 | 2.43E-05 |
| CRK7 | Bo4g133160.1 | 0 | -1.028417 | 2.238084 | 0.002689 | 2.982558 | 4.81E-06 |
| DJ1F | Bo6g068170.1 | 0 | -1.6335127 | -1.6089 | 0.000309 | 7.544022 | 4.23E-07 |
| | Bo6g104060.1 | 0 | -1.1825671 | -2.63691 | 0.001773 | 4.238526 | 3.04E-06 |
| BAT5 | Bo9g094080.1 | 0 | -5.0532465 | -4.75121 | 1.09E-10 | 2.662263 | 7.74E-14 |
| GSL-OH | Bo4g173560.1 | 0 | -4.7375613 | -5.34721 | 5.06E-15 | 2.662498 | 2.31E-18 |
| APK1 | Bo9g022580.1 | 0 | -1.4934486 | -1.44616 | 0.020157 | 6.011563 | 5.38E-05 |
| | Bo01233s030.1 | 0 | -0.5272035 | 4.904662 | 0.001277 | 1.7586 | 2.11E-06 |
| SOT17 | Bo5g025610.1 | 0 | -2.6971631 | -3.24514 | 1.55E-11 | 3.293144 | 9.43E-15 |
| Phosphoenolpyruvate carboxylase family protein | Bo5g034350.1 | 0 | -1.9308943 | -1.82766 | 2.5E-05 | 4.41423 | 2.74E-08 |
| FLS2 | Bo3g103270.1 | 0 | -2.1713074 | -1.11795 | 0.003484 | 4.343025 | 6.47E-06 |
| APK2 | Bo1g002960.1 | 0 | -3.2642501 | -3.84551 | 1.33E-12 | 2.958638 | 7.43E-16 |
| SOT18 | Bo6g118360.1 | 0 | -3.1168414 | -2.67226 | 6.06E-14 | 3.305718 | 3.07E-17 |

| | | | | | | | |
|---|---|---|---|---|---|---|---|
| Lactoylglutathione lyase / glyoxalase I family protein | Bo2g096220.1 | 0 | 1.3607316 | 6.927535 | 0.018174 | 1.583906 | 4.76E-05 |
| DREB1F | Bo01335s010.1 | 0 | 4.94620033 | 0.321194 | 1.08E-05 | 1.967745 | 1.12E-08 |
| | Bo4g030290.1 | 0 | 0.68600032 | 3.932522 | 0.024529 | 1.790529 | 6.92E-05 |
| POX1 | Bo2g149510.1 | 0 | 1.32933066 | 3.154791 | 0.016046 | 2.478253 | 4.12E-05 |
| | Bo3g045840.1 | 0 | -1.344985 | 5.56086 | 0.024587 | 1.336428 | 6.99E-05 |
| APK2 | Bo7g118850.1 | 0 | -1.9733898 | -2.51853 | 0.001071 | 3.351605 | 1.74E-06 |
| | Bo6g098930.1 | 0 | 4.45495552 | -0.32919 | 3.92E-07 | 1.910727 | 3.11E-10 |
| | Bo5g016960.1 | 0 | 1.49963399 | 6.902228 | 0.003579 | 1.138757 | 6.77E-06 |
| AOP3 | Bo3g052110.1 | 0 | -4.6426063 | -5.16382 | 3.41E-24 | 2.101627 | 4.03E-28 |
| INT1 | Bo4g019720.1 | 0 | 1.78850525 | 6.451081 | 0.01264 | 1.101044 | 2.97E-05 |
| | Bo4g191120.1 | 0 | -5.2082246 | -4.00562 | 5.07E-06 | 1.929704 | 4.80E-09 |
| | Bo6g119910.1 | 0 | 1.94995321 | -0.6528 | 0.024587 | 2.494506 | 7.02E-05 |
| WAK3 | Bo5g033140.1 | 0 | 1.19671638 | 5.64921 | 0.016046 | 1.08059 | 4.1E-05 |
| AHL | Bo2g041820.1 | 0 | -1.6536626 | -1.49906 | 0.000319 | 4.163174 | 4.47E-07 |
| MRSA2 | Bo3g004360.1 | 0 | -3.6694705 | -4.9324 | 8.96E-23 | 2.034639 | 1.51E-26 |
| | Bo4g197170.1 | 0 | -1.171927 | -3.5354 | 2.17E-06 | 2.531722 | 1.94E-09 |
| TAAC | Bo3g001430.1 | 0 | -1.5439232 | -1.57398 | 0.043023 | 3.471386 | 0.000138 |
| ORP2A | Bo3g164360.1 | 0 | 0.11881485 | -2.27327 | 1.87E-05 | 3.06248 | 2.02E-08 |
| | Bo4g030270.1 | 0 | 1.93865705 | 6.895797 | 0.000429 | 0.644043 | 6.30E-07 |
| | Bo4g136450.1 | 0 | -1.4092596 | -2.91138 | 0.000197 | 2.572918 | 2.62E-07 |

| | | | | | | | |
|---|---|---|---|---|---|---|---|
| beta carbonic anhydrase 3 | Bo7g055990.1 | 0 | -5.1543246 | -5.08309 | 0.009826 | 0.92505 | 2.21E-05 |
| SOT14 | Bo2g007480.1 | 0 | -0.6050605 | -4.24831 | 0.042742 | 1.539801 | 0.000136 |
| | Bo9g011230.1 | 0 | 4.37934419 | 0.73181 | 2.75E-11 | 1.454493 | 1.76E-14 |
| | Bo05195s010.1 | 0 | -9.4573495 | -9.45735 | 0.000319 | 0.366164 | 4.44E-07 |
| ICMEL2 | Bo5g152640.1 | 0 | 1.82106908 | 7.325893 | 0.018671 | 0.2773 | 4.92E-05 |
| AOP3 | Bo2g102190.1 | 0 | -10.099435 | -10.0994 | 1.30E-24 | 0.971935 | 1.10E-28 |
| DREB1F | Bo8g108050.1 | 0 | 6.21197664 | 0.530967 | 0.000157 | 0.515927 | 2.04E-07 |
| | Bo1g103490.1 | 0 | -5.3103759 | -5.58474 | 3.54E-16 | 1.145082 | 1.43E-19 |
| | Bo8g101690.1 | 0 | 3.88662384 | 0.959388 | 0.004772 | 0.824787 | 9.35E-06 |
| | Bo9g174920.1 | 0 | 6.14226651 | 5.139158 | 1.84E-13 | 1.591232 | 9.61E-17 |
| HSP22 | Bo3g044590.1 | 0 | 1.86426675 | 8.958969 | 0.002008 | -0.06685 | 3.53E-06 |
| XTH23 | Bo1g040530.1 | 0 | 4.20742852 | 1.280327 | 0.008325 | 0.610617 | 1.81E-05 |
| UGT74D1 | Bo3g026420.1 | 0 | -2.4162375 | -3.0291 | 0.040115 | 1.343422 | 0.000126 |
| MPK8 | Bo5g025110.1 | 0 | 6.93025971 | 1.494456 | 0.009172 | -0.10127 | 2.01E-05 |
| | Bo5g081320.1 | 0 | -6.0706835 | -7.18104 | 1.99E-23 | 0.731828 | 3.03E-27 |
| | Bo04741s010.1 | 0 | -5.9052784 | -6.49738 | 8.06E-18 | 0.56565 | 2.72E-21 |
| | Bo6g119090.1 | 0 | -0.2625804 | -4.13705 | 0.009341 | 1.189853 | 2.07E-05 |
| SRO5 | Bo2g164210.1 | 0 | 4.58972262 | 0.897199 | 0.005192 | 0.329794 | 1.04E-05 |
| | Bo9g026210.1 | 0 | 4.1379749 | -0.85183 | 0.046562 | 0.305277 | 0.000152 |

| | | | | | | | |
|---|---|---|---|---|---|---|---|
| ORG3 | Bo01495s010.1 | 0 | 3.98814668 | -0.59056 | 0.01552 | 0.267949 | 3.88E-05 |
| | Bo9g140620.1 | 0 | 6.3404998 | 3.061345 | 0.024614 | 0.178263 | 7.07E-05 |
| XTH25 | Bo2g030320.1 | 0 | -3.9419342 | 4.757947 | 0.000771 | -0.2646 | 1.22E-06 |
| | Bo7g093650.1 | 0 | 1.48638629 | 7.288988 | 2.30E-18 | 0.243438 | 7.00E-22 |
| | Bo4g078140.1 | 0 | 1.51579991 | 6.386317 | 0.001953 | -0.52758 | 3.4E-06 |
| NAC081 | Bo02109s010.1 | 0 | -6.0934513 | -9.00894 | 3.21E-06 | -0.05043 | 2.98E-09 |
| | Bo5g024020.1 | 0 | 2.87247447 | 4.852529 | 0.032959 | -0.1342 | 9.96E-05 |
| APK2 | Bo3g152770.1 | 0 | -5.2981459 | -4.4768 | 2.61E-11 | 0.373378 | 1.63E-14 |
| GSTU23 | Bo6g081660.1 | 0 | -7.2786776 | -9.09682 | 7.03E-17 | 0.027386 | 2.49E-20 |
| | Bo2g097420.1 | 0 | 5.18127327 | 0.8148 | 0.043023 | -0.06053 | 0.000139 |
| | Bo5g071320.1 | 0 | 2.40804089 | -1.86058 | 0.014829 | 0.122775 | 3.61E-05 |
| | Bo2g023740.1 | 0 | -9.0958081 | -7.24697 | 8.24E-20 | 0.023424 | 1.67E-23 |
| GSTU23 | Bo6g081670.1 | 0 | -7.0749987 | -8.90771 | 2.08E-15 | -0.14585 | 8.77E-19 |
| | Bo7g080240.1 | 0 | 3.21248568 | 8.406338 | 1.38E-06 | -0.53486 | 1.19E-09 |
| | Bo7g101140.1 | 0 | 2.86277537 | 8.070564 | 0.021355 | -0.82881 | 5.85E-05 |
| | Bo2g147800.1 | 0 | -5.7015217 | -7.31281 | 1.30E-18 | 0.110266 | 3.50E-22 |
| ERF025 | Bo9g104780.1 | 0 | 8.35025017 | 1.843935 | 1.31E-05 | -0.53897 | 1.40E-08 |
| | Bo9g066910.1 | 0 | 2.15975535 | 3.650586 | 0.033876 | 0.047591 | 0.000103 |

| | | | | | | | |
|---|---|---|---|---|---|---|---|
| | Bo3g053230.1 | 0 | -3.3888521 | -5.17331 | 8.87E-05 | -0.2056 | 1.08E-07 |
| DREB1F | Bo5g016750.1 | 0 | 6.48674493 | -1.97391 | 2.37E-06 | -0.46773 | 2.16E-09 |
| AGL28 | Bo4g165540.1 | 0 | 0 | 7.892573 | 0.004221 | -1.00587 | 8.2E-06 |
| | Bo4g025390.1 | 0 | 1.45526063 | -7.36101 | 6.29E-06 | 0.199308 | 6.05E-09 |
| KIPK1 | Bo4g122970.1 | 0 | 6.42470109 | 6.709992 | 8.02E-06 | 0.484967 | 7.99E-09 |
| | Bo8g066810.1 | 0 | 2.66444975 | 7.844995 | 0.011769 | -1.01747 | 2.74E-05 |
| | Bo8g032010.1 | 0 | -1.9799588 | 5.797663 | 0.013219 | -1.08583 | 3.15E-05 |
| AOP3 | Bo9g006220.1 | 0 | -7.093919 | -8.90736 | 2.33E-19 | -0.15332 | 5.52E-23 |
| OEP162 | Bo7g104260.1 | 0 | -0.155724 | 5.745721 | 0.040558 | -1.08437 | 0.000128 |
| | Bo4g187150.1 | 0 | 5.48984115 | 5.827528 | 0.031116 | -0.34873 | 9.3E-05 |
| | Bo04963s010.1 | 0 | -6.1027679 | -8.6647 | 2.10E-16 | -0.36758 | 8.15E-20 |
| BHLH101 | Bo3g002630.1 | 0 | 5.40631496 | 1.031554 | 0.001457 | -0.69409 | 2.44E-06 |
| SYP21 | Bo9g151640.1 | 0 | -7.6129512 | -7.61295 | 0.003579 | -1.27292 | 6.74E-06 |
| | Bo3g173610.1 | 0 | 0 | 7.515908 | 0.011769 | -1.31213 | 2.74E-05 |
| | Bo3g085440.1 | 0 | -1.934558 | -3.98867 | 0.029789 | 0.068464 | 8.7E-05 |
| | Bo1g094560.1 | 0 | -5.8411818 | -8.4025 | 3.20E-14 | -0.59648 | 1.57E-17 |
| MSD2 | Bo8g090780.1 | 0 | 7.76061274 | 5.657053 | 0.010697 | -0.88734 | 2.46E-05 |

| | | | | | | | |
|---|---|---|---|---|---|---|---|
| | Bo8g076250.1 | 0 | 3.64345465 | -2.10074 | 0.01497 | -1.06594 | 3.67E-05 |
| | Bo00640s070.1 | 0 | 3.10686016 | 7.452528 | 0.002948 | -1.31302 | 5.38E-06 |
| SWEET14 | Bo1g039530.1 | 0 | 2.01401896 | 4.20562 | 0.028721 | -0.73456 | 8.34E-05 |
| | Bo1g071050.1 | 0 | -6.7082393 | -8.22896 | 4.37E-11 | -0.77448 | 2.96E-14 |
| CPK14 | Bo4g014560.1 | 0 | 5.61521081 | 7.343779 | 0.036232 | -1.14431 | 0.000112 |
| | Bo3g170900.1 | 0 | -2.4397144 | -5.94719 | 0.015919 | -0.93995 | 4.01E-05 |
| | Bo6g106530.1 | 0 | 5.82933181 | 0.674999 | 0.000437 | -1.07622 | 6.59E-07 |
| ORG2 | Bo4g112600.1 | 0 | 5.70347686 | -0.14572 | 0.001079 | -1.16688 | 1.77E-06 |
| | Bo8g090360.1 | 0 | -5.5047188 | -7.36918 | 0.042742 | -1.49965 | 0.000136 |
| | Bo3g062120.1 | 0 | 3.60179197 | 3.697212 | 0.006166 | -0.06682 | 1.28E-05 |
| BCAT4 | Bo3g074470.1 | 0 | -8.1119364 | -5.47609 | 1.70E-12 | -0.84919 | 9.78E-16 |
| UGT74F2 | Bo4g021060.1 | 0 | 3.10426139 | 7.194603 | 0.002323 | -1.50465 | 4.12E-06 |
| | Bo6g089280.1 | 0 | -7.9924283 | -7.99243 | 4.79E-12 | -0.97456 | 2.83E-15 |
| | Bo9g082100.1 | 0 | -7.9661411 | -5.33037 | 8.64E-11 | -0.96839 | 5.98E-14 |
| HEMA3 | Bo5g016620.1 | 0 | -1.812781 | -3.4014 | 0.00543 | -0.30235 | 1.1E-05 |
| | Bo3g021600.1 | 0 | 7.82153391 | 7.411683 | 1.6E-06 | -0.36959 | 1.40E-09 |
| | Bo1g139190.1 | 0 | -3.7750871 | -2.96143 | 0.033876 | -1.10794 | 0.000104 |

| | | | | | | | |
|---|---|---|---|---|---|---|---|
| | Bo3g005520.1 | 0 | 6.18349101 | 6.274392 | 2.20E-07 | -0.28328 | 1.71E-10 |
| | Bo9g088590.1 | 0 | -7.5398929 | -2.32142 | 0.000437 | -1.12461 | 6.59E-07 |
| MKK5 | Bo3g077410.1 | 0 | -1.8341256 | -2.68491 | 0.008325 | -0.33821 | 1.8E-05 |
| | Bo9g174670.1 | 0 | 7.85679688 | 6.256865 | 6.31E-07 | -0.69226 | 5.11E-10 |
| SRO4 | Bo8g079760.1 | 0 | 5.16148983 | -0.12528 | 0.000757 | -1.49633 | 1.18E-06 |
| | Bo8g021720.1 | 0 | 3.35122947 | 3.642077 | 0.001638 | -0.35679 | 2.77E-06 |
| | Bo9g120850.1 | 0 | -1.5377643 | -5.05805 | 0.007137 | -0.83237 | 1.51E-05 |
| | Bo7g032970.1 | 0 | 0 | 6.707861 | 0.003212 | -1.91439 | 5.91E-06 |
| | Bo2g131800.1 | 0 | -3.6999489 | -7.2665 | 0.002704 | -1.49226 | 4.89E-06 |
| | Bo2g159530.1 | 0 | 5.33500954 | 2.613953 | 0.016953 | -1.27937 | 4.38E-05 |
| | Bo2g140690.1 | 0 | 1.44074429 | -2.49173 | 0.021467 | -0.96784 | 5.94E-05 |
| | Bo1g094280.1 | 0 | 0 | 6.925465 | 0.000256 | -1.7635 | 3.46E-07 |
| NHL25 | Bo6g057620.1 | 0 | -4.0213733 | -4.4118 | 9.86E-06 | -1.21756 | 9.99E-09 |
| | Bo4g137320.1 | 0 | 6.83927247 | 2.059223 | 0.029983 | -1.84471 | 8.86E-05 |
| | Bo9g134900.1 | 0 | 5.08213958 | 6.713617 | 0.013921 | -1.62475 | 3.34E-05 |
| | Bo1g080260.1 | 0 | -4.6319527 | -4.48944 | 7.39E-05 | -1.5938 | 8.86E-08 |
| TMA7 | Bo5g126470.1 | 0 | -2.8664848 | -3.58346 | 0.014398 | -1.48666 | 3.48E-05 |

| | | | | | | | |
|---|---|---|---|---|---|---|---|
| | Bo8g099930.1 | 0 | -6.9222768 | -6.92228 | 2.97E-05 | -1.83384 | 3.31E-08 |
| | Bo4g151870.1 | 0 | -6.8366969 | -4.97933 | 4.31E-05 | -1.86593 | 4.88E-08 |
| MAM3 | Bo2g161090.1 | 0 | -6.7405661 | -4.88951 | 0.000106 | -1.92727 | 1.37E-07 |
| | Bo9g090760.1 | 0 | -6.514576 | -6.51458 | 0.000631 | -2.10208 | 9.70E-07 |
| | Bo9g090750.1 | 0 | -6.524064 | -6.52406 | 0.000407 | -2.09893 | 5.91E-07 |
| | Bo3g123500.1 | 0 | -3.594794 | -3.48414 | 0.019357 | -1.88767 | 5.13E-05 |
| | Bo7g021590.1 | 0 | -2.1732129 | -6.32842 | 0.048796 | -2.03683 | 0.000161 |
| PHT1-4 | Bo07610s010.1 | 0 | 5.99955622 | 1.04E-07 | 0.01497 | -2.38963 | 3.72E-05 |
| | Bo3g155610.1 | 0 | -6.2249612 | -4.37986 | 0.009409 | -2.25961 | 2.1E-05 |
| | Bo7g032910.1 | 0 | -6.2096816 | -3.55885 | 0.039285 | -2.23072 | 0.000123 |
| TET12 | Bo2g161070.1 | 0 | -6.0962038 | -3.45421 | 0.021313 | -2.29789 | 5.76E-05 |
| | Bo1g103480.1 | 0 | -5.9753177 | -5.97532 | 0.004177 | -2.43821 | 8.04E-06 |
| | Bo5g126320.1 | 0 | -5.9378774 | -5.93788 | 0.017046 | -2.46895 | 4.43E-05 |

**Supplementary Table 6**

All significantly differentially expressed genes (p<0.05) in the floret material of both mutant lines, as normalised to wild type, including False Discovery rate correction (FDR), average expression and p values. Note: not all differentially expressed genes (denoted by gene IDs) were able to be associated with a gene name.

| Gene Names | Gene | Wild-Type | *myb28-1ge* (Log2 fold change) | *myb28-2ge* (Log2 fold change) | FDR (False Discovery Rate) | AveExpr | P value (Raw P Value) |
|---|---|---|---|---|---|---|---|
| Glycosyltransferase | Bo4g049480.1 | 0 | -4.65277 | -4.71622 | 5.36E-42 | 5.272001 | 1.64E-46 |
| | Bo5g081320.1 | 0 | -8.1242 | -10.0051 | 5.48E-27 | 0.889045 | 3.35E-31 |
| | Bo2g147800.1 | 0 | -7.85158 | -9.74782 | 2.37E-26 | 0.644979 | 2.18E-30 |
| MAM3 | Bo2g161100.1 | 0 | -4.97702 | -4.6956 | 3.99E-26 | 4.74852 | 4.88E-30 |
| glutathione S-transferase F11 | Bo5g150180.1 | 0 | -3.55443 | -3.72746 | 9.97E-25 | 4.697186 | 1.52E-28 |
| MAM3 | Bo7g098000.1 | 0 | -5.87412 | -5.60682 | 1.46E-22 | 4.224402 | 2.68E-26 |
| GSL-OH | Bo4g173530.1 | 0 | -3.11624 | -3.08855 | 8.15E-21 | 4.495668 | 1.74E-24 |
| thioredoxin-dependent peroxidase 1 | Bo6g099630.1 | 0 | -5.93379 | -5.52438 | 2.38E-20 | 6.130602 | 6.56E-24 |
| CYP79F1 | Bo5g021810.1 | 0 | -4.80178 | -4.79768 | 2.38E-20 | 4.004275 | 6.07E-24 |
| | Bo2g023740.1 | 0 | -7.61638 | -5.99066 | 1.28E-19 | 0.432692 | 3.91E-23 |
| BCAT4 | Bo5g113720.1 | 0 | -3.17408 | -3.72849 | 4.08E-19 | 4.951622 | 1.37E-22 |
| CYP83A1 | Bo4g130780.1 | 0 | -2.49371 | -3.18184 | 6.24E-19 | 6.732921 | 2.29E-22 |
| | Bo9g174920.1 | 0 | 7.219799 | 6.485808 | 2.26E-17 | 1.916695 | 8.98E-21 |

| | | | | | | | |
|---|---|---|---|---|---|---|---|
| BAT5 | Bo3g045530.1 | 0 | -4.05462 | -4.12426 | 2.58E-17 | 2.276624 | 1.10E-20 |
| SUR1 | Bo7g003330.1 | 0 | -2.27002 | -2.3851 | 2.57E-16 | 5.920779 | 1.18E-19 |
| BCAT4 | Bo3g073430.1 | 0 | -5.86303 | -4.42673 | 1.64E-15 | 2.315325 | 8.00E-19 |
| AK3 | Bo2g041340.1 | 0 | -4.76147 | -5.08579 | 1.17E-14 | 2.420731 | 6.10E-18 |
| AOP3 | Bo2g102190.1 | 0 | -8.67849 | -7.78361 | 2.19E-14 | -0.34471 | 1.20E-17 |
| IPMI2 | Bo4g018590.1 | 0 | -4.2119 | -3.29382 | 3.68E-14 | 4.681572 | 2.14E-17 |
| CYTB5-C | Bo4g007080.1 | 0 | -2.60089 | -2.94961 | 7.31E-14 | 3.414005 | 4.47E-17 |
| AOP3 | Bo3g052110.1 | 0 | -4.28745 | -3.20858 | 1.05E-12 | 1.004778 | 6.74E-16 |
| IMDH3 | Bo2g011730.1 | 0 | -3.66983 | -3.78824 | 1.83E-12 | 3.903266 | 1.23E-15 |
| GSL-OH | Bo4g173560.1 | 0 | -2.8947 | -2.93596 | 4.16E-11 | 2.549207 | 2.92E-14 |
| | Bo8g099930.1 | 0 | -4.40755 | -4.85315 | 8.21E-11 | 0.826642 | 6.03E-14 |
| KIPK1 | Bo4g122970.1 | 0 | 4.16534 | 4.980866 | 2.72E-10 | 0.833861 | 2.08E-13 |
| | Bo2g127110.1 | 0 | 3.306875 | 3.177187 | 5.19E-09 | 6.594418 | 4.13E-12 |
| BAT5 | Bo9g094080.1 | 0 | -3.90653 | -4.4419 | 2.28E-08 | 1.947237 | 1.90E-11 |
| SOT18 | Bo6g118360.1 | 0 | -2.38592 | -2.19056 | 2.28E-08 | 2.915119 | 1.96E-11 |
| SUR1 | Bo11842s010.1 | 0 | -2.74234 | -2.5673 | 3.50E-08 | 2.91203 | 3.11E-11 |
| HMT3 | Bo5g094570.1 | 0 | -3.24471 | -3.40235 | 3.68E-08 | 3.470788 | 3.38E-11 |

| | | | | | | | |
|---|---|---|---|---|---|---|---|
| | Bo9g082100.1 | 0 | -6.07904 | -6.12708 | 4.23E-08 | -0.92881 | 4.01E-11 |
| | Bo1g103490.1 | 0 | -3.54283 | -4.74715 | 4.59E-08 | 0.107022 | 4.49E-11 |
| | Bo4g025390.1 | 0 | 2.151564 | -2.45739 | 4.03E-07 | 0.521178 | 4.06E-10 |
| | Bo3g062120.1 | 0 | 4.403229 | 4.658832 | 5.24E-07 | 0.301815 | 5.45E-10 |
| CYTB5-C | Bo4g197240.1 | 0 | -1.91326 | -2.31828 | 8.74E-07 | 5.209267 | 9.35E-10 |
| | Bo3g053230.1 | 0 | -5.25492 | -5.65929 | 8.98E-07 | -0.16101 | 9.89E-10 |
| | Bo04741s010.1 | 0 | -4.13283 | -3.52087 | 1.03E-06 | -0.21094 | 1.17E-09 |
| SBP2 | Bo8g036770.1 | 0 | -1.57862 | -1.70776 | 1.22E-06 | 6.324262 | 1.42E-09 |
| | Bo1g123340.1 | 0 | -3.37645 | 2.967185 | 1.26E-06 | 0.586819 | 1.50E-09 |
| RAB18 | Bo2g167630.1 | 0 | -2.43822 | -2.63078 | 1.4E-06 | 2.731664 | 1.71E-09 |
| | Bo1g071050.1 | 0 | -6.59114 | -8.45996 | 1.99E-06 | -0.54251 | 2.49E-09 |
| | Bo4g136450.1 | 0 | -1.55207 | -2.70507 | 4.41E-06 | 2.473391 | 5.66E-09 |
| | Bo3g021600.1 | 0 | 6.880155 | 7.820672 | 7.62E-06 | -0.58058 | 1.00E-08 |
| | Bo4g191120.1 | 0 | -3.5214 | -2.48054 | 7.92E-06 | 1.496089 | 1.07E-08 |
| RID1 | Bo2g028590.1 | 0 | -5.43386 | -4.7698 | 1.55E-05 | 2.616405 | 2.13E-08 |
| APK2 | Bo1g002960.1 | 0 | -1.21828 | -2.58496 | 2.09E-05 | 2.591669 | 2.94E-08 |
| APK2 | Bo3g152770.1 | 0 | -4.10479 | -5.59853 | 2.36E-05 | -0.65276 | 3.39E-08 |

| | | | | | | | |
|---|---|---|---|---|---|---|---|
| MRSA2 | Bo3g004360.1 | 0 | -2.54491 | -3.37714 | 4.59E-05 | 0.494115 | 6.74E-08 |
| HOP3 | Bo2g105000.1 | 0 | -0.52969 | 1.763973 | 4.93E-05 | 5.367744 | 7.39E-08 |
| | Bo9g155960.1 | 0 | 1.580726 | 1.823958 | 0.000202 | 5.473348 | 3.15E-07 |
| | Bo8g102330.1 | 0 | -1.11963 | 1.748506 | 0.000202 | 4.502251 | 3.10E-07 |
| | Bo04963s010.1 | 0 | -3.39799 | -3.82174 | 0.000223 | -0.38392 | 3.69E-07 |
| FLS2 | Bo3g103270.1 | 0 | -1.47699 | -1.34279 | 0.000223 | 4.580898 | 3.55E-07 |
| SOT17 | Bo5g025610.1 | 0 | -1.37771 | -2.2375 | 0.000223 | 3.850066 | 3.75E-07 |
| BCAT4 | Bo3g074470.1 | 0 | -5.83295 | -4.11377 | 0.000223 | -1.10467 | 3.70E-07 |
| | Bo5g122560.1 | 0 | -7.61654 | -7.61654 | 0.000344 | -1.26657 | 5.90E-07 |
| UGT74B1 | Bo5g041080.1 | 0 | -1.30468 | -1.375 | 0.000579 | 5.45855 | 1.01431E-06 |
| | Bo9g174670.1 | 0 | 7.397564 | 5.456138 | 0.000579 | -1.17738 | 1.02747E-06 |
| | Bo7g033020.1 | 0 | -1.02975 | 2.029987 | 0.000866 | 3.647544 | 1.56241E-06 |
| AHA5 | Bo3g100630.1 | 0 | -7.98703 | -3.298 | 0.00094 | -0.83662 | 1.72504E-06 |
| | Bo5g118950.1 | 0 | 4.417191 | 3.873492 | 0.001228 | 1.690163 | 2.28985E-06 |
| | Bo8g013980.1 | 0 | -6.83465 | -0.5192 | 0.001424 | 0.370772 | 2.71154E-06 |
| | Bo3g032000.1 | 0 | -4.42998 | -6.35056 | 0.001424 | -0.65049 | 2.74343E-06 |
| | Bo9g088590.1 | 0 | -7.58105 | -3.63027 | 0.001455 | -1.19911 | 2.84863E-06 |

| | | | | | | | |
|---|---|---|---|---|---|---|---|
| | Bo8g102270.1 | 0 | -1.27485 | -1.43555 | 0.002108 | 4.937314 | 4.19005E-06 |
| ELIP1 | Bo3g081430.1 | 0 | -2.41752 | -1.99291 | 0.002164 | 5.294742 | 4.36845E-06 |
| HSP17 | Bo3g184440.1 | 0 | -0.75461 | 1.436827 | 0.002781 | 5.200598 | 5.69881E-06 |
| HGL1 | Bo1g101520.1 | 0 | 1.839715 | 0.986736 | 0.002999 | 3.888444 | 6.23749E-06 |
| GH9B14 | Bo1g036810.1 | 0 | -5.18657 | -7.82584 | 0.003128 | -1.06443 | 6.60063E-06 |
| | Bo4g197170.1 | 0 | -1.21549 | -3.20772 | 0.00394 | 2.587653 | 8.435E-06 |
| | Bo3g004420.1 | 0 | -9.45917 | 0.965808 | 0.004463 | 1.856675 | 9.69113E-06 |
| PER22 | Bo3g032440.1 | 0 | 2.273678 | 1.119362 | 0.005238 | 3.875137 | 1.15322E-05 |
| UPS1 | Bo9g074290.1 | 0 | -3.74205 | -3.25402 | 0.006074 | 1.29751 | 1.35594E-05 |
| | Bo4g195770.1 | 0 | 2.36194 | 1.136038 | 0.006143 | 6.142988 | 1.39027E-05 |
| DJ1F | Bo6g068170.1 | 0 | -1.51628 | -1.19737 | 0.006524 | 6.695567 | 1.49639E-05 |
| | Bo5g115730.1 | 0 | -2.26307 | -0.50319 | 0.006778 | 2.344661 | 1.57525E-05 |
| | Bo1g021490.1 | 0 | -1.54418 | 2.030074 | 0.008832 | 4.759035 | 2.10304E-05 |
| APK2 | Bo7g118850.1 | 0 | -1.56277 | -2.44627 | 0.008832 | 3.067667 | 2.1066E-05 |
| Bidirectional sugar transporter SWEET | Bo2g011110.1 | 0 | 1.456398 | 1.013074 | 0.010613 | 4.600799 | 2.59121E-05 |
| | Bo4g024930.1 | 0 | 1.952708 | 1.026226 | 0.010613 | 5.346896 | 2.59644E-05 |
| | Bo20568s010.1 | 0 | -1.92166 | -3.12335 | 0.010649 | 0.576004 | 2.63777E-05 |
| | Bo2g131800.1 | 0 | -2.42588 | -2.14251 | 0.010888 | 0.448139 | 2.73036E-05 |

| | | | | | | | |
|---|---|---|---|---|---|---|---|
| ATL6 | Bo3g058810.1 | 0 | -2.23772 | -2.41175 | 0.011592 | 1.130277 | 2.97775E-05 |
| GSTU21 | Bo6g121970.1 | 0 | 2.309282 | 1.10415 | 0.011592 | 4.86646 | 2.95324E-05 |
| | Bo3g071810.1 | 0 | 2.148125 | -0.44848 | 0.013344 | -0.23401 | 3.46866E-05 |
| | Bo4g125510.1 | 0 | -2.08751 | -1.21523 | 0.013642 | 1.609771 | 3.58791E-05 |
| HSP17 | Bo9g169680.1 | 0 | -1.53582 | 1.427684 | 0.013832 | 1.513107 | 3.72248E-05 |
| | Bo3g005520.1 | 0 | 3.429366 | 3.5713 | 0.013832 | 0.255519 | 3.68107E-05 |
| | Bo8g051290.1 | 0 | 0.616228 | -6.89555 | 0.014121 | -0.74987 | 3.8432E-05 |
| MED37C | Bo1g139130.1 | 0 | -0.09391 | 1.312288 | 0.014411 | 4.970245 | 3.96639E-05 |
| | Bo5g003820.1 | 0 | 2.51891 | 1.299547 | 0.014516 | 3.747528 | 4.03966E-05 |
| CLPB1 | Bo6g087090.1 | 0 | -0.58393 | 0.929233 | 0.016213 | 4.502605 | 4.5614E-05 |
| | Bo01487s010.1 | 0 | -3.10759 | -2.33199 | 0.016314 | 5.156308 | 4.63962E-05 |
| RVE7 | Bo5g025310.1 | 0 | -1.01324 | 1.273655 | 0.016687 | 4.793453 | 4.84805E-05 |
| AHL16 | Bo4g018450.1 | 0 | 2.68252 | 1.460247 | 0.016687 | 5.381853 | 4.80963E-05 |
| CYP702A5 | Bo7g103690.1 | 0 | -1.88181 | -7.89148 | 0.016755 | -0.73812 | 4.97019E-05 |
| | Bo5g116750.1 | 0 | -2.60731 | -0.07091 | 0.016755 | 1.973689 | 4.94042E-05 |
| | Bo3g057050.1 | 0 | 0.735378 | 4.829231 | 0.017273 | 0.171832 | 5.22938E-05 |
| ADPG2 | Bo4g191100.1 | 0 | -7.43376 | -4.81919 | 0.017273 | -1.36754 | 5.21111E-05 |
| CHC2 | Bo5g139850.1 | 0 | 2.763624 | 1.408513 | 0.018573 | 1.899302 | 5.67994E-05 |
| | Bo9g123440.1 | 0 | 2.013686 | 1.038784 | 0.019285 | 3.903425 | 5.95667E-05 |

| | | | | | | | |
|---|---|---|---|---|---|---|---|
| xyloglucan endotransglucosylase/hydrolase 3 | Bo7g031510.1 | 0 | -7.65519 | 0.224891 | 0.021434 | 6.781521 | 6.68579E-05 |
| ABCG35 | Bo5g020620.1 | 0 | 2.583905 | 1.379383 | 0.022704 | 2.569325 | 7.15148E-05 |
| SAG12 | Bo2g127490.1 | 0 | -10.0976 | -0.96616 | 0.025838 | 1.542984 | 8.29669E-05 |
| | Bo12355s010.1 | 0 | 4.739088 | 6.905502 | 0.025838 | -1.58175 | 8.29618E-05 |
| | Bo9g076150.1 | 0 | -1.16407 | 1.030579 | 0.026597 | 3.125014 | 8.70305E-05 |
| | Bo8g021770.1 | 0 | 1.94893 | 0.630613 | 0.026597 | 5.870878 | 8.6251E-05 |
| DTX56 | Bo1g030730.1 | 0 | 1.244496 | 0.212276 | 0.027316 | 4.362198 | 9.02175E-05 |
| ACT4 | Bo3g018230.1 | 0 | 3.043831 | -0.75992 | 0.030835 | 1.905822 | 0.000102782 |
| Phosphoenolpyruvate carboxylase family protein | Bo5g034350.1 | 0 | -1.15206 | -0.97645 | 0.031039 | 4.662463 | 0.000104411 |
| ACA7 | Bo5g008760.1 | 0 | -2.68197 | -2.32395 | 0.03287 | 0.878668 | 0.000111576 |
| | Bo9g063050.1 | 0 | 1.802637 | 1.124232 | 0.0334 | 4.424348 | 0.000114397 |
| | Bo1g037570.1 | 0 | 1.394983 | 0.491934 | 0.033713 | 4.369327 | 0.0001165 |
| GH3.17 | Bo5g053450.1 | 0 | -0.33704 | -2.65354 | 0.034072 | 0.401131 | 0.000118782 |
| | Bo5g003500.1 | 0 | 1.786021 | 0.808235 | 0.034337 | 3.315835 | 0.000120755 |
| RING-type E3 ubiquitin transferase | Bo3g029820.1 | 0 | 0.875459 | -1.30715 | 0.03475 | 1.317603 | 0.000123271 |
| | Bo5g123500.1 | 0 | -1.3476 | -0.55981 | 0.035699 | 4.052225 | 0.000127729 |
| PIP2B | Bo3g031550.1 | 0 | -2.04539 | -1.59045 | 0.035888 | 3.439107 | 0.000129505 |
| 3-ketoacyl-CoA synthase 18 | Bo3g168810.1 | 0 | -7.14399 | -7.14399 | 0.037873 | -1.64334 | 0.000137825 |
| Beta-galactosidase | Bo1g042930.1 | 0 | 1.287432 | 0.73435 | 0.038314 | 3.735557 | 0.000140604 |

| | | | | | | | |
|---|---|---|---|---|---|---|---|
| | Bo3g102950.1 | 0 | -4.52312 | -5.40958 | 0.039535 | -1.49295 | 0.000146291 |
| JUB1 | Bo3g036290.1 | 0 | 2.822932 | 1.385796 | 0.039944 | 3.701262 | 0.000149025 |
| | Bo5g127130.1 | 0 | -4.94586 | 1.345348 | 0.040349 | 7.011637 | 0.00015177 |
| | Bo6g009310.1 | 0 | -10.3807 | -1.61144 | 0.041186 | 1.650543 | 0.000157439 |
| | Bo4g106440.1 | 0 | -14.8581 | -0.21583 | 0.041186 | 6.568686 | 0.000157088 |
| | Bo01904s010.1 | 0 | 3.488754 | 3.024441 | 0.041782 | -0.21386 | 0.000162017 |
| | Bo6g089280.1 | 0 | -6.17092 | -6.17092 | 0.041782 | -2.28354 | 0.000164558 |
| | Bo07665s010.1 | 0 | 3.735232 | 1.353204 | 0.041782 | -0.55648 | 0.000164599 |
| | Bo7g032910.1 | 0 | -4.96637 | -3.75755 | 0.041782 | -1.72945 | 0.000164829 |
| | Bo9g140800.1 | 0 | -1.55741 | -2.61572 | 0.044459 | 0.029428 | 0.000176749 |
| SSL13 | Bo01353s050.1 | 0 | 2.341752 | 1.358546 | 0.045047 | 6.294444 | 0.000181842 |
| | Bo9g011230.1 | 0 | 4.130319 | 1.412663 | 0.045047 | 0.221593 | 0.000181192 |
| | Bo2g008670.1 | 0 | -12.8276 | 1.108907 | 0.045316 | 5.301459 | 0.000184311 |

**Supplementary Table 7**

Gene Ontology (GO) Enrichment Analysis of significantly differentially expressed genes ($p<0.05$) in the vegetative leaf tissue of both mutant lines (above) using TopGo (Alexa and Rahnenführer, 2009). The gene set analysis statistically compared the representation of GO (Gene Ontology) terms, which were first assigned to the *Brassica oleracea* gene models using the *Arabidopsis thaliana* org.At.tair.db library (version 3.2.3) (Carlson, 2016), in the gene set above to that of the 'expected' value, to determine those to be considered 'over-represented' using the BP parameter with a node size of 10, then confirming the statistical significance of this using Fisher's exact test. This was performed in RStudio v.1.2.5042 (Racine, 2012).

| | GO.ID | Term | Annotated | Significant | Expected | classicFisher |
|---|---|---|---|---|---|---|
| 1 | GO:0016144 | S-glycoside biosynthetic process | 300 | 32 | 0.71 | < 1e-30 |
| 2 | GO:0019758 | glycosinolate biosynthetic process | 300 | 32 | 0.71 | < 1e-30 |
| 3 | GO:0019761 | glucosinolate biosynthetic process | 300 | 32 | 0.71 | < 1e-30 |
| 4 | GO:1901659 | glycosyl compound biosynthetic process | 311 | 32 | 0.73 | < 1e-30 |
| 5 | GO:0016143 | S-glycoside metabolic process | 368 | 32 | 0.87 | < 1e-30 |
| 6 | GO:0019757 | glycosinolate metabolic process | 368 | 32 | 0.87 | < 1e-30 |
| 7 | GO:0019760 | glucosinolate metabolic process | 368 | 32 | 0.87 | < 1e-30 |
| 8 | GO:1901657 | glycosyl compound metabolic process | 412 | 32 | 0.97 | < 1e-30 |
| 9 | GO:0044550 | secondary metabolite biosynthetic proces... | 546 | 33 | 1.29 | < 1e-30 |
| 10 | GO:0009684 | indoleacetic acid biosynthetic process | 193 | 25 | 0.46 | < 1e-30 |
| 11 | GO:0009683 | indoleacetic acid metabolic process | 195 | 25 | 0.46 | < 1e-30 |
| 12 | GO:0044272 | sulfur compound biosynthetic process | 845 | 36 | 2 | < 1e-30 |
| 13 | GO:0009851 | auxin biosynthetic process | 223 | 25 | 0.53 | < 1e-30 |
| 14 | GO:0042435 | indole-containing compound biosynthetic ... | 236 | 25 | 0.56 | < 1e-30 |
| 15 | GO:0009850 | auxin metabolic process | 253 | 25 | 0.6 | < 1e-30 |
| 16 | GO:0019748 | secondary metabolic process | 1119 | 37 | 2.64 | < 1e-30 |
| 17 | GO:0042430 | indole-containing compound metabolic pro... | 273 | 25 | 0.65 | < 1e-30 |
| 18 | GO:0006790 | sulfur compound metabolic process | 1232 | 37 | 2.91 | < 1e-30 |
| 19 | GO:0034754 | cellular hormone metabolic process | 307 | 25 | 0.73 | < 1e-30 |

| | | | | | | |
|---|---|---|---|---|---|---|
| 20 | GO:1901605 | alpha-amino acid metabolic process | 936 | 30 | 2.21 | 1.00E-26 |
| 21 | GO:0006520 | cellular amino acid metabolic process | 1185 | 30 | 2.8 | 9.60E-24 |
| 22 | GO:1901137 | carbohydrate derivative biosynthetic pro... | 1538 | 32 | 3.63 | 7.20E-23 |
| 23 | GO:0043436 | oxoacid metabolic process | 3444 | 41 | 8.14 | 5.00E-21 |
| 24 | GO:0006082 | organic acid metabolic process | 3445 | 41 | 8.14 | 5.00E-21 |
| 25 | GO:1901607 | alpha-amino acid biosynthetic process | 672 | 23 | 1.59 | 6.90E-21 |
| 26 | GO:1901566 | organonitrogen compound biosynthetic pro... | 3504 | 41 | 8.28 | 9.50E-21 |
| 27 | GO:0019344 | cysteine biosynthetic process | 375 | 19 | 0.89 | 2.00E-20 |
| 28 | GO:0006534 | cysteine metabolic process | 377 | 19 | 0.89 | 2.20E-20 |
| 29 | GO:0009070 | serine family amino acid biosynthetic pr... | 381 | 19 | 0.9 | 2.70E-20 |
| 30 | GO:0016053 | organic acid biosynthetic process | 2157 | 33 | 5.1 | 1.60E-19 |
| 31 | GO:0046394 | carboxylic acid biosynthetic process | 2157 | 33 | 5.1 | 1.60E-19 |
| 32 | GO:0008652 | cellular amino acid biosynthetic process | 788 | 23 | 1.86 | 2.40E-19 |
| 33 | GO:0000097 | sulfur amino acid biosynthetic process | 525 | 20 | 1.24 | 4.60E-19 |
| 34 | GO:0009069 | serine family amino acid metabolic proce... | 463 | 19 | 1.09 | 1.00E-18 |
| 35 | GO:0000096 | sulfur amino acid metabolic process | 598 | 20 | 1.41 | 5.80E-18 |
| 36 | GO:1901135 | carbohydrate derivative metabolic proces... | 2247 | 32 | 5.31 | 5.80E-18 |
| 37 | GO:0044283 | small molecule biosynthetic process | 2481 | 33 | 5.86 | 1.10E-17 |
| 38 | GO:0072330 | monocarboxylic acid biosynthetic process | 1471 | 27 | 3.48 | 1.10E-17 |
| 39 | GO:0044281 | small molecule metabolic process | 4793 | 43 | 11.32 | 1.50E-17 |
| 40 | GO:0042446 | hormone biosynthetic process | 1385 | 25 | 3.27 | 3.80E-16 |
| 41 | GO:0019752 | carboxylic acid metabolic process | 3262 | 34 | 7.71 | 4.40E-15 |
| 42 | GO:0042445 | hormone metabolic process | 1548 | 25 | 3.66 | 4.90E-15 |
| 43 | GO:0006569 | tryptophan catabolic process | 130 | 11 | 0.31 | 1.30E-14 |
| 44 | GO:0046218 | indolalkylamine catabolic process | 130 | 11 | 0.31 | 1.30E-14 |

| | | | | | | |
|---|---|---|---|---|---|---|
| 45 | GO:0042436 | indole-containing compound catabolic pro... | 135 | 11 | 0.32 | 2.00E-14 |
| 46 | GO:0009074 | aromatic amino acid family catabolic pro... | 139 | 11 | 0.33 | 2.80E-14 |
| 47 | GO:0010817 | regulation of hormone levels | 1791 | 25 | 4.23 | 1.30E-13 |
| 48 | GO:0006568 | tryptophan metabolic process | 161 | 11 | 0.38 | 1.40E-13 |
| 49 | GO:0006586 | indolalkylamine metabolic process | 161 | 11 | 0.38 | 1.40E-13 |
| 50 | GO:0032787 | monocarboxylic acid metabolic process | 2388 | 28 | 5.64 | 2.00E-13 |
| 51 | GO:0009098 | leucine biosynthetic process | 13 | 6 | 0.03 | 2.40E-13 |
| 52 | GO:0009310 | amine catabolic process | 209 | 11 | 0.49 | 2.50E-12 |
| 53 | GO:0042402 | cellular biogenic amine catabolic proces... | 209 | 11 | 0.49 | 2.50E-12 |
| 54 | GO:1901564 | organonitrogen compound metabolic proces... | 7265 | 44 | 17.17 | 1.20E-11 |
| 55 | GO:0006551 | leucine metabolic process | 23 | 6 | 0.05 | 1.40E-11 |
| 56 | GO:0009082 | branched-chain amino acid biosynthetic p... | 27 | 6 | 0.06 | 4.00E-11 |
| 57 | GO:1901606 | alpha-amino acid catabolic process | 275 | 11 | 0.65 | 4.80E-11 |
| 58 | GO:0009063 | cellular amino acid catabolic process | 279 | 11 | 0.66 | 5.60E-11 |
| 59 | GO:0006576 | cellular biogenic amine metabolic proces... | 281 | 11 | 0.66 | 6.10E-11 |
| 60 | GO:0044106 | cellular amine metabolic process | 296 | 11 | 0.7 | 1.10E-10 |
| 61 | GO:0010439 | regulation of glucosinolate biosynthetic... | 16 | 5 | 0.04 | 2.70E-10 |
| 62 | GO:0009081 | branched-chain amino acid metabolic proc... | 37 | 6 | 0.09 | 3.10E-10 |
| 63 | GO:0065008 | regulation of biological quality | 2914 | 26 | 6.88 | 8.70E-10 |
| 64 | GO:0009072 | aromatic amino acid family metabolic pro... | 406 | 11 | 0.96 | 2.90E-09 |
| 65 | GO:0009308 | amine metabolic process | 408 | 11 | 0.96 | 3.10E-09 |
| 66 | GO:1900376 | regulation of secondary metabolite biosy... | 28 | 5 | 0.07 | 6.00E-09 |
| 67 | GO:0044249 | cellular biosynthetic process | 9247 | 45 | 21.85 | 1.20E-08 |
| 68 | GO:0042762 | regulation of sulfur metabolic process | 36 | 5 | 0.09 | 2.30E-08 |
| 69 | GO:1901576 | organic substance biosynthetic process | 9857 | 46 | 23.29 | 2.80E-08 |

| | | | | | | |
|---|---|---|---|---|---|---|
| 70 | GO:0010675 | regulation of cellular carbohydrate meta... | 41 | 5 | 0.1 | 4.50E-08 |
| 71 | GO:0009058 | biosynthetic process | 10049 | 46 | 23.74 | 5.50E-08 |
| 72 | GO:0043255 | regulation of carbohydrate biosynthetic ... | 44 | 5 | 0.1 | 6.40E-08 |
| 73 | GO:0062012 | regulation of small molecule metabolic p... | 95 | 6 | 0.22 | 1.00E-07 |
| 74 | GO:0044248 | cellular catabolic process | 2666 | 22 | 6.3 | 1.10E-07 |
| 75 | GO:0043455 | regulation of secondary metabolic proces... | 50 | 5 | 0.12 | 1.20E-07 |
| 76 | GO:0044270 | cellular nitrogen compound catabolic pro... | 890 | 13 | 2.1 | 1.50E-07 |
| 77 | GO:0019439 | aromatic compound catabolic process | 893 | 13 | 2.11 | 1.50E-07 |
| 78 | GO:0046700 | heterocycle catabolic process | 897 | 13 | 2.12 | 1.60E-07 |
| 79 | GO:1901361 | organic cyclic compound catabolic proces... | 911 | 13 | 2.15 | 1.90E-07 |
| 80 | GO:0009056 | catabolic process | 3010 | 23 | 7.11 | 2.10E-07 |
| 81 | GO:0016054 | organic acid catabolic process | 677 | 11 | 1.6 | 5.20E-07 |
| 82 | GO:0046395 | carboxylic acid catabolic process | 677 | 11 | 1.6 | 5.20E-07 |
| 83 | GO:0006109 | regulation of carbohydrate metabolic pro... | 70 | 5 | 0.17 | 6.80E-07 |
| 84 | GO:0044237 | cellular metabolic process | 13631 | 52 | 32.21 | 1.40E-06 |
| 85 | GO:0008152 | metabolic process | 15048 | 55 | 35.55 | 1.50E-06 |
| 86 | GO:0009407 | toxin catabolic process | 359 | 8 | 0.85 | 2.10E-06 |
| 87 | GO:0009404 | toxin metabolic process | 386 | 8 | 0.91 | 3.60E-06 |
| 88 | GO:0018130 | heterocycle biosynthetic process | 5004 | 28 | 11.82 | 3.80E-06 |
| 89 | GO:0098754 | detoxification | 389 | 8 | 0.92 | 3.80E-06 |
| 90 | GO:0009987 | cellular process | 17469 | 59 | 41.27 | 4.60E-06 |
| 91 | GO:0006807 | nitrogen compound metabolic process | 11102 | 45 | 26.23 | 4.80E-06 |
| 92 | GO:0019438 | aromatic compound biosynthetic process | 5386 | 29 | 12.73 | 5.00E-06 |
| 93 | GO:0044262 | cellular carbohydrate metabolic process | 1066 | 12 | 2.52 | 6.90E-06 |
| 94 | GO:0044282 | small molecule catabolic process | 1092 | 12 | 2.58 | 8.80E-06 |

| | | | | | | |
|---|---|---|---|---|---|---|
| 95 | GO:0009617 | response to bacterium | 1124 | 12 | 2.66 | 1.20E-05 |
| 96 | GO:0000103 | sulfate assimilation | 20 | 3 | 0.05 | 1.40E-05 |
| 97 | GO:1901362 | organic cyclic compound biosynthetic pro... | 5771 | 29 | 13.64 | 2.00E-05 |
| 98 | GO:0042742 | defense response to bacterium | 826 | 10 | 1.95 | 2.40E-05 |
| 99 | GO:0016051 | carbohydrate biosynthetic process | 1417 | 13 | 3.35 | 2.40E-05 |
| 100 | GO:0044271 | cellular nitrogen compound biosynthetic ... | 5604 | 28 | 13.24 | 3.40E-05 |
| 101 | GO:1901565 | organonitrogen compound catabolic proces... | 1310 | 12 | 3.1 | 5.20E-05 |
| 102 | GO:0019252 | starch biosynthetic process | 280 | 6 | 0.66 | 5.30E-05 |
| 103 | GO:0071704 | organic substance metabolic process | 14207 | 50 | 33.57 | 5.70E-05 |
| 104 | GO:0009636 | response to toxic substance | 748 | 9 | 1.77 | 6.50E-05 |
| 105 | GO:0009607 | response to biotic stimulus | 2610 | 17 | 6.17 | 9.30E-05 |
| 106 | GO:0043207 | response to external biotic stimulus | 2610 | 17 | 6.17 | 9.30E-05 |
| 107 | GO:0051707 | response to other organism | 2610 | 17 | 6.17 | 9.30E-05 |
| 108 | GO:0009611 | response to wounding | 619 | 8 | 1.46 | 0.0001 |
| 109 | GO:0050896 | response to stimulus | 10855 | 41 | 25.65 | 0.00015 |
| 110 | GO:0005975 | carbohydrate metabolic process | 2247 | 15 | 5.31 | 0.0002 |
| 111 | GO:0005982 | starch metabolic process | 361 | 6 | 0.85 | 0.00021 |
| 112 | GO:0009250 | glucan biosynthetic process | 433 | 6 | 1.02 | 0.00056 |
| 113 | GO:0006950 | response to stress | 6643 | 28 | 15.7 | 0.00071 |
| 114 | GO:1901575 | organic substance catabolic process | 2543 | 15 | 6.01 | 0.00075 |
| 115 | GO:0046483 | heterocycle metabolic process | 7468 | 30 | 17.64 | 0.00094 |
| 116 | GO:1901360 | organic cyclic compound metabolic proces... | 8211 | 32 | 19.4 | 0.00097 |
| 117 | GO:0009067 | aspartate family amino acid biosynthetic... | 196 | 4 | 0.46 | 0.00121 |
| 118 | GO:0048232 | male gamete generation | 90 | 3 | 0.21 | 0.00128 |
| 119 | GO:0006725 | cellular aromatic compound metabolic pro... | 7976 | 31 | 18.84 | 0.0013 |

| | | | | | | |
|---|---|---|---|---|---|---|
| 120 | GO:0033692 | cellular polysaccharide biosynthetic pro... | 726 | 7 | 1.72 | 0.00162 |
| 121 | GO:0051704 | multi-organism process | 3662 | 18 | 8.65 | 0.00173 |
| 122 | GO:0034641 | cellular nitrogen compound metabolic pro... | 7840 | 30 | 18.52 | 0.00216 |
| 123 | GO:0009605 | response to external stimulus | 3770 | 18 | 8.91 | 0.0024 |
| 124 | GO:0009066 | aspartate family amino acid metabolic pr... | 238 | 4 | 0.56 | 0.00246 |
| 125 | GO:0034637 | cellular carbohydrate biosynthetic proce... | 783 | 7 | 1.85 | 0.00248 |
| 126 | GO:0006073 | cellular glucan metabolic process | 612 | 6 | 1.45 | 0.00325 |
| 127 | GO:0044042 | glucan metabolic process | 612 | 6 | 1.45 | 0.00325 |
| 128 | GO:0006566 | threonine metabolic process | 36 | 2 | 0.09 | 0.00329 |
| 129 | GO:0044238 | primary metabolic process | 12883 | 42 | 30.44 | 0.00394 |
| 130 | GO:0006098 | pentose-phosphate shunt | 290 | 4 | 0.69 | 0.00497 |
| 131 | GO:0051156 | glucose 6-phosphate metabolic process | 290 | 4 | 0.69 | 0.00497 |
| 132 | GO:0000271 | polysaccharide biosynthetic process | 892 | 7 | 2.11 | 0.00507 |
| 133 | GO:0044264 | cellular polysaccharide metabolic proces... | 900 | 7 | 2.13 | 0.00532 |
| 134 | GO:0046496 | nicotinamide nucleotide metabolic proces... | 681 | 6 | 1.61 | 0.00545 |
| 135 | GO:0098542 | defense response to other organism | 1935 | 11 | 4.57 | 0.00548 |
| 136 | GO:0019362 | pyridine nucleotide metabolic process | 686 | 6 | 1.62 | 0.00564 |
| 137 | GO:0006739 | NADP metabolic process | 302 | 4 | 0.71 | 0.00573 |
| 138 | GO:0072524 | pyridine-containing compound metabolic p... | 693 | 6 | 1.64 | 0.00592 |
| 139 | GO:0006090 | pyruvate metabolic process | 767 | 6 | 1.81 | 0.00952 |
| 140 | GO:0065007 | biological regulation | 9405 | 32 | 22.22 | 0.0099 |

**Supplementary Table 8**

Gene Ontology (GO) Enrichment Analysis of significantly differentially expressed genes ($p<0.05$) in the generative leaf tissue of both mutant lines (above) using TopGo (Alexa and Rahnenführer, 2009). The gene set analysis statistically compared the representation of GO (Gene Ontology) terms ,which were first assigned to the *Brassica oleracea* gene models using the *Arabidopsis thaliana* org.At.tair.db library (version 3.2.3) (Carlson, 2016), in the gene set above to that of the 'expected' value, to determine those to be considered 'over-represented' using the BP parameter with a node size of 10, then confirming the statistical significance of this using Fisher's exact test. This was performed in RStudio v.1.2.5042 (Racine, 2012).

| | GO.ID | Term | Annotated | Significant | Expected | classicFisher |
|---|---|---|---|---|---|---|
| 1 | GO:0016143 | S-glycoside metabolic process | 368 | 30 | 1.53 | 2.80E-30 |
| 2 | GO:0019757 | glycosinolate metabolic process | 368 | 30 | 1.53 | 2.80E-30 |
| 3 | GO:0019760 | glucosinolate metabolic process | 368 | 30 | 1.53 | 2.80E-30 |
| 4 | GO:0016144 | S-glycoside biosynthetic process | 300 | 28 | 1.25 | 6.50E-30 |
| 5 | GO:0019758 | glycosinolate biosynthetic process | 300 | 28 | 1.25 | 6.50E-30 |
| 6 | GO:0019761 | glucosinolate biosynthetic process | 300 | 28 | 1.25 | 6.50E-30 |
| 7 | GO:1901659 | glycosyl compound biosynthetic process | 311 | 28 | 1.29 | 1.80E-29 |
| 8 | GO:1901657 | glycosyl compound metabolic process | 412 | 30 | 1.71 | 8.40E-29 |
| 9 | GO:0019748 | secondary metabolic process | 1119 | 39 | 4.65 | 1.40E-25 |
| 10 | GO:0044550 | secondary metabolite biosynthetic proces... | 546 | 28 | 2.27 | 1.00E-22 |
| 11 | GO:0009684 | indoleacetic acid biosynthetic process | 193 | 19 | 0.8 | 6.30E-21 |
| 12 | GO:0009683 | indoleacetic acid metabolic process | 195 | 19 | 0.81 | 7.70E-21 |
| 13 | GO:0009850 | auxin metabolic process | 253 | 20 | 1.05 | 4.60E-20 |
| 14 | GO:0009851 | auxin biosynthetic process | 223 | 19 | 0.93 | 1.00E-19 |
| 15 | GO:0042430 | indole-containing compound metabolic pro... | 273 | 20 | 1.13 | 2.10E-19 |
| 16 | GO:0042435 | indole-containing compound biosynthetic ... | 236 | 19 | 0.98 | 3.00E-19 |
| 17 | GO:0044272 | sulfur compound biosynthetic process | 845 | 28 | 3.51 | 1.10E-17 |
| 18 | GO:0034754 | cellular hormone metabolic process | 307 | 19 | 1.27 | 4.10E-17 |
| 19 | GO:0006790 | sulfur compound metabolic process | 1232 | 31 | 5.11 | 3.20E-16 |

| | | | | | | |
|---|---|---|---|---|---|---|
| 20 | GO:0043455 | regulation of secondary metabolic proces... | 50 | 9 | 0.21 | 5.90E-13 |
| 21 | GO:1901137 | carbohydrate derivative biosynthetic pro... | 1538 | 29 | 6.39 | 4.70E-12 |
| 22 | GO:1901605 | alpha-amino acid metabolic process | 936 | 23 | 3.89 | 6.00E-12 |
| 23 | GO:0043436 | oxoacid metabolic process | 3444 | 43 | 14.3 | 1.10E-11 |
| 24 | GO:0006082 | organic acid metabolic process | 3445 | 43 | 14.3 | 1.10E-11 |
| 25 | GO:0006520 | cellular amino acid metabolic process | 1185 | 25 | 4.92 | 1.70E-11 |
| 26 | GO:0010439 | regulation of glucosinolate biosynthetic... | 16 | 6 | 0.07 | 3.50E-11 |
| 27 | GO:0042445 | hormone metabolic process | 1548 | 27 | 6.43 | 1.70E-10 |
| 28 | GO:0006569 | tryptophan catabolic process | 130 | 10 | 0.54 | 1.80E-10 |
| 29 | GO:0046218 | indolalkylamine catabolic process | 130 | 10 | 0.54 | 1.80E-10 |
| 30 | GO:0062012 | regulation of small molecule metabolic p... | 95 | 9 | 0.39 | 2.40E-10 |
| 31 | GO:0042436 | indole-containing compound catabolic pro... | 135 | 10 | 0.56 | 2.70E-10 |
| 32 | GO:0009074 | aromatic amino acid family catabolic pro... | 139 | 10 | 0.58 | 3.60E-10 |
| 33 | GO:0006950 | response to stress | 6643 | 59 | 27.58 | 3.90E-10 |
| 34 | GO:0042446 | hormone biosynthetic process | 1385 | 25 | 5.75 | 4.50E-10 |
| 35 | GO:0006568 | tryptophan metabolic process | 161 | 10 | 0.67 | 1.50E-09 |
| 36 | GO:0006586 | indolalkylamine metabolic process | 161 | 10 | 0.67 | 1.50E-09 |
| 37 | GO:1900376 | regulation of secondary metabolite biosy... | 28 | 6 | 0.12 | 1.60E-09 |
| 38 | GO:1901135 | carbohydrate derivative metabolic proces... | 2247 | 31 | 9.33 | 1.90E-09 |
| 39 | GO:0009636 | response to toxic substance | 748 | 18 | 3.11 | 2.10E-09 |
| 40 | GO:0010817 | regulation of hormone levels | 1791 | 27 | 7.44 | 4.10E-09 |
| 41 | GO:0042762 | regulation of sulfur metabolic process | 36 | 6 | 0.15 | 8.00E-09 |
| 42 | GO:0016053 | organic acid biosynthetic process | 2157 | 29 | 8.96 | 1.20E-08 |
| 43 | GO:0046394 | carboxylic acid biosynthetic process | 2157 | 29 | 8.96 | 1.20E-08 |
| 44 | GO:0010675 | regulation of cellular carbohydrate meta... | 41 | 6 | 0.17 | 1.80E-08 |

| | | | | | | |
|---|---|---|---|---|---|---|
| 45 | GO:0009310 | amine catabolic process | 209 | 10 | 0.87 | 1.80E-08 |
| 46 | GO:0042402 | cellular biogenic amine catabolic proces... | 209 | 10 | 0.87 | 1.80E-08 |
| 47 | GO:1901607 | alpha-amino acid biosynthetic process | 672 | 16 | 2.79 | 2.10E-08 |
| 48 | GO:1901606 | alpha-amino acid catabolic process | 275 | 11 | 1.14 | 2.20E-08 |
| 49 | GO:0009063 | cellular amino acid catabolic process | 279 | 11 | 1.16 | 2.60E-08 |
| 50 | GO:0043255 | regulation of carbohydrate biosynthetic ... | 44 | 6 | 0.18 | 2.80E-08 |
| 51 | GO:0050896 | response to stimulus | 10855 | 75 | 45.07 | 3.50E-08 |
| 52 | GO:0019344 | cysteine biosynthetic process | 375 | 12 | 1.56 | 5.80E-08 |
| 53 | GO:0006534 | cysteine metabolic process | 377 | 12 | 1.57 | 6.10E-08 |
| 54 | GO:0065008 | regulation of biological quality | 2914 | 33 | 12.1 | 6.30E-08 |
| 55 | GO:0009070 | serine family amino acid biosynthetic pr... | 381 | 12 | 1.58 | 6.90E-08 |
| 56 | GO:0009069 | serine family amino acid metabolic proce... | 463 | 13 | 1.92 | 7.20E-08 |
| 57 | GO:0098754 | detoxification | 389 | 12 | 1.61 | 8.60E-08 |
| 58 | GO:0044281 | small molecule metabolic process | 4793 | 44 | 19.9 | 9.80E-08 |
| 59 | GO:0009072 | aromatic amino acid family metabolic pro... | 406 | 12 | 1.69 | 1.40E-07 |
| 60 | GO:0008652 | cellular amino acid biosynthetic process | 788 | 16 | 3.27 | 1.80E-07 |
| 61 | GO:0009098 | leucine biosynthetic process | 13 | 4 | 0.05 | 2.00E-07 |
| 62 | GO:0044283 | small molecule biosynthetic process | 2481 | 29 | 10.3 | 2.60E-07 |
| 63 | GO:0019752 | carboxylic acid metabolic process | 3262 | 34 | 13.54 | 2.70E-07 |
| 64 | GO:0006576 | cellular biogenic amine metabolic proces... | 281 | 10 | 1.17 | 2.90E-07 |
| 65 | GO:0009407 | toxin catabolic process | 359 | 11 | 1.49 | 3.30E-07 |
| 66 | GO:0044106 | cellular amine metabolic process | 296 | 10 | 1.23 | 4.70E-07 |
| 67 | GO:0006109 | regulation of carbohydrate metabolic pro... | 70 | 6 | 0.29 | 4.80E-07 |
| 68 | GO:1901566 | organonitrogen compound biosynthetic pro... | 3504 | 35 | 14.55 | 4.80E-07 |
| 69 | GO:0001101 | response to acid chemical | 2896 | 31 | 12.02 | 6.10E-07 |

| | | | | | | |
|---|---|---|---|---|---|---|
| 70 | GO:0009404 | toxin metabolic process | 386 | 11 | 1.6 | 6.70E-07 |
| 71 | GO:0042221 | response to chemical | 6670 | 52 | 27.69 | 7.80E-07 |
| 72 | GO:1901700 | response to oxygen-containing compound | 4324 | 39 | 17.95 | 1.20E-06 |
| 73 | GO:0000103 | sulfate assimilation | 20 | 4 | 0.08 | 1.30E-06 |
| 74 | GO:0000097 | sulfur amino acid biosynthetic process | 525 | 12 | 2.18 | 2.00E-06 |
| 75 | GO:0009987 | cellular process | 17469 | 97 | 72.52 | 2.30E-06 |
| 76 | GO:0006551 | leucine metabolic process | 23 | 4 | 0.1 | 2.40E-06 |
| 77 | GO:0015770 | sucrose transport | 23 | 4 | 0.1 | 2.40E-06 |
| 78 | GO:0072330 | monocarboxylic acid biosynthetic process | 1471 | 20 | 6.11 | 2.80E-06 |
| 79 | GO:0015766 | disaccharide transport | 25 | 4 | 0.1 | 3.30E-06 |
| 80 | GO:0015772 | oligosaccharide transport | 25 | 4 | 0.1 | 3.30E-06 |
| 81 | GO:0009082 | branched-chain amino acid biosynthetic p... | 27 | 4 | 0.11 | 4.60E-06 |
| 82 | GO:0016054 | organic acid catabolic process | 677 | 13 | 2.81 | 5.10E-06 |
| 83 | GO:0046395 | carboxylic acid catabolic process | 677 | 13 | 2.81 | 5.10E-06 |
| 84 | GO:0000096 | sulfur amino acid metabolic process | 598 | 12 | 2.48 | 7.70E-06 |
| 85 | GO:0009308 | amine metabolic process | 408 | 10 | 1.69 | 8.30E-06 |
| 86 | GO:0042742 | defense response to bacterium | 826 | 14 | 3.43 | 8.90E-06 |
| 87 | GO:0032787 | monocarboxylic acid metabolic process | 2388 | 25 | 9.91 | 1.40E-05 |
| 88 | GO:0009081 | branched-chain amino acid metabolic proc... | 37 | 4 | 0.15 | 1.70E-05 |
| 89 | GO:0010035 | response to inorganic substance | 2535 | 25 | 10.52 | 3.80E-05 |
| 90 | GO:0019747 | regulation of isoprenoid metabolic proce... | 17 | 3 | 0.07 | 4.60E-05 |
| 91 | GO:0009686 | gibberellin biosynthetic process | 96 | 5 | 0.4 | 5.10E-05 |
| 92 | GO:0016102 | diterpenoid biosynthetic process | 97 | 5 | 0.4 | 5.40E-05 |
| 93 | GO:0009617 | response to bacterium | 1124 | 15 | 4.67 | 6.70E-05 |
| 94 | GO:0044248 | cellular catabolic process | 2666 | 25 | 11.07 | 8.70E-05 |

| | | | | | | |
|---|---|---|---|---|---|---|
| 95 | GO:0008643 | carbohydrate transport | 60 | 4 | 0.25 | 0.00012 |
| 96 | GO:0009685 | gibberellin metabolic process | 115 | 5 | 0.48 | 0.00012 |
| 97 | GO:0016101 | diterpenoid metabolic process | 122 | 5 | 0.51 | 0.00016 |
| 98 | GO:0044282 | small molecule catabolic process | 1092 | 14 | 4.53 | 0.00018 |
| 99 | GO:0006979 | response to oxidative stress | 842 | 12 | 3.5 | 0.00021 |
| 100 | GO:0008299 | isoprenoid biosynthetic process | 718 | 11 | 2.98 | 0.00021 |
| 101 | GO:0009056 | catabolic process | 3010 | 26 | 12.5 | 0.00023 |
| 102 | GO:0046890 | regulation of lipid biosynthetic process | 33 | 3 | 0.14 | 0.00035 |
| 103 | GO:0009414 | response to water deprivation | 764 | 11 | 3.17 | 0.00035 |
| 104 | GO:0006720 | isoprenoid metabolic process | 767 | 11 | 3.18 | 0.00036 |
| 105 | GO:0046700 | heterocycle catabolic process | 897 | 12 | 3.72 | 0.00037 |
| 106 | GO:0006970 | response to osmotic stress | 1616 | 17 | 6.71 | 0.00038 |
| 107 | GO:0009415 | response to water | 777 | 11 | 3.23 | 0.0004 |
| 108 | GO:1901361 | organic cyclic compound catabolic proces... | 911 | 12 | 3.78 | 0.00042 |
| 109 | GO:0009607 | response to biotic stimulus | 2610 | 23 | 10.84 | 0.00043 |
| 110 | GO:0043207 | response to external biotic stimulus | 2610 | 23 | 10.84 | 0.00043 |
| 111 | GO:0051707 | response to other organism | 2610 | 23 | 10.84 | 0.00043 |
| 112 | GO:0048232 | male gamete generation | 90 | 4 | 0.37 | 0.00055 |
| 113 | GO:0009605 | response to external stimulus | 3770 | 29 | 15.65 | 0.00066 |
| 114 | GO:0046885 | regulation of hormone biosynthetic proce... | 42 | 3 | 0.17 | 0.00071 |
| 115 | GO:0048506 | regulation of timing of meristematic pha... | 48 | 3 | 0.2 | 0.00105 |
| 116 | GO:0048510 | regulation of timing of transition from ... | 48 | 3 | 0.2 | 0.00105 |
| 117 | GO:0098542 | defense response to other organism | 1935 | 18 | 8.03 | 0.00106 |
| 118 | GO:0044270 | cellular nitrogen compound catabolic pro... | 890 | 11 | 3.69 | 0.00123 |
| 119 | GO:0019439 | aromatic compound catabolic process | 893 | 11 | 3.71 | 0.00126 |

| | | | | | | |
|---|---|---|---|---|---|---|
| 120 | GO:0071470 | cellular response to osmotic stress | 53 | 3 | 0.22 | 0.00141 |
| 121 | GO:0016114 | terpenoid biosynthetic process | 402 | 7 | 1.67 | 0.00148 |
| 122 | GO:0010106 | cellular response to iron ion starvation | 203 | 5 | 0.84 | 0.00162 |
| 123 | GO:1901564 | organonitrogen compound metabolic proces... | 7265 | 45 | 30.16 | 0.00188 |
| 124 | GO:0044237 | cellular metabolic process | 13631 | 73 | 56.59 | 0.00198 |
| 125 | GO:0006879 | cellular iron ion homeostasis | 215 | 5 | 0.89 | 0.00208 |
| 126 | GO:0032350 | regulation of hormone metabolic process | 61 | 3 | 0.25 | 0.00211 |
| 127 | GO:0000302 | response to reactive oxygen species | 561 | 8 | 2.33 | 0.00242 |
| 128 | GO:0006721 | terpenoid metabolic process | 451 | 7 | 1.87 | 0.00283 |
| 129 | GO:0040034 | regulation of development, heterochronic | 68 | 3 | 0.28 | 0.00287 |
| 130 | GO:0042542 | response to hydrogen peroxide | 338 | 6 | 1.4 | 0.00293 |
| 131 | GO:1901565 | organonitrogen compound catabolic proces... | 1310 | 13 | 5.44 | 0.00316 |
| 132 | GO:0008152 | metabolic process | 15048 | 78 | 62.47 | 0.00316 |
| 133 | GO:0010565 | regulation of cellular ketone metabolic ... | 72 | 3 | 0.3 | 0.00338 |
| 134 | GO:0010150 | leaf senescence | 148 | 4 | 0.61 | 0.00343 |
| 135 | GO:0090693 | plant organ senescence | 148 | 4 | 0.61 | 0.00343 |
| 136 | GO:0055072 | iron ion homeostasis | 244 | 5 | 1.01 | 0.00357 |
| 137 | GO:0009611 | response to wounding | 619 | 8 | 2.57 | 0.00438 |
| 138 | GO:0009408 | response to heat | 497 | 7 | 2.06 | 0.0048 |
| 139 | GO:0044262 | cellular carbohydrate metabolic process | 1066 | 11 | 4.43 | 0.00491 |
| 140 | GO:0019216 | regulation of lipid metabolic process | 83 | 3 | 0.34 | 0.00504 |
| 141 | GO:0006952 | defense response | 2793 | 21 | 11.6 | 0.00539 |
| 142 | GO:0031668 | cellular response to extracellular stimu... | 642 | 8 | 2.67 | 0.00543 |
| 143 | GO:0071496 | cellular response to external stimulus | 644 | 8 | 2.67 | 0.00553 |
| 144 | GO:0009628 | response to abiotic stimulus | 4939 | 32 | 20.5 | 0.00561 |

| | | | | | | |
|---|---|---|---|---|---|---|
| 145 | GO:0044249 | cellular biosynthetic process | 9247 | 52 | 38.39 | 0.00616 |
| 146 | GO:0006560 | proline metabolic process | 29 | 2 | 0.12 | 0.00645 |
| 147 | GO:0046916 | cellular transition metal ion homeostasi... | 281 | 5 | 1.17 | 0.00645 |
| 148 | GO:0046677 | response to antibiotic | 1281 | 12 | 5.32 | 0.00709 |
| 149 | GO:0006546 | glycine catabolic process | 95 | 3 | 0.39 | 0.00732 |
| 150 | GO:0009991 | response to extracellular stimulus | 682 | 8 | 2.83 | 0.00771 |
| 151 | GO:0009071 | serine family amino acid catabolic proce... | 97 | 3 | 0.4 | 0.00775 |
| 152 | GO:0009725 | response to hormone | 3457 | 24 | 14.35 | 0.00778 |
| 153 | GO:0009719 | response to endogenous stimulus | 3463 | 24 | 14.38 | 0.00794 |
| 154 | GO:0007568 | aging | 296 | 5 | 1.23 | 0.00799 |
| 155 | GO:0042135 | neurotransmitter catabolic process | 100 | 3 | 0.42 | 0.00843 |
| 156 | GO:1901576 | organic substance biosynthetic process | 9857 | 54 | 40.92 | 0.00887 |
| 157 | GO:0009838 | abscission | 102 | 3 | 0.42 | 0.00889 |
| 158 | GO:0006544 | glycine metabolic process | 104 | 3 | 0.43 | 0.00938 |

**Supplementary Table 9**

Gene Ontology (GO) Enrichment Analysis of significantly differentially expressed genes ($p<0.05$) in the floret tissue of both mutant lines (above) using TopGo (Alexa and Rahnenführer, 2009). The gene set analysis statistically compared the representation of GO (Gene Ontology) terms ,which were first assigned to the *Brassica oleracea* gene models using the *Arabidopsis thaliana* org.At.tair.db library (version 3.2.3) (Carlson, 2016), in the gene set above to that of the 'expected' value, to determine those to be considered 'over-represented' using the BP parameter with a node size of 10, then confirming the statistical significance of this using Fisher's exact test. This was performed in RStudio v.1.2.5042 (Racine, 2012).

| | GO.ID | Term | Annotated | Significant | Expected | classicFisher |
|---|---|---|---|---|---|---|
| 1 | GO:0016144 | S-glycoside biosynthetic process | 300 | 25 | 0.82 | 1.60E-30 |
| 2 | GO:0019758 | glycosinolate biosynthetic process | 300 | 25 | 0.82 | 1.60E-30 |
| 3 | GO:0019761 | glucosinolate biosynthetic process | 300 | 25 | 0.82 | 1.60E-30 |
| 4 | GO:1901659 | glycosyl compound biosynthetic process | 311 | 25 | 0.85 | 3.90E-30 |
| 5 | GO:0016143 | S-glycoside metabolic process | 368 | 25 | 1.01 | 2.80E-28 |
| 6 | GO:0019757 | glycosinolate metabolic process | 368 | 25 | 1.01 | 2.80E-28 |
| 7 | GO:0019760 | glucosinolate metabolic process | 368 | 25 | 1.01 | 2.80E-28 |
| 8 | GO:1901657 | glycosyl compound metabolic process | 412 | 25 | 1.13 | 4.70E-27 |
| 9 | GO:0044550 | secondary metabolite biosynthetic proces... | 546 | 26 | 1.49 | 2.00E-25 |
| 10 | GO:0042435 | indole-containing compound biosynthetic ... | 236 | 18 | 0.65 | 2.50E-21 |
| 11 | GO:0009684 | indoleacetic acid biosynthetic process | 193 | 17 | 0.53 | 3.00E-21 |
| 12 | GO:0009683 | indoleacetic acid metabolic process | 195 | 17 | 0.53 | 3.60E-21 |
| 13 | GO:0044272 | sulfur compound biosynthetic process | 845 | 26 | 2.31 | 1.20E-20 |
| 14 | GO:0042430 | indole-containing compound metabolic pro... | 273 | 18 | 0.75 | 3.50E-20 |
| 15 | GO:0009851 | auxin biosynthetic process | 223 | 17 | 0.61 | 3.60E-20 |
| 16 | GO:0019748 | secondary metabolic process | 1119 | 28 | 3.06 | 6.80E-20 |
| 17 | GO:0009850 | auxin metabolic process | 253 | 17 | 0.69 | 3.10E-19 |
| 18 | GO:0034754 | cellular hormone metabolic process | 307 | 17 | 0.84 | 8.30E-18 |
| 19 | GO:0006790 | sulfur compound metabolic process | 1232 | 26 | 3.37 | 1.20E-16 |

| | | | | | | |
|---|---|---|---|---|---|---|
| 20 | GO:1901137 | carbohydrate derivative biosynthetic pro... | 1538 | 25 | 4.2 | 2.00E-13 |
| 21 | GO:1901605 | alpha-amino acid metabolic process | 936 | 20 | 2.56 | 6.10E-13 |
| 22 | GO:0006520 | cellular amino acid metabolic process | 1185 | 20 | 3.24 | 4.30E-11 |
| 23 | GO:0006569 | tryptophan catabolic process | 130 | 9 | 0.36 | 9.10E-11 |
| 24 | GO:0046218 | indolalkylamine catabolic process | 130 | 9 | 0.36 | 9.10E-11 |
| 25 | GO:0042436 | indole-containing compound catabolic pro... | 135 | 9 | 0.37 | 1.30E-10 |
| 26 | GO:0009074 | aromatic amino acid family catabolic pro... | 139 | 9 | 0.38 | 1.70E-10 |
| 27 | GO:0010439 | regulation of glucosinolate biosynthetic... | 16 | 5 | 0.04 | 5.70E-10 |
| 28 | GO:0006568 | tryptophan metabolic process | 161 | 9 | 0.44 | 6.20E-10 |
| 29 | GO:0006586 | indolalkylamine metabolic process | 161 | 9 | 0.44 | 6.20E-10 |
| 30 | GO:1901135 | carbohydrate derivative metabolic proces... | 2247 | 25 | 6.14 | 7.00E-10 |
| 31 | GO:1901607 | alpha-amino acid biosynthetic process | 672 | 14 | 1.84 | 3.80E-09 |
| 32 | GO:1901566 | organonitrogen compound biosynthetic pro... | 3504 | 30 | 9.58 | 4.20E-09 |
| 33 | GO:0042446 | hormone biosynthetic process | 1385 | 19 | 3.79 | 4.50E-09 |
| 34 | GO:0009310 | amine catabolic process | 209 | 9 | 0.57 | 6.10E-09 |
| 35 | GO:0042402 | cellular biogenic amine catabolic proces... | 209 | 9 | 0.57 | 6.10E-09 |
| 36 | GO:0019344 | cysteine biosynthetic process | 375 | 11 | 1.03 | 6.30E-09 |
| 37 | GO:0006534 | cysteine metabolic process | 377 | 11 | 1.03 | 6.70E-09 |
| 38 | GO:0009070 | serine family amino acid biosynthetic pr... | 381 | 11 | 1.04 | 7.50E-09 |
| 39 | GO:1900376 | regulation of secondary metabolite biosy... | 28 | 5 | 0.08 | 1.30E-08 |
| 40 | GO:0042445 | hormone metabolic process | 1548 | 19 | 4.23 | 2.70E-08 |
| 41 | GO:0008652 | cellular amino acid biosynthetic process | 788 | 14 | 2.15 | 2.80E-08 |
| 42 | GO:0042762 | regulation of sulfur metabolic process | 36 | 5 | 0.1 | 4.80E-08 |
| 43 | GO:0043436 | oxoacid metabolic process | 3444 | 28 | 9.42 | 5.30E-08 |
| 44 | GO:0006082 | organic acid metabolic process | 3445 | 28 | 9.42 | 5.40E-08 |

| | | | | | | |
|---|---|---|---|---|---|---|
| 45 | GO:0009069 | serine family amino acid metabolic proce... | 463 | 11 | 1.27 | 5.50E-08 |
| 46 | GO:1901606 | alpha-amino acid catabolic process | 275 | 9 | 0.75 | 6.50E-08 |
| 47 | GO:0072330 | monocarboxylic acid biosynthetic process | 1471 | 18 | 4.02 | 7.00E-08 |
| 48 | GO:0009063 | cellular amino acid catabolic process | 279 | 9 | 0.76 | 7.40E-08 |
| 49 | GO:0006576 | cellular biogenic amine metabolic proces... | 281 | 9 | 0.77 | 7.90E-08 |
| 50 | GO:0010675 | regulation of cellular carbohydrate meta... | 41 | 5 | 0.11 | 9.30E-08 |
| 51 | GO:0044106 | cellular amine metabolic process | 296 | 9 | 0.81 | 1.20E-07 |
| 52 | GO:0043255 | regulation of carbohydrate biosynthetic ... | 44 | 5 | 0.12 | 1.30E-07 |
| 53 | GO:0000097 | sulfur amino acid biosynthetic process | 525 | 11 | 1.44 | 1.90E-07 |
| 54 | GO:0016053 | organic acid biosynthetic process | 2157 | 21 | 5.9 | 2.20E-07 |
| 55 | GO:0046394 | carboxylic acid biosynthetic process | 2157 | 21 | 5.9 | 2.20E-07 |
| 56 | GO:0043455 | regulation of secondary metabolic proces... | 50 | 5 | 0.14 | 2.60E-07 |
| 57 | GO:0010817 | regulation of hormone levels | 1791 | 19 | 4.9 | 2.60E-07 |
| 58 | GO:0000096 | sulfur amino acid metabolic process | 598 | 11 | 1.63 | 7.00E-07 |
| 59 | GO:0009636 | response to toxic substance | 748 | 12 | 2.05 | 8.80E-07 |
| 60 | GO:0006109 | regulation of carbohydrate metabolic pro... | 70 | 5 | 0.19 | 1.40E-06 |
| 61 | GO:0009072 | aromatic amino acid family metabolic pro... | 406 | 9 | 1.11 | 1.70E-06 |
| 62 | GO:0009308 | amine metabolic process | 408 | 9 | 1.12 | 1.80E-06 |
| 63 | GO:0044283 | small molecule biosynthetic process | 2481 | 21 | 6.78 | 2.20E-06 |
| 64 | GO:0009098 | leucine biosynthetic process | 13 | 3 | 0.04 | 5.50E-06 |
| 65 | GO:0062012 | regulation of small molecule metabolic p... | 95 | 5 | 0.26 | 6.40E-06 |
| 66 | GO:0065008 | regulation of biological quality | 2914 | 22 | 7.97 | 7.60E-06 |
| 67 | GO:0006950 | response to stress | 6643 | 36 | 18.16 | 8.80E-06 |
| 68 | GO:0044281 | small molecule metabolic process | 4793 | 29 | 13.1 | 1.40E-05 |
| 69 | GO:0042742 | defense response to bacterium | 826 | 11 | 2.26 | 1.50E-05 |

| | | | | | | |
|---|---|---|---|---|---|---|
| 70 | GO:0032787 | monocarboxylic acid metabolic process | 2388 | 19 | 6.53 | 1.80E-05 |
| 71 | GO:0000103 | sulfate assimilation | 20 | 3 | 0.05 | 2.20E-05 |
| 72 | GO:1901564 | organonitrogen compound metabolic proces... | 7265 | 37 | 19.86 | 2.70E-05 |
| 73 | GO:0006551 | leucine metabolic process | 23 | 3 | 0.06 | 3.40E-05 |
| 74 | GO:0019752 | carboxylic acid metabolic process | 3262 | 22 | 8.92 | 4.50E-05 |
| 75 | GO:0009617 | response to bacterium | 1124 | 12 | 3.07 | 5.30E-05 |
| 76 | GO:0009082 | branched-chain amino acid biosynthetic p... | 27 | 3 | 0.07 | 5.50E-05 |
| 77 | GO:0009644 | response to high light intensity | 359 | 7 | 0.98 | 5.80E-05 |
| 78 | GO:0009404 | toxin metabolic process | 386 | 7 | 1.06 | 9.20E-05 |
| 79 | GO:0016054 | organic acid catabolic process | 677 | 9 | 1.85 | 9.70E-05 |
| 80 | GO:0046395 | carboxylic acid catabolic process | 677 | 9 | 1.85 | 9.70E-05 |
| 81 | GO:0009081 | branched-chain amino acid metabolic proc... | 37 | 3 | 0.1 | 0.00014 |
| 82 | GO:0044262 | cellular carbohydrate metabolic process | 1066 | 11 | 2.91 | 0.00015 |
| 83 | GO:0050896 | response to stimulus | 10855 | 46 | 29.68 | 0.00017 |
| 84 | GO:0009642 | response to light intensity | 476 | 7 | 1.3 | 0.00033 |
| 85 | GO:0042542 | response to hydrogen peroxide | 338 | 6 | 0.92 | 0.00033 |
| 86 | GO:0019953 | sexual reproduction | 496 | 7 | 1.36 | 0.00042 |
| 87 | GO:0009408 | response to heat | 497 | 7 | 1.36 | 0.00043 |
| 88 | GO:0009407 | toxin catabolic process | 359 | 6 | 0.98 | 0.00046 |
| 89 | GO:0044703 | multi-organism reproductive process | 507 | 7 | 1.39 | 0.00048 |
| 90 | GO:0051704 | multi-organism process | 3662 | 21 | 10.01 | 0.00067 |
| 91 | GO:0098754 | detoxification | 389 | 6 | 1.06 | 0.0007 |
| 92 | GO:0044270 | cellular nitrogen compound catabolic pro... | 890 | 9 | 2.43 | 0.00072 |
| 93 | GO:0019439 | aromatic compound catabolic process | 893 | 9 | 2.44 | 0.00074 |
| 94 | GO:0048544 | recognition of pollen | 15 | 2 | 0.04 | 0.00076 |

| | | | | | | |
|---|---|---|---|---|---|---|
| 95 | GO:0046700 | heterocycle catabolic process | 897 | 9 | 2.45 | 0.00076 |
| 96 | GO:1901361 | organic cyclic compound catabolic proces... | 911 | 9 | 2.49 | 0.00085 |
| 97 | GO:0008037 | cell recognition | 16 | 2 | 0.04 | 0.00086 |
| 98 | GO:0006457 | protein folding | 431 | 6 | 1.18 | 0.00118 |
| 99 | GO:0009987 | cellular process | 17469 | 61 | 47.76 | 0.00154 |
| 100 | GO:0009607 | response to biotic stimulus | 2610 | 16 | 7.14 | 0.00169 |
| 101 | GO:0043207 | response to external biotic stimulus | 2610 | 16 | 7.14 | 0.00169 |
| 102 | GO:0051707 | response to other organism | 2610 | 16 | 7.14 | 0.00169 |
| 103 | GO:1901576 | organic substance biosynthetic process | 9857 | 40 | 26.95 | 0.00192 |
| 104 | GO:0048232 | male gamete generation | 90 | 3 | 0.25 | 0.00195 |
| 105 | GO:0044248 | cellular catabolic process | 2666 | 16 | 7.29 | 0.0021 |
| 106 | GO:0006979 | response to oxidative stress | 842 | 8 | 2.3 | 0.00213 |
| 107 | GO:0098542 | defense response to other organism | 1935 | 13 | 5.29 | 0.00219 |
| 108 | GO:0009875 | pollen-pistil interaction | 28 | 2 | 0.08 | 0.00266 |
| 109 | GO:0009058 | biosynthetic process | 10049 | 40 | 27.47 | 0.00286 |
| 110 | GO:0044282 | small molecule catabolic process | 1092 | 9 | 2.99 | 0.00294 |
| 111 | GO:0065007 | biological regulation | 9405 | 38 | 25.71 | 0.00304 |
| 112 | GO:0006952 | defense response | 2793 | 16 | 7.64 | 0.00335 |
| 113 | GO:0000302 | response to reactive oxygen species | 561 | 6 | 1.53 | 0.00439 |
| 114 | GO:0009813 | flavonoid biosynthetic process | 404 | 5 | 1.1 | 0.00505 |
| 115 | GO:0016051 | carbohydrate biosynthetic process | 1417 | 10 | 3.87 | 0.0052 |
| 116 | GO:0008152 | metabolic process | 15048 | 53 | 41.14 | 0.00545 |
| 117 | GO:0044271 | cellular nitrogen compound biosynthetic ... | 5604 | 25 | 15.32 | 0.00663 |
| 118 | GO:0018130 | heterocycle biosynthetic process | 5004 | 23 | 13.68 | 0.00669 |
| 119 | GO:0009056 | catabolic process | 3010 | 16 | 8.23 | 0.0069 |

| | | | | | | |
|---|---|---|---|---|---|---|
| 120 | GO:0019252 | starch biosynthetic process | 280 | 4 | 0.77 | 0.00735 |
| 121 | GO:0009812 | flavonoid metabolic process | 446 | 5 | 1.22 | 0.0076 |
| 122 | GO:0010150 | leaf senescence | 148 | 3 | 0.4 | 0.00784 |
| 123 | GO:0090693 | plant organ senescence | 148 | 3 | 0.4 | 0.00784 |
| 124 | GO:0006098 | pentose-phosphate shunt | 290 | 4 | 0.79 | 0.00829 |
| 125 | GO:0051156 | glucose 6-phosphate metabolic process | 290 | 4 | 0.79 | 0.00829 |
| 126 | GO:0032270 | positive regulation of cellular protein ... | 51 | 2 | 0.14 | 0.00863 |
| 127 | GO:0042221 | response to chemical | 6670 | 28 | 18.24 | 0.00887 |
| 128 | GO:0009416 | response to light stimulus | 2039 | 12 | 5.57 | 0.00923 |
| 129 | GO:1901565 | organonitrogen compound catabolic proces... | 1310 | 9 | 3.58 | 0.00943 |
| 130 | GO:0006739 | NADP metabolic process | 302 | 4 | 0.83 | 0.00953 |
| 131 | GO:0051247 | positive regulation of protein metabolic... | 54 | 2 | 0.15 | 0.00964 |
| 132 | GO:1901362 | organic cyclic compound biosynthetic pro... | 5771 | 25 | 15.78 | 0.00966 |

**Supplementary Table 10**

All statistical analyses relating to vegetative leaf tissues as performed in Graphpad Prism v8.2.0. M1 = *bolC2.myb28-1*$^{ge}$; M2 = *bolC2.myb28-2*$^{ge}$

| Vegetative Leaf Glucoraphanin | | | | | | | | |
|---|---|---|---|---|---|---|---|---|
| Tukey's multiple comparisons test | Mean Diff. | 95.00% CI of diff. | Significant? | Summary | Adjusted P Value | | | |
| WT vs. M1 | 0.9376 | 0.4519 to 1.423 | Yes | *** | 0.0003 | B-C | | |
| WT vs. M2 | 0.9329 | 0.4202 to 1.446 | Yes | *** | 0.0005 | B-D | | |
| M1 vs. M2 | -0.00468 | -0.4691 to 0.4598 | No | ns | 0.9996 | C-D | | |
| Test details | Mean 1 | Mean 2 | Mean Diff. | SE of diff. | n1 | n2 | q | DF |
| WT vs. M1 | 0.9705 | 0.03289 | 0.9376 | 0.1912 | 6 | 9 | 6.935 | 19 |
| WT vs. M2 | 0.9705 | 0.03757 | 0.9329 | 0.2018 | 6 | 7 | 6.537 | 19 |
| M1 vs. M2 | 0.03289 | 0.03757 | -0.00468 | 0.1828 | 9 | 7 | 0.03622 | 19 |
| Vegetative Leaf Glucoiberin | | | | | | | | |
| Tukey's multiple comparisons test | Mean Diff. | 95.00% CI of diff. | Significant? | Summary | Adjusted P Value | | | |
| WT vs. M1 | -0.1877 | -0.5054 to 0.1301 | No | ns | 0.313 | B-C | | |
| WT vs. M2 | -0.02738 | -0.3628 to 0.3081 | No | ns | 0.9766 | B-D | | |
| M1 vs. M2 | 0.1603 | -0.1436 to 0.4641 | No | ns | 0.3912 | C-D | | |
| Test details | Mean 1 | Mean 2 | Mean Diff. | SE of diff. | n1 | n2 | q | DF |
| WT vs. M1 | 0.5353 | 0.723 | -0.1877 | 0.1251 | 6 | 9 | 2.122 | 19 |
| WT vs. M2 | 0.5353 | 0.5627 | -0.02738 | 0.132 | 6 | 7 | 0.2933 | 19 |
| M1 vs. M2 | 0.723 | 0.5627 | 0.1603 | 0.1196 | 9 | 7 | 1.895 | 19 |
| Vegetative Leaf Hydroxyindolylmethyl | | | | | | | | |
| Tukey's multiple comparisons test | Mean Diff. | 95.00% CI of diff. | Significant? | Summary | Adjusted P Value | | | |
| WT vs. M1 | 0.06728 | -0.05118 to 0.1857 | No | ns | 0.3399 | B-C | | |
| WT vs. M2 | 0.06702 | -0.05802 to 0.1921 | No | ns | 0.3801 | B-D | | |
| M1 vs. M2 | -0.00025 | -0.1135 to 0.1130 | No | ns | >0.9999 | C-D | | |

| | | | | | | | | |
|---|---|---|---|---|---|---|---|---|
| Test details | Mean 1 | Mean 2 | Mean Diff. | SE of diff. | n1 | n2 | q | DF |
| WT vs. M1 | 0.07017 | 0.002889 | 0.06728 | 0.04663 | 6 | 9 | 2.04 | 19 |
| WT vs. M2 | 0.07017 | 0.003143 | 0.06702 | 0.04922 | 6 | 7 | 1.926 | 19 |
| M1 vs. M2 | 0.002889 | 0.003143 | -0.00025 | 0.04459 | 9 | 7 | 0.008056 | 19 |
| Vegetative Leaf Indolylmethyl | | | | | | | | |
| Tukey's multiple comparisons test | Mean Diff. | 95.00% CI of diff. | Significant? | Summary | Adjusted P Value | | | |
| WT vs. M1 | 0.2751 | -0.6350 to 1.185 | No | ns | 0.7267 | B-C | | |
| WT vs. M2 | 0.2203 | -0.7404 to 1.181 | No | ns | 0.8309 | B-D | | |
| M1 vs. M2 | -0.05478 | -0.9250 to 0.8155 | No | ns | 0.986 | C-D | | |
| Test details | Mean 1 | Mean 2 | Mean Diff. | SE of diff. | n1 | n2 | q | DF |
| WT vs. M1 | 1.001 | 0.7262 | 0.2751 | 0.3582 | 6 | 9 | 1.086 | 19 |
| WT vs. M2 | 1.001 | 0.781 | 0.2203 | 0.3782 | 6 | 7 | 0.824 | 19 |
| M1 vs. M2 | 0.7262 | 0.781 | -0.05478 | 0.3425 | 9 | 7 | 0.2262 | 19 |
| Vegetative Leaf 1-methoxyindolylmethl | | | | | | | | |
| Tukey's multiple comparisons test | Mean Diff. | 95.00% CI of diff. | Significant? | Summary | Adjusted P Value | | | |
| WT vs. M1 | 0.01733 | -0.02402 to 0.05868 | No | ns | 0.5466 | B-C | | |
| WT vs. M2 | 0.02976 | -0.01389 to 0.07341 | No | ns | 0.2194 | B-D | | |
| M1 vs. M2 | 0.01243 | -0.02711 to 0.05197 | No | ns | 0.7084 | C-D | | |
| Test details | Mean 1 | Mean 2 | Mean Diff. | SE of diff. | n1 | n2 | q | DF |
| WT vs. M1 | 0.06633 | 0.049 | 0.01733 | 0.01628 | 6 | 9 | 1.506 | 19 |
| WT vs. M2 | 0.06633 | 0.03657 | 0.02976 | 0.01718 | 6 | 7 | 2.45 | 19 |
| M1 vs. M2 | 0.049 | 0.03657 | 0.01243 | 0.01556 | 9 | 7 | 1.129 | 19 |
| Vegetative Leaf 4-methoxyindolylmethyl | | | | | | | | |
| Tukey's multiple comparisons test | Mean Diff. | 95.00% CI of diff. | Significant? | Summary | Adjusted P Value | | | |

| | | | | | | | | |
|---|---|---|---|---|---|---|---|---|
| WT vs. M1 | 0.05306 | -0.04893 to 0.1550 | No | ns | 0.4009 | B-C | | |
| WT vs. M2 | 0.06655 | -0.04110 to 0.1742 | No | ns | 0.2822 | B-D | | |
| M1 vs. M2 | 0.01349 | -0.08402 to 0.1110 | No | ns | 0.9344 | C-D | | |
| Test details | Mean 1 | Mean 2 | Mean Diff. | SE of diff. | n1 | n2 | q | DF |
| WT vs. M1 | 0.1548 | 0.1018 | 0.05306 | 0.04014 | 6 | 9 | 1.869 | 19 |
| WT vs. M2 | 0.1548 | 0.08829 | 0.06655 | 0.04238 | 6 | 7 | 2.221 | 19 |
| M1 vs. M2 | 0.1018 | 0.08829 | 0.01349 | 0.03838 | 9 | 7 | 0.4971 | 19 |
| Vegetative Leaf Sulphate | | | | | | | | |
| Tukey's multiple comparisons test | Mean Diff. | 95.00% CI of diff. | Significant? | Summary | Adjusted P Value | | | |
| WT vs. M1 | 108.7 | -37.89 to 255.3 | No | ns | 0.1706 | B-C | | |
| WT vs. M2 | 88.84 | -65.90 to 243.6 | No | ns | 0.3324 | B-D | | |
| M1 vs. M2 | -19.86 | -160.0 to 120.3 | No | ns | 0.9313 | C-D | | |
| Test details | Mean 1 | Mean 2 | Mean Diff. | SE of diff. | n1 | n2 | q | DF |
| WT vs. M1 | 503.6 | 394.9 | 108.7 | 57.7 | 6 | 9 | 2.664 | 19 |
| WT vs. M2 | 503.6 | 414.7 | 88.84 | 60.91 | 6 | 7 | 2.063 | 19 |
| M1 vs. M2 | 394.9 | 414.7 | -19.86 | 55.17 | 9 | 7 | 0.509 | 19 |
| Vegetative Leaf SMCSO | | | | | | | | |
| Tukey's multiple comparisons test | | | | | | | | |
| Tukey's multiple comparisons test | Mean Diff. | 95.00% CI of diff. | Significant? | Summary | Adjusted P Value | | | |
| WT vs. M1 | 3.181 | -1.085 to 7.447 | No | ns | 0.1676 | B-C | | |
| WT vs. M2 | 1.041 | -3.462 to 5.545 | No | ns | 0.8284 | B-D | | |
| M1 vs. M2 | -2.14 | -6.219 to 1.940 | No | ns | 0.3952 | C-D | | |
| Test details | Mean 1 | Mean 2 | Mean Diff. | SE of diff. | n1 | n2 | q | DF |
| WT vs. M1 | 11.58 | 8.399 | 3.181 | 1.679 | 6 | 9 | 2.679 | 19 |
| WT vs. M2 | 11.58 | 10.54 | 1.041 | 1.773 | 6 | 7 | 0.8308 | 19 |
| M1 vs. M2 | 8.399 | 10.54 | -2.14 | 1.606 | 9 | 7 | 1.884 | 19 |

| | | | | | |
|---|---|---|---|---|---|
| Vegetative Leaf Gene Expression | | | | | |
| Tukey's multiple comparisons test | Mean Diff. | 95.00% CI of diff. | Significant? | Summary | Adjusted P Value |
| BCAT4 C5 | | | | | |
| WT vs. M1 | 70.75 | 54.11 to 87.38 | Yes | **** | <0.0001 |
| WT vs. M2 | 70.52 | 53.89 to 87.15 | Yes | **** | <0.0001 |
| M1 vs. M2 | -0.2239 | -16.86 to 16.41 | No | ns | 0.9994 |
| BAT5 C3 | | | | | |
| WT vs. M1 | 11.51 | -5.125 to 28.14 | No | ns | 0.2293 |
| WT vs. M2 | 11.51 | -5.119 to 28.14 | No | ns | 0.229 |
| M1 vs. M2 | 0.006143 | -16.63 to 16.64 | No | ns | >0.9999 |
| MAM3 C2 | | | | | |
| WT vs. M1 | 81.55 | 64.92 to 98.18 | Yes | **** | <0.0001 |
| WT vs. M2 | 81.56 | 64.93 to 98.19 | Yes | **** | <0.0001 |
| M1 vs. M2 | 0.006408 | -16.62 to 16.64 | No | ns | >0.9999 |
| IPMI | | | | | |
| WT vs. M1 | 69.04 | 52.41 to 85.68 | Yes | **** | <0.0001 |
| WT vs. M2 | 68.35 | 51.72 to 84.98 | Yes | **** | <0.0001 |
| M1 vs. M2 | -0.6977 | -17.33 to 15.93 | No | ns | 0.9945 |
| IMDH3 | | | | | |
| WT vs. M1 | 40.51 | 23.88 to 57.14 | Yes | **** | <0.0001 |
| WT vs. M2 | 40.53 | 23.90 to 57.16 | Yes | **** | <0.0001 |
| M1 vs. M2 | 0.02029 | -16.61 to 16.65 | No | ns | >0.9999 |
| CYP79F1 | | | | | |
| WT vs. M1 | 21.56 | 4.927 to 38.19 | Yes | ** | 0.0076 |

| | | | | | |
|---|---|---|---|---|---|
| WT vs. M2 | 21.56 | 4.932 to 38.20 | Yes | ** | 0.0076 |
| M1 vs. M2 | 0.005152 | -16.63 to 16.64 | No | ns | >0.9999 |
| CYP83A1 | | | | | |
| WT vs. M1 | 182.8 | 166.1 to 199.4 | Yes | **** | <0.0001 |
| WT vs. M2 | 185.1 | 168.5 to 201.7 | Yes | **** | <0.0001 |
| M1 vs. M2 | 2.303 | -14.33 to 18.93 | No | ns | 0.9413 |
| SUR1 | | | | | |
| WT vs. M1 | 107.1 | 90.52 to 123.8 | Yes | **** | <0.0001 |
| WT vs. M2 | 106.2 | 89.61 to 122.9 | Yes | **** | <0.0001 |
| M1 vs. M2 | -0.9098 | -17.54 to 15.72 | No | ns | 0.9906 |
| UGT74C1 | | | | | |
| WT vs. M1 | 29.69 | 13.06 to 46.33 | Yes | *** | 0.0002 |
| WT vs. M2 | 28.23 | 11.60 to 44.86 | Yes | *** | 0.0004 |
| M1 vs. M2 | -1.466 | -18.10 to 15.17 | No | ns | 0.9758 |
| SOT17 | | | | | |
| WT vs. M1 | 31.42 | 14.79 to 48.05 | Yes | **** | <0.0001 |
| WT vs. M2 | 33.08 | 16.44 to 49.71 | Yes | **** | <0.0001 |
| M1 vs. M2 | 1.656 | -14.98 to 18.29 | No | ns | 0.9692 |
| SOT18 | | | | | |
| WT vs. M1 | 12.34 | -4.294 to 28.97 | No | ns | 0.1852 |
| WT vs. M2 | 12.32 | -4.308 to 28.95 | No | ns | 0.1859 |
| M1 vs. M2 | -0.01394 | -16.65 to 16.62 | No | ns | >0.9999 |
| FMO GS-OX2 | | | | | |
| WT vs. M1 | 53.71 | 37.08 to 70.35 | Yes | **** | <0.0001 |

| | | | | | | | | |
|---|---|---|---|---|---|---|---|---|
| WT vs. M2 | 61.19 | 44.56 to 77.82 | Yes | **** | <0.0001 | | | |
| M1 vs. M2 | 7.476 | -9.156 to 24.11 | No | ns | 0.5321 | | | |
| Test details | Mean 1 | Mean 2 | Mean Diff. | SE of diff. | N1 | N2 | q | DF |
| BCAT4 C5 | | | | | | | | |
| WT vs. M1 | 72.2 | 1.454 | 70.75 | 6.95 | 3 | 3 | 14.4 | 72 |
| WT vs. M2 | 72.2 | 1.678 | 70.52 | 6.95 | 3 | 3 | 14.35 | 72 |
| M1 vs. M2 | 1.454 | 1.678 | -0.2239 | 6.95 | 3 | 3 | 0.04557 | 72 |
| BAT5 C3 | | | | | | | | |
| WT vs. M1 | 11.51 | 0.006143 | 11.51 | 6.95 | 3 | 3 | 2.342 | 72 |
| WT vs. M2 | 11.51 | 0 | 11.51 | 6.95 | 3 | 3 | 2.343 | 72 |
| M1 vs. M2 | 0.006143 | 0 | 0.006143 | 6.95 | 3 | 3 | 0.00125 | 72 |
| MAM3 C2 | | | | | | | | |
| WT vs. M1 | 81.57 | 0.01509 | 81.55 | 6.95 | 3 | 3 | 16.6 | 72 |
| WT vs. M2 | 81.57 | 0.008679 | 81.56 | 6.95 | 3 | 3 | 16.6 | 72 |
| M1 vs. M2 | 0.01509 | 0.008679 | 0.006408 | 6.95 | 3 | 3 | 0.001304 | 72 |
| IPMI | | | | | | | | |
| WT vs. M1 | 69.27 | 0.2232 | 69.04 | 6.95 | 3 | 3 | 14.05 | 72 |
| WT vs. M2 | 69.27 | 0.9209 | 68.35 | 6.95 | 3 | 3 | 13.91 | 72 |
| M1 vs. M2 | 0.2232 | 0.9209 | -0.6977 | 6.95 | 3 | 3 | 0.142 | 72 |
| IMDH3 | | | | | | | | |
| WT vs. M1 | 40.62 | 0.1097 | 40.51 | 6.95 | 3 | 3 | 8.244 | 72 |
| WT vs. M2 | 40.62 | 0.08939 | 40.53 | 6.95 | 3 | 3 | 8.248 | 72 |
| M1 vs. M2 | 0.1097 | 0.08939 | 0.02029 | 6.95 | 3 | 3 | 0.004128 | 72 |
| CYP79F1 | | | | | | | | |
| WT vs. M1 | 21.57 | 0.009165 | 21.56 | 6.95 | 3 | 3 | 4.387 | 72 |
| WT vs. M2 | 21.57 | 0.004013 | 21.56 | 6.95 | 3 | 3 | 4.388 | 72 |
| M1 vs. M2 | 0.009165 | 0.004013 | 0.005152 | 6.95 | 3 | 3 | 0.001048 | 72 |
| CYP83A1 | | | | | | | | |
| WT vs. M1 | 220 | 37.24 | 182.8 | 6.95 | 3 | 3 | 37.19 | 72 |

| | | | | | | | | |
|---|---|---|---|---|---|---|---|---|
| WT vs. M2 | 220 | 34.94 | 185.1 | 6.95 | 3 | 3 | 37.66 | 72 |
| M1 vs. M2 | 37.24 | 34.94 | 2.303 | 6.95 | 3 | 3 | 0.4686 | 72 |
| SUR1 | | | | | | | | |
| WT vs. M1 | 118.2 | 11.06 | 107.1 | 6.95 | 3 | 3 | 21.8 | 72 |
| WT vs. M2 | 118.2 | 11.97 | 106.2 | 6.95 | 3 | 3 | 21.62 | 72 |
| M1 vs. M2 | 11.06 | 11.97 | -0.9098 | 6.95 | 3 | 3 | 0.1851 | 72 |
| UGT74C1 | | | | | | | | |
| WT vs. M1 | 43.22 | 13.53 | 29.69 | 6.95 | 3 | 3 | 6.043 | 72 |
| WT vs. M2 | 43.22 | 15 | 28.23 | 6.95 | 3 | 3 | 5.744 | 72 |
| M1 vs. M2 | 13.53 | 15 | -1.466 | 6.95 | 3 | 3 | 0.2982 | 72 |
| SOT17 | | | | | | | | |
| WT vs. M1 | 42.73 | 11.31 | 31.42 | 6.95 | 3 | 3 | 6.394 | 72 |
| WT vs. M2 | 42.73 | 9.656 | 33.08 | 6.95 | 3 | 3 | 6.731 | 72 |
| M1 vs. M2 | 11.31 | 9.656 | 1.656 | 6.95 | 3 | 3 | 0.3371 | 72 |
| SOT18 | | | | | | | | |
| WT vs. M1 | 14.03 | 1.698 | 12.34 | 6.95 | 3 | 3 | 2.511 | 72 |
| WT vs. M2 | 14.03 | 1.712 | 12.32 | 6.95 | 3 | 3 | 2.508 | 72 |
| M1 vs. M2 | 1.698 | 1.712 | -0.01394 | 6.95 | 3 | 3 | 0.002836 | 72 |
| FMO GS-OX2 | | | | | | | | |
| WT vs. M1 | 112.3 | 58.61 | 53.71 | 6.95 | 3 | 3 | 10.93 | 72 |
| WT vs. M2 | 112.3 | 51.14 | 61.19 | 6.95 | 3 | 3 | 12.45 | 72 |
| M1 vs. M2 | 58.61 | 51.14 | 7.476 | 6.95 | 3 | 3 | 1.521 | 72 |

**Supplementary Table 11**
All statistical analyses relating to generative leaf tissues as performed in Graphpad Prism v8.2.0. M1 = *bolC2.myb28-1*$^{ge}$; M2 = *bolC2.myb28-2*$^{ge}$

| | | | | | | | | |
|---|---|---|---|---|---|---|---|---|
| Generative Leaf Glucoraphanin | | | | | | | | |
| Tukey's multiple comparisons test | Mean Diff. | 95.00% CI of diff. | Significant? | Summary | Adjusted P Value | | | |
| WT vs. M1 | 5.507 | 3.287 to 7.726 | Yes | **** | <0.0001 | B-C | | |
| WT vs. M2 | 5.519 | 3.212 to 7.825 | Yes | **** | <0.0001 | B-D | | |
| M1 vs. M2 | 0.01197 | -2.070 to 2.094 | No | ns | 0.9999 | C-D | | |
| Test details | Mean 1 | Mean 2 | Mean Diff. | SE of diff. | n1 | n2 | q | DF |
| WT vs. M1 | 5.719 | 0.2122 | 5.507 | 0.8319 | 4 | 6 | 9.361 | 12 |
| WT vs. M2 | 5.719 | 0.2002 | 5.519 | 0.8645 | 4 | 5 | 9.027 | 12 |
| M1 vs. M2 | 0.2122 | 0.2002 | 0.01197 | 0.7804 | 6 | 5 | 0.02169 | 12 |
| Generative Leaf Glucoiberin | | | | | | | | |
| Tukey's multiple comparisons test | Mean Diff. | 95.00% CI of diff. | Significant? | Summary | Adjusted P Value | | | |
| WT vs. M1 | 0.5123 | 0.1398 to 0.8847 | Yes | ** | 0.0083 | B-C | | |
| WT vs. M2 | 0.4872 | 0.1001 to 0.8742 | Yes | * | 0.0146 | B-D | | |
| M1 vs. M2 | -0.0251 | -0.3745 to 0.3243 | No | ns | 0.98 | C-D | | |
| Test details | Mean 1 | Mean 2 | Mean Diff. | SE of diff. | n1 | n2 | q | DF |
| WT vs. M1 | 1.121 | 0.6085 | 0.5123 | 0.1396 | 4 | 6 | 5.189 | 12 |
| WT vs. M2 | 1.121 | 0.6336 | 0.4872 | 0.1451 | 4 | 5 | 4.748 | 12 |
| M1 vs. M2 | 0.6085 | 0.6336 | -0.0251 | 0.131 | 6 | 5 | 0.271 | 12 |
| Generative Leaf Hydroxyindolylmethyl | | | | | | | | |
| Tukey's multiple comparisons test | Mean Diff. | 95.00% CI of diff. | Significant? | Summary | Adjusted P Value | | | |
| WT vs. M1 | -0.08492 | -0.4049 to 0.2351 | No | ns | 0.7635 | B-C | | |
| WT vs. M2 | -0.08865 | -0.4212 to 0.2439 | No | ns | 0.7616 | B-D | | |

| | | | | | | | | |
|---|---|---|---|---|---|---|---|---|
| M1 vs. M2 | -0.00373 | -0.3039 to 0.2964 | No | ns | 0.9994 | C-D | | |
| Test details | Mean 1 | Mean 2 | Mean Diff. | SE of diff. | n1 | n2 | q | DF |
| WT vs. M1 | 0.03475 | 0.1197 | -0.08492 | 0.1199 | 4 | 6 | 1.001 | 12 |
| WT vs. M2 | 0.03475 | 0.1234 | -0.08865 | 0.1246 | 4 | 5 | 1.006 | 12 |
| M1 vs. M2 | 0.1197 | 0.1234 | -0.00373 | 0.1125 | 6 | 5 | 0.04693 | 12 |
| Generative Leaf Indolylmethyl | | | | | | | | |
| Tukey's multiple comparisons test | Mean Diff. | 95.00% CI of diff. | Significant? | Summary | Adjusted P Value | | | |
| WT vs. M1 | 2.196 | 0.4888 to 3.903 | Yes | * | 0.0128 | B-C | | |
| WT vs. M2 | 2.731 | 0.9565 to 4.505 | Yes | ** | 0.0038 | B-D | | |
| M1 vs. M2 | 0.5347 | -1.067 to 2.136 | No | ns | 0.656 | C-D | | |
| Test details | Mean 1 | Mean 2 | Mean Diff. | SE of diff. | n1 | n2 | q | DF |
| WT vs. M1 | 5.607 | 3.411 | 2.196 | 0.6399 | 4 | 6 | 4.853 | 12 |
| WT vs. M2 | 5.607 | 2.877 | 2.731 | 0.665 | 4 | 5 | 5.807 | 12 |
| M1 vs. M2 | 3.411 | 2.877 | 0.5347 | 0.6003 | 6 | 5 | 1.26 | 12 |
| Generative Leaf 1-methoxyindolylmethl | | | | | | | | |
| Tukey's multiple comparisons test | Mean Diff. | 95.00% CI of diff. | Significant? | Summary | Adjusted P Value | | | |
| WT vs. M1 | 0.05175 | -0.05932 to 0.1628 | No | ns | 0.452 | B-C | | |
| WT vs. M2 | 0.03775 | -0.07768 to 0.1532 | No | ns | 0.667 | B-D | | |
| M1 vs. M2 | -0.014 | -0.1182 to 0.09020 | No | ns | 0.932 | C-D | | |
| Test details | Mean 1 | Mean 2 | Mean Diff. | SE of diff. | n1 | n2 | q | DF |
| WT vs. M1 | 0.1758 | 0.124 | 0.05175 | 0.04163 | 4 | 6 | 1.758 | 12 |
| WT vs. M2 | 0.1758 | 0.138 | 0.03775 | 0.04327 | 4 | 5 | 1.234 | 12 |
| M1 vs. M2 | 0.124 | 0.138 | -0.014 | 0.03906 | 6 | 5 | 0.5069 | 12 |
| Generative Leaf 4-methoxyindolylmethyl | | | | | | | | |

| Tukey's multiple comparisons test | Mean Diff. | 95.00% CI of diff. | Significant? | Summary | Adjusted P Value | | | |
|---|---|---|---|---|---|---|---|---|
| WT vs. M1 | 0.4701 | -0.2498 to 1.190 | No | ns | 0.2303 | B-C | | |
| WT vs. M2 | 0.6724 | -0.07582 to 1.421 | No | ns | 0.0799 | B-D | | |
| M1 vs. M2 | 0.2023 | -0.4731 to 0.8776 | No | ns | 0.7107 | C-D | | |
| Test details | Mean 1 | Mean 2 | Mean Diff. | SE of diff. | n1 | n2 | q | DF |
| WT vs. M1 | 1.267 | 0.7967 | 0.4701 | 0.2699 | 4 | 6 | 2.464 | 12 |
| WT vs. M2 | 1.267 | 0.5944 | 0.6724 | 0.2804 | 4 | 5 | 3.391 | 12 |
| M1 vs. M2 | 0.7967 | 0.5944 | 0.2023 | 0.2531 | 6 | 5 | 1.13 | 12 |
| Generative Leaf Sulphate | | | | | | | | |
| Tukey's multiple comparisons test | Mean Diff. | 95.00% CI of diff. | Significant? | Summary | Adjusted P Value | | | |
| WT vs. M1 | 93.81 | -222.1 to 409.8 | No | ns | 0.7147 | B-C | | |
| WT vs. M2 | 156.6 | -171.7 to 484.9 | No | ns | 0.4363 | B-D | | |
| M1 vs. M2 | 62.78 | -233.6 to 359.2 | No | ns | 0.8408 | C-D | | |
| Test details | Mean 1 | Mean 2 | Mean Diff. | SE of diff. | n1 | n2 | q | DF |
| WT vs. M1 | 438.6 | 344.8 | 93.81 | 118.4 | 4 | 6 | 1.12 | 12 |
| WT vs. M2 | 438.6 | 282.1 | 156.6 | 123.1 | 4 | 5 | 1.799 | 12 |
| M1 vs. M2 | 344.8 | 282.1 | 62.78 | 111.1 | 6 | 5 | 0.7992 | 12 |
| Generative Leaf SMCSO | | | | | | | | |
| Tukey's multiple comparisons test | Mean Diff. | 95.00% CI of diff. | Significant? | Summary | Adjusted P Value | | | |
| WT vs. M1 | 4.789 | -6.271 to 15.85 | No | ns | 0.5005 | B-C | | |
| WT vs. M2 | 3.512 | -7.983 to 15.01 | No | ns | 0.7012 | B-D | | |
| M1 vs. M2 | -1.278 | -11.65 to 9.098 | No | ns | 0.9425 | C-D | | |
| Test details | Mean 1 | Mean 2 | Mean Diff. | SE of diff. | n1 | n2 | q | DF |
| WT vs. M1 | 18.03 | 13.24 | 4.789 | 4.146 | 4 | 6 | 1.634 | 12 |
| WT vs. M2 | 18.03 | 14.52 | 3.512 | 4.308 | 4 | 5 | 1.153 | 12 |
| M1 vs. M2 | 13.24 | 14.52 | -1.278 | 3.889 | 6 | 5 | 0.4646 | 12 |

| Generative Leaf Gene Expression | | | | | |
|---|---|---|---|---|---|
| Tukey's multiple comparisons test | Mean Diff. | 95.00% CI of diff. | Significant? | Summary | Adjusted P Value |
| BCAT4 C5 | | | | | |
| WT vs. M1 | 73.89 | 55.29 to 92.50 | Yes | **** | <0.0001 |
| WT vs. M2 | 72.34 | 53.73 to 90.94 | Yes | **** | <0.0001 |
| M1 vs. M2 | -1.556 | -20.16 to 17.05 | No | ns | 0.9781 |
| BAT5 C3 | | | | | |
| WT vs. M1 | 10.27 | -8.333 to 28.88 | No | ns | 0.3872 |
| WT vs. M2 | 10.4 | -8.200 to 29.01 | No | ns | 0.3779 |
| M1 vs. M2 | 0.133 | -18.47 to 18.74 | No | ns | 0.9998 |
| MAM3 C2 | | | | | |
| WT vs. M1 | 72.74 | 54.14 to 91.35 | Yes | **** | <0.0001 |
| WT vs. M2 | 72.52 | 53.91 to 91.12 | Yes | **** | <0.0001 |
| M1 vs. M2 | -0.2232 | -18.83 to 18.38 | No | ns | 0.9995 |
| IPMI | | | | | |
| WT vs. M1 | 70.57 | 51.96 to 89.17 | Yes | **** | <0.0001 |
| WT vs. M2 | 70.43 | 51.83 to 89.04 | Yes | **** | <0.0001 |
| M1 vs. M2 | -0.135 | -18.74 to 18.47 | No | ns | 0.9998 |
| IMDH3 | | | | | |
| WT vs. M1 | 33.02 | 14.41 to 51.62 | Yes | *** | 0.0002 |
| WT vs. M2 | 33.05 | 14.44 to 51.65 | Yes | *** | 0.0002 |
| M1 vs. M2 | 0.03151 | -18.57 to 18.64 | No | ns | >0.9999 |
| CYP79F1 | | | | | |
| WT vs. M1 | 32.38 | 13.78 to 50.99 | Yes | *** | 0.0003 |

| | | | | | | | | |
|---|---|---|---|---|---|---|---|---|
| WT vs. M2 | 32.43 | 13.83 to 51.04 | Yes | *** | 0.0003 | | | |
| M1 vs. M2 | 0.04818 | -18.56 to 18.65 | No | ns | >0.9999 | | | |
| CYP83A1 | | | | | | | | |
| WT vs. M1 | 197.7 | 179.1 to 216.3 | Yes | **** | <0.0001 | | | |
| WT vs. M2 | 217.5 | 198.9 to 236.1 | Yes | **** | <0.0001 | | | |
| M1 vs. M2 | 19.75 | 1.150 to 38.36 | Yes | * | 0.0349 | | | |
| SUR1 | | | | | | | | |
| WT vs. M1 | 71.16 | 52.55 to 89.76 | Yes | **** | <0.0001 | | | |
| WT vs. M2 | 69.55 | 50.94 to 88.15 | Yes | **** | <0.0001 | | | |
| M1 vs. M2 | -1.609 | -20.21 to 17.00 | No | ns | 0.9766 | | | |
| UGT74C1 | | | | | | | | |
| WT vs. M1 | 23.06 | 4.459 to 41.67 | Yes | * | 0.0113 | | | |
| WT vs. M2 | 24.09 | 5.485 to 42.69 | Yes | ** | 0.0078 | | | |
| M1 vs. M2 | 1.026 | -17.58 to 19.63 | No | ns | 0.9904 | | | |
| SOT17 | | | | | | | | |
| WT vs. M1 | 10.84 | -7.765 to 29.45 | No | ns | 0.3483 | | | |
| WT vs. M2 | 11.54 | -7.066 to 30.14 | No | ns | 0.3036 | | | |
| M1 vs. M2 | 0.6987 | -17.91 to 19.30 | No | ns | 0.9955 | | | |
| SOT18 | | | | | | | | |
| WT vs. M1 | 11.01 | -7.600 to 29.61 | No | ns | 0.3374 | | | |
| WT vs. M2 | 10.42 | -8.188 to 29.02 | No | ns | 0.377 | | | |
| M1 vs. M2 | -0.5879 | -19.19 to 18.02 | No | ns | 0.9968 | | | |
| Test details | Mean 1 | Mean 2 | Mean Diff. | SE of diff. | N1 | N2 | q | DF |

| | | | | | | | | |
|---|---|---|---|---|---|---|---|---|
| BCAT4 C5 | | | | | | | | |
| WT vs. M1 | 76.21 | 2.317 | 73.89 | 7.76 | 3 | 3 | 13.47 | 66 |
| WT vs. M2 | 76.21 | 3.873 | 72.34 | 7.76 | 3 | 3 | 13.18 | 66 |
| M1 vs. M2 | 2.317 | 3.873 | -1.556 | 7.76 | 3 | 3 | 0.2835 | 66 |
| BAT5 C3 | | | | | | | | |
| WT vs. M1 | 10.76 | 0.4878 | 10.27 | 7.76 | 3 | 3 | 1.872 | 66 |
| WT vs. M2 | 10.76 | 0.3548 | 10.4 | 7.76 | 3 | 3 | 1.896 | 66 |
| M1 vs. M2 | 0.4878 | 0.3548 | 0.133 | 7.76 | 3 | 3 | 0.02424 | 66 |
| MAM3 C2 | | | | | | | | |
| WT vs. M1 | 73.35 | 0.604 | 72.74 | 7.76 | 3 | 3 | 13.26 | 66 |
| WT vs. M2 | 73.35 | 0.8272 | 72.52 | 7.76 | 3 | 3 | 13.22 | 66 |
| M1 vs. M2 | 0.604 | 0.8272 | -0.2232 | 7.76 | 3 | 3 | 0.04067 | 66 |
| IPMI | | | | | | | | |
| WT vs. M1 | 73.21 | 2.637 | 70.57 | 7.76 | 3 | 3 | 12.86 | 66 |
| WT vs. M2 | 73.21 | 2.772 | 70.43 | 7.76 | 3 | 3 | 12.84 | 66 |
| M1 vs. M2 | 2.637 | 2.772 | -0.135 | 7.76 | 3 | 3 | 0.0246 | 66 |
| IMDH3 | | | | | | | | |
| WT vs. M1 | 34.58 | 1.558 | 33.02 | 7.76 | 3 | 3 | 6.018 | 66 |
| WT vs. M2 | 34.58 | 1.527 | 33.05 | 7.76 | 3 | 3 | 6.023 | 66 |
| M1 vs. M2 | 1.558 | 1.527 | 0.03151 | 7.76 | 3 | 3 | 0.005742 | 66 |
| CYP79F1 | | | | | | | | |
| WT vs. M1 | 33.19 | 0.8106 | 32.38 | 7.76 | 3 | 3 | 5.902 | 66 |
| WT vs. M2 | 33.19 | 0.7625 | 32.43 | 7.76 | 3 | 3 | 5.911 | 66 |
| M1 vs. M2 | 0.8106 | 0.7625 | 0.04818 | 7.76 | 3 | 3 | 0.00878 | 66 |
| CYP83A1 | | | | | | | | |
| WT vs. M1 | 258.8 | 61.05 | 197.7 | 7.76 | 3 | 3 | 36.03 | 66 |
| WT vs. M2 | 258.8 | 41.3 | 217.5 | 7.76 | 3 | 3 | 39.63 | 66 |
| M1 vs. M2 | 61.05 | 41.3 | 19.75 | 7.76 | 3 | 3 | 3.6 | 66 |
| SUR1 | | | | | | | | |
| WT vs. M1 | 77.56 | 6.406 | 71.16 | 7.76 | 3 | 3 | 12.97 | 66 |
| WT vs. M2 | 77.56 | 8.016 | 69.55 | 7.76 | 3 | 3 | 12.68 | 66 |
| M1 vs. M2 | 6.406 | 8.016 | -1.609 | 7.76 | 3 | 3 | 0.2933 | 66 |

| | | | | | | | | |
|---|---|---|---|---|---|---|---|---|
| UGT74C1 | | | | | | | | |
| WT vs. M1 | 29.87 | 6.811 | 23.06 | 7.76 | 3 | 3 | 4.203 | 66 |
| WT vs. M2 | 29.87 | 5.785 | 24.09 | 7.76 | 3 | 3 | 4.39 | 66 |
| M1 vs. M2 | 6.811 | 5.785 | 1.026 | 7.76 | 3 | 3 | 0.187 | 66 |
| SOT17 | | | | | | | | |
| WT vs. M1 | 13.05 | 2.212 | 10.84 | 7.76 | 3 | 3 | 1.976 | 66 |
| WT vs. M2 | 13.05 | 1.514 | 11.54 | 7.76 | 3 | 3 | 2.103 | 66 |
| M1 vs. M2 | 2.212 | 1.514 | 0.6987 | 7.76 | 3 | 3 | 0.1273 | 66 |
| SOT18 | | | | | | | | |
| WT vs. M1 | 12.64 | 1.634 | 11.01 | 7.76 | 3 | 3 | 2.006 | 66 |
| WT vs. M2 | 12.64 | 2.222 | 10.42 | 7.76 | 3 | 3 | 1.899 | 66 |
| M1 vs. M2 | 1.634 | 2.222 | -0.5879 | 7.76 | 3 | 3 | 0.1071 | 66 |

**Supplementary Table 12**

All statistical analyses relating to floret material as performed in Graphpad Prism v8.2.0. M1 = *bolC2.myb28-1$^{ge}$*; M2 = *bolC2.myb28-2$^{ge}$*

| | | | | | | | | |
|---|---|---|---|---|---|---|---|---|
| Floret Glucoraphanin | | | | | | | | |
| Tukey's multiple comparisons test | Mean Diff. | 95.00% CI of diff. | Significant? | Summary | Adjusted P Value | | | |
| WT vs. M1 | 12.26 | 4.836 to 19.69 | Yes | ** | 0.0025 | B-C | | |
| WT vs. M2 | 12.61 | 5.186 to 20.04 | Yes | ** | 0.0021 | B-D | | |
| M1 vs. M2 | 0.3506 | -6.652 to 7.353 | No | ns | 0.99 | C-D | | |
| Test details | Mean 1 | Mean 2 | Mean Diff. | SE of diff. | n1 | n2 | q | DF |
| WT vs. M1 | 12.99 | 0.7234 | 12.26 | 2.75 | 4 | 5 | 6.306 | 11 |
| WT vs. M2 | 12.99 | 0.3728 | 12.61 | 2.75 | 4 | 5 | 6.487 | 11 |
| M1 vs. M2 | 0.7234 | 0.3728 | 0.3506 | 2.593 | 5 | 5 | 0.1912 | 11 |
| Floret Glucoiberin | | | | | | | | |
| Tukey's multiple comparisons test | Mean Diff. | 95.00% CI of diff. | Significant? | Summary | Adjusted P Value | | | |
| WT vs. M1 | 1.64 | 0.1992 to 3.081 | Yes | * | 0.0264 | B-C | | |
| WT vs. M2 | 1.422 | -0.01925 to 2.863 | No | ns | 0.0532 | B-D | | |
| M1 vs. M2 | -0.2184 | -1.577 to 1.140 | No | ns | 0.9022 | C-D | | |
| Test details | Mean 1 | Mean 2 | Mean Diff. | SE of diff. | n1 | n2 | q | DF |
| WT vs. M1 | 2.056 | 0.4152 | 1.64 | 0.5336 | 4 | 5 | 4.347 | 11 |
| WT vs. M2 | 2.056 | 0.6336 | 1.422 | 0.5336 | 4 | 5 | 3.769 | 11 |
| M1 vs. M2 | 0.4152 | 0.6336 | -0.2184 | 0.5031 | 5 | 5 | 0.614 | 11 |
| Floret Hydroxyindolylmethyl | | | | | | | | |
| Tukey's multiple comparisons test | Mean Diff. | 95.00% CI of diff. | Significant? | Summary | Adjusted P Value | | | |
| WT vs. M1 | -0.01175 | -0.1960 to 0.1725 | No | ns | 0.9838 | B-C | | |
| WT vs. M2 | 0.05765 | -0.1266 to 0.2419 | No | ns | 0.684 | B-D | | |
| M1 vs. M2 | 0.0694 | -0.1043 to 0.2431 | No | ns | 0.5456 | C-D | | |

| | | | | | | | | |
|---|---|---|---|---|---|---|---|---|
| Test details | Mean 1 | Mean 2 | Mean Diff. | SE of diff. | n1 | n2 | q | DF |
| WT vs. M1 | 0.1463 | 0.158 | -0.01175 | 0.06821 | 4 | 5 | 0.2436 | 11 |
| WT vs. M2 | 0.1463 | 0.0886 | 0.05765 | 0.06821 | 4 | 5 | 1.195 | 11 |
| M1 vs. M2 | 0.158 | 0.0886 | 0.0694 | 0.0643 | 5 | 5 | 1.526 | 11 |
| Floret Indolylmethyl | | | | | | | | |
| Tukey's multiple comparisons test | Mean Diff. | 95.00% CI of diff. | Significant? | Summary | Adjusted P Value | | | |
| WT vs. M1 | 0.726 | -4.059 to 5.511 | No | ns | 0.9123 | B-C | | |
| WT vs. M2 | 2.127 | -2.658 to 6.912 | No | ns | 0.4771 | B-D | | |
| M1 vs. M2 | 1.401 | -3.111 to 5.913 | No | ns | 0.6879 | C-D | | |
| Test details | Mean 1 | Mean 2 | Mean Diff. | SE of diff. | n1 | n2 | q | DF |
| WT vs. M1 | 4.982 | 4.256 | 0.726 | 1.772 | 4 | 5 | 0.5795 | 11 |
| WT vs. M2 | 4.982 | 2.855 | 2.127 | 1.772 | 4 | 5 | 1.698 | 11 |
| M1 vs. M2 | 4.256 | 2.855 | 1.401 | 1.67 | 5 | 5 | 1.186 | 11 |
| Floret 1-methoxyindolylmethl | | | | | | | | |
| Tukey's multiple comparisons test | Mean Diff. | 95.00% CI of diff. | Significant? | Summary | Adjusted P Value | | | |
| WT vs. M1 | 0.0216 | -0.09150 to 0.1347 | No | ns | 0.8653 | B-C | | |
| WT vs. M2 | 0.0336 | -0.07950 to 0.1467 | No | ns | 0.7093 | B-D | | |
| M1 vs. M2 | 0.012 | -0.09463 to 0.1186 | No | ns | 0.9506 | C-D | | |
| Test details | Mean 1 | Mean 2 | Mean Diff. | SE of diff. | n1 | n2 | q | DF |
| WT vs. M1 | 0.12 | 0.0984 | 0.0216 | 0.04188 | 4 | 5 | 0.7295 | 11 |
| WT vs. M2 | 0.12 | 0.0864 | 0.0336 | 0.04188 | 4 | 5 | 1.135 | 11 |
| M1 vs. M2 | 0.0984 | 0.0864 | 0.012 | 0.03948 | 5 | 5 | 0.4298 | 11 |
| Floret 4-methoxyindolylmethyl | | | | | | | | |
| Tukey's multiple comparisons test | Mean Diff. | 95.00% CI of diff. | Significant? | Summary | Adjusted P Value | | | |
| WT vs. M1 | 0.5259 | -1.716 to 2.768 | No | ns | 0.8051 | B-C | | |

| | | | | | | | | |
|---|---|---|---|---|---|---|---|---|
| WT vs. M2 | 1.201 | -1.041 to 3.442 | No | ns | 0.352 4 | B-D | | |
| M1 vs. M2 | 0.674 6 | -1.439 to 2.788 | No | ns | 0.674 | C-D | | |
| Test details | Mean 1 | Mean 2 | Mean Diff. | SE of diff. | n1 | n2 | q | DF |
| WT vs. M1 | 2.41 | 1.884 | 0.5259 | 0.83 | 4 | 5 | 0.896 | 11 |
| WT vs. M2 | 2.41 | 1.209 | 1.201 | 0.83 | 4 | 5 | 2.045 | 11 |
| M1 vs. M2 | 1.884 | 1.209 | 0.6746 | 0.7826 | 5 | 5 | 1.219 | 11 |
| Floret Sulphate | | | | | | | | |
| Tukey's multiple comparisons test | Mean Diff. | 95.00% CI of diff. | Significa nt? | Summa ry | Adjusted P Value | | | |
| WT vs. M1 | 101.2 | -52.70 to 255.0 | No | ns | 0.226 2 | B-C | | |
| WT vs. M2 | 114.4 | -33.65 to 262.5 | No | ns | 0.14 | B-D | | |
| M1 vs. M2 | 13.24 | -125.7 to 152.1 | No | ns | 0.965 1 | C-D | | |
| Test details | Mean 1 | Mean 2 | Mean Diff. | SE of diff. | n1 | n2 | q | DF |
| WT vs. M1 | 253.1 | 152 | 101.2 | 57.68 | 4 | 5 | 2.481 | 12 |
| WT vs. M2 | 253.1 | 138.7 | 114.4 | 55.5 | 4 | 6 | 2.916 | 12 |
| M1 vs. M2 | 152 | 138.7 | 13.24 | 52.06 | 5 | 6 | 0.3597 | 12 |
| Floret SMCSO | | | | | | | | |
| Tukey's multiple comparisons test | Mean Diff. | 95.00% CI of diff. | Significa nt? | Summa ry | Adjusted P Value | | | |
| WT vs. M1 | 30.54 | -15.88 to 76.96 | No | ns | 0.222 | B-C | | |
| WT vs. M2 | 24.53 | -21.89 to 70.95 | No | ns | 0.361 3 | B-D | | |
| M1 vs. M2 | -6.014 | -49.78 to 37.75 | No | ns | 0.927 4 | C-D | | |
| Test details | Mean 1 | Mean 2 | Mean Diff. | SE of diff. | n1 | n2 | q | DF |
| WT vs. M1 | 88.08 | 57.54 | 30.54 | 17.19 | 4 | 5 | 2.513 | 11 |
| WT vs. M2 | 88.08 | 63.55 | 24.53 | 17.19 | 4 | 5 | 2.018 | 11 |
| M1 vs. M2 | 57.54 | 63.55 | -6.014 | 16.2 | 5 | 5 | 0.5249 | 11 |
| Floret Gene Expression | | | | | | | | |
| Tukey's multiple comparisons test | Mean Diff. | 95.00% CI of diff. | Significa nt? | Summa ry | Adjusted P Value | | | |
| BCAT4 C5 | | | | | | | | |

| | | | | | |
|---|---|---|---|---|---|
| WT vs. M1 | 35.26 | 26.65 to 43.87 | Yes | **** | <0.0001 |
| WT vs. M2 | 36.77 | 28.16 to 45.38 | Yes | **** | <0.0001 |
| M1 vs. M2 | 1.505 | -7.104 to 10.11 | No | ns | 0.9079 |
| BAT5 C3 | | | | | |
| WT vs. M1 | 5.218 | -3.391 to 13.83 | No | ns | 0.32 |
| WT vs. M2 | 5.243 | -3.366 to 13.85 | No | ns | 0.3166 |
| M1 vs. M2 | 0.02484 | -8.584 to 8.634 | No | ns | >0.9999 |
| MAM3 C2 | | | | | |
| WT vs. M1 | 24.32 | 15.72 to 32.93 | Yes | **** | <0.0001 |
| WT vs. M2 | 24.2 | 15.59 to 32.81 | Yes | **** | <0.0001 |
| M1 vs. M2 | -0.1248 | -8.734 to 8.484 | No | ns | 0.9993 |
| IPMI | | | | | |
| WT vs. M1 | 50.42 | 41.81 to 59.03 | Yes | **** | <0.0001 |
| WT vs. M2 | 48.18 | 39.57 to 56.79 | Yes | **** | <0.0001 |
| M1 vs. M2 | -2.246 | -10.85 to 6.363 | No | ns | 0.8067 |
| IMDH3 | | | | | |
| WT vs. M1 | 15.6 | 6.994 to 24.21 | Yes | *** | 0.0001 |
| WT vs. M2 | 15.78 | 7.167 to 24.39 | Yes | *** | 0.0001 |
| M1 vs. M2 | 0.1738 | -8.435 to 8.783 | No | ns | 0.9987 |
| CYP79F1 | | | | | |
| WT vs. M1 | 13.28 | 4.674 to 21.89 | Yes | ** | 0.0013 |
| WT vs. M2 | 13.31 | 4.697 to 21.91 | Yes | ** | 0.0012 |
| M1 vs. M2 | 0.02254 | -8.586 to 8.631 | No | ns | >0.9999 |
| CYP83A1 | | | | | |

| | | | | | | | | |
|---|---|---|---|---|---|---|---|---|
| WT vs. M1 | 68.44 | 59.83 to 77.04 | Yes | **** | <0.0001 | | | |
| WT vs. M2 | 74.28 | 65.67 to 82.89 | Yes | **** | <0.0001 | | | |
| M1 vs. M2 | 5.841 | -2.768 to 14.45 | No | ns | 0.2416 | | | |
| SUR1 | | | | | | | | |
| WT vs. M1 | 38.73 | 30.12 to 47.33 | Yes | **** | <0.0001 | | | |
| WT vs. M2 | 39.84 | 31.24 to 48.45 | Yes | **** | <0.0001 | | | |
| M1 vs. M2 | 1.119 | -7.490 to 9.728 | No | ns | 0.9479 | | | |
| UGT74C1 | | | | | | | | |
| WT vs. M1 | 15.93 | 7.323 to 24.54 | Yes | *** | 0.0001 | | | |
| WT vs. M2 | 16.71 | 8.106 to 25.32 | Yes | **** | <0.0001 | | | |
| M1 vs. M2 | 0.7825 | -7.826 to 9.391 | No | ns | 0.9742 | | | |
| SOT17 | | | | | | | | |
| WT vs. M1 | 9.076 | 0.4673 to 17.68 | Yes | * | 0.0365 | | | |
| WT vs. M2 | 11.77 | 3.163 to 20.38 | Yes | ** | 0.0047 | | | |
| M1 vs. M2 | 2.696 | -5.913 to 11.30 | No | ns | 0.7341 | | | |
| SOT18 | | | | | | | | |
| WT vs. M1 | 6.884 | -1.724 to 15.49 | No | ns | 0.1418 | | | |
| WT vs. M2 | 6.72 | -1.889 to 15.33 | No | ns | 0.155 | | | |
| M1 vs. M2 | -0.1646 | -8.773 to 8.444 | No | ns | 0.9988 | | | |
| Test details | Mean 1 | Mean 2 | Mean Diff. | SE of diff. | N1 | N2 | q | DF |
| BCAT4 C5 | | | | | | | | |
| WT vs. M1 | 39.72 | 4.458 | 35.26 | 3.59 | 3 | 3 | 13.89 | 66 |
| WT vs. M2 | 39.72 | 2.954 | 36.77 | 3.59 | 3 | 3 | 14.48 | 66 |
| M1 vs. M2 | 4.458 | 2.954 | 1.505 | 3.59 | 3 | 3 | 0.5927 | 66 |

| | | | | | | | | |
|---|---|---|---|---|---|---|---|---|
| BAT5 C3 | | | | | | | | |
| WT vs. M1 | 5.551 | 0.3336 | 5.218 | 3.59 | 3 | 3 | 2.055 | 66 |
| WT vs. M2 | 5.551 | 0.3088 | 5.243 | 3.59 | 3 | 3 | 2.065 | 66 |
| M1 vs. M2 | 0.3336 | 0.3088 | 0.02484 | 3.59 | 3 | 3 | 0.009783 | 66 |
| MAM3 C2 | | | | | | | | |
| WT vs. M1 | 25.12 | 0.8008 | 24.32 | 3.59 | 3 | 3 | 9.581 | 66 |
| WT vs. M2 | 25.12 | 0.9257 | 24.2 | 3.59 | 3 | 3 | 9.532 | 66 |
| M1 vs. M2 | 0.8008 | 0.9257 | -0.1248 | 3.59 | 3 | 3 | 0.04917 | 66 |
| IPMI | | | | | | | | |
| WT vs. M1 | 53.35 | 2.925 | 50.42 | 3.59 | 3 | 3 | 19.86 | 66 |
| WT vs. M2 | 53.35 | 5.171 | 48.18 | 3.59 | 3 | 3 | 18.98 | 66 |
| M1 vs. M2 | 2.925 | 5.171 | -2.246 | 3.59 | 3 | 3 | 0.8846 | 66 |
| IMDH3 | | | | | | | | |
| WT vs. M1 | 16.95 | 1.344 | 15.6 | 3.59 | 3 | 3 | 6.145 | 66 |
| WT vs. M2 | 16.95 | 1.17 | 15.78 | 3.59 | 3 | 3 | 6.214 | 66 |
| M1 vs. M2 | 1.344 | 1.17 | 0.1738 | 3.59 | 3 | 3 | 0.06844 | 66 |
| CYP79F1 | | | | | | | | |
| WT vs. M1 | 13.78 | 0.4966 | 13.28 | 3.59 | 3 | 3 | 5.232 | 66 |
| WT vs. M2 | 13.78 | 0.4741 | 13.31 | 3.59 | 3 | 3 | 5.241 | 66 |
| M1 vs. M2 | 0.4966 | 0.4741 | 0.02254 | 3.59 | 3 | 3 | 0.008878 | 66 |
| CYP83A1 | | | | | | | | |
| WT vs. M1 | 83.37 | 14.93 | 68.44 | 3.59 | 3 | 3 | 26.96 | 66 |
| WT vs. M2 | 83.37 | 9.089 | 74.28 | 3.59 | 3 | 3 | 29.26 | 66 |
| M1 vs. M2 | 14.93 | 9.089 | 5.841 | 3.59 | 3 | 3 | 2.301 | 66 |
| SUR1 | | | | | | | | |
| WT vs. M1 | 48.98 | 10.25 | 38.73 | 3.59 | 3 | 3 | 15.25 | 66 |
| WT vs. M2 | 48.98 | 9.134 | 39.84 | 3.59 | 3 | 3 | 15.69 | 66 |
| M1 vs. M2 | 10.25 | 9.134 | 1.119 | 3.59 | 3 | 3 | 0.4408 | 66 |
| UGT74C1 | | | | | | | | |
| WT vs. M1 | 26.98 | 11.05 | 15.93 | 3.59 | 3 | 3 | 6.275 | 66 |
| WT vs. M2 | 26.98 | 10.27 | 16.71 | 3.59 | 3 | 3 | 6.584 | 66 |
| M1 vs. M2 | 11.05 | 10.27 | 0.7825 | 3.59 | 3 | 3 | 0.3082 | 66 |

| | | | | | | | | |
|---|---|---|---|---|---|---|---|---|
| SOT17 | | | | | | | | |
| WT vs. M1 | 14.9 | 5.826 | 9.076 | 3.59 | 3 | 3 | 3.575 | 66 |
| WT vs. M2 | 14.9 | 3.13 | 11.77 | 3.59 | 3 | 3 | 4.637 | 66 |
| M1 vs. M2 | 5.826 | 3.13 | 2.696 | 3.59 | 3 | 3 | 1.062 | 66 |
| SOT18 | | | | | | | | |
| WT vs. M1 | 8.541 | 1.657 | 6.884 | 3.59 | 3 | 3 | 2.712 | 66 |
| WT vs. M2 | 8.541 | 1.822 | 6.72 | 3.59 | 3 | 3 | 2.647 | 66 |
| M1 vs. M2 | 1.657 | 1.822 | -0.1646 | 3.59 | 3 | 3 | 0.06485 | 66 |
